# Discovery and Development of First-in-Class Cereblon-Recruiting RIPK1 Degraders

**DOI:** 10.64898/2026.04.10.717852

**Authors:** Dong Lu, Xin Yu, Hanfeng Lin, Ran Cheng, Min Zhang, Bin Yang, Jingjing Chen, Feng Li, Xiaoli Qi, Jin Wang

**Affiliations:** Verna and Marrs McLean Department of Biochemistry and Molecular Pharmacology, Baylor College of Medicine, Houston, TX 77030; Center for NextGen Therapeutics, Baylor College of Medicine, Houston, TX 77030; Department of Molecular and Cellular Biology, Baylor College of Medicine, Houston, TX 77030; Department of Pathology, Baylor College of Medicine, Houston, TX 77030; Center for Drug Discovery, Baylor College of Medicine, Houston, TX 77030

**Keywords:** RIPK1, PROTAC, Cereblon, Optimization, First-in-class

## Abstract

Receptor-interacting protein kinase 1 (RIPK1) is a critical regulator of programmed cell death and is implicated in various pathological conditions, particularly in mediating tumor resistance to immune checkpoint inhibitors (ICBs). In this study, we have pioneered the development of a novel cereblon (CRBN)-recruiting RIPK1 degrader, **LD5095**, through systematic optimization of linker and CRBN ligand portion. **LD5095** demonstrates potent and selective RIPK1 degradation across cell lines, with rapid kinetics and sustained degradation over 72h post-washout. Functionally, RIPK1 degradation by **LD5095** significantly sensitized Jurkat cells to TNF_α_-induced apoptosis. Furthermore, **LD5095** exhibited favorable pharmacokinetics, including metabolic stability and an extended half-life. Strikingly, *in vivo*, a single dose of **LD5095** achieved durable RIPK1 degradation in xenograft tumors over 6 days. These findings underscore the potential of **LD5095** as a chemical probe for studying RIPK1 biology and a promising candidate for cancer treatment.

## INTRODUCTION

Receptor-interacting protein kinase 1 (RIPK1) serves as a pivotal regulator in two major forms of programmed cell death, namely apoptosis and necroptosis.^1–4^ Its involvement spans across a wide spectrum of pathological conditions, including ischemia-reperfusion injury^5^, neurodegenerative disorders^3,6^, inflammatory diseases^7,8^, infection ailments^9,10^, and tumor metastasis.^11–14^

Recent studies have revealed that the scaffolding function of RIPK1 regulates IFN-γ-driven immune checkpoint blockade (ICB) resistance.^15^ Deletion of tumor RIPK1 leads to beneficial alterations in the tumor microenvironment, such as an increased frequency of effector T cells, a decreased frequency of immuno-suppressive myeloid cells, and changes in inflammatory cytokine secretion. These alterations can influence immune infiltrate dynamics, resulting in heightened tumor sensitivity to anti-PD1 therapy.^15–17^ Conversely, RIPK1 kinase inhibitors have demonstrated limited efficacy in synergizing with anti-PD1 treatments to bolster antitumor immunity.^15,17,18^ Hence, pharmacological degradation of RIPK1 emerges as a potentially safe and effective therapeutic approach, with promising prospects for synergizing with immune checkpoint inhibitors (ICBs) to enhance antitumor immunity. Additionally, given the lethality associated with RIPK1 knockout mice, RIPK1 degraders offer a valuable tool for studying the biological role of RIPK1 *in vivo*, encompassing adult development, infectious diseases, and beyond.^19^

So far, several RIPK1 degraders have been developed (Figure 1).^18,20–23^ These degraders demonstrated potent and highly specific induction of RIPK1 degradation across diverse cancer cell lines.^18,24,25^ Importantly, our findings, as well as those from other labs, highlighted significant synergies between RIPK1 degraders and either anti-PD1 immunotherapy or radiotherapy.^18,20,22^ This synergy resulted in a remarkable delay in tumor development, underscoring the promising potential of RIPK1 degraders in cancer treatment strategies.

**Figure 1.**
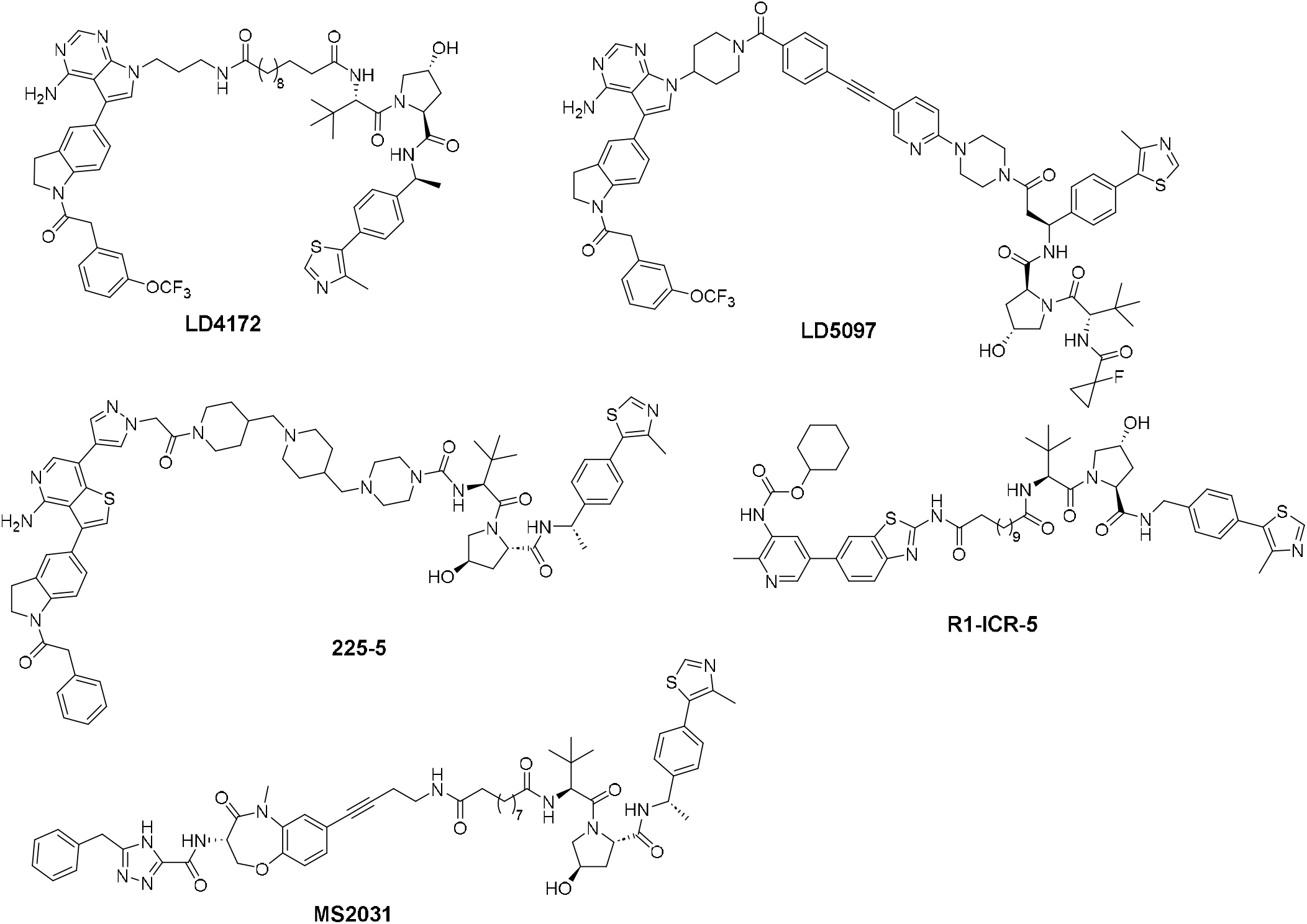
The chemical structures of reported RIPK1 degraders.

Despite the encouraging preclinical outcomes observed with recent RIPK1 degraders, their suboptimal pharmacokinetic characteristics, notably high *in vivo* clearance and low plasma drug concentration^18,20^, pose significant hurdles for their translation into clinical applications. The incorporation of a VHL ligand as the E3 ligase component in these degraders may exacerbate these pharmacokinetic limitations. In contrast, ligands for Cereblon (CRBN), such as thalidomide, have been extensively utilized in PROTACs as E3 ligase ligands.^26–32^ Given their smaller molecular weight and optimal physicochemical properties, CRBN-based PROTACs generally exhibit favorable pharmacokinetic profiles. Notably, at least 24 CRBN-based PROTACs in clinical trials have demonstrated oral availability.^33,34^ Crucially, to date, there have been no reports of a fully characterized RIPK1 PROTAC utilizing the CRBN recruitment strategy in the public domain. Therefore, switching to a CRBN-recruiting strategy represents a compelling, rational design approach to overcome the PK liabilities of prior RIPK1 degraders and accelerate their clinical potential.

Herein, we present our design, synthesis, and evaluation of RIPK1 degraders utilizing the CRBN recruitment strategy. Given the availability of numerous CRBN ligands, we systematically assessed several candidates to inform the design of our RIPK1 degraders. Additionally, recognizing the critical role of the linker in PROTACs for degradation potency and stability, we conducted thorough optimization of this component. Our efforts culminated in the identification of a highly potent RIPK1 degrader, designated as compound **LD5095** . Compound **LD5095** exhibited a DC_50_ value of 1.4 nM and D_max_ > 90% for endogenous RIPK1 in the Jurkat cancer cell line. Furthermore, it demonstrated a markedly prolonged plasma half-life (T_1/2_ = 21.2 h) and acceptable plasma concentration (AUC_0∼last_ = 2903 ng*h/mL). Notably, **LD5095** displayed robust and persistent RIPK1 degradation *in vivo*, positioning it as a valuable tool for RIPK1 biology study as well as a promising candidate for further advancement in cancer therapies.

## RESULTS

### Chemistry

The synthetic route for the preparation of compounds **3** is outlined in Scheme 1. The process commences with the nucleophilic aromatic substitution reaction of the starting material 2-(2,6-dioxopiperidin-3-yl)-4-fluoroisoindoline-1,3-dione with diverse aliphatic amines, followed by deprotection to yield intermediate **1**. Subsequently, the amide coupling reaction of intermediate **1** with compound **2** furnishes compound **3**.

**Scheme 1.**
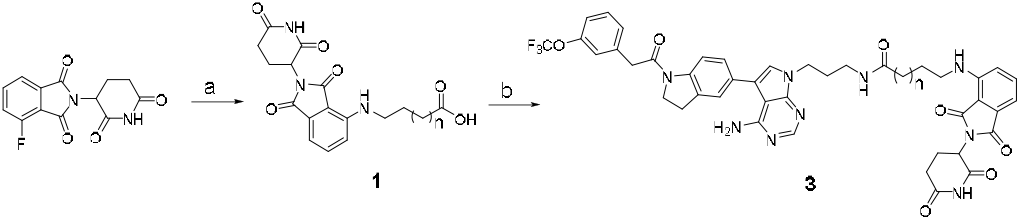
Synthesis Route for compounds 3a-d^a^. ^a^Reagents and conditions: (a) (i) DIPEA, DMF, 90°C; (ii) TFA, DCM, rt; (b) **2**, HATU, DMF, rt.

Scheme 2 illustrates the synthetic pathway for the preparation of compound **5**. Initially, the amide coupling reaction of compound **2** with tert-butyl (10-aminodecyl)carbamate was performed, followed by the removal of the Boc-protecting group under acidic conditions, yielding intermediate **4**. Additionally, various CRBN binders were either commercially available or synthesized following reported procedures.^35–38^ These binders were subsequently coupled with compound **4** to afford compound **5**.

**Scheme 2.**
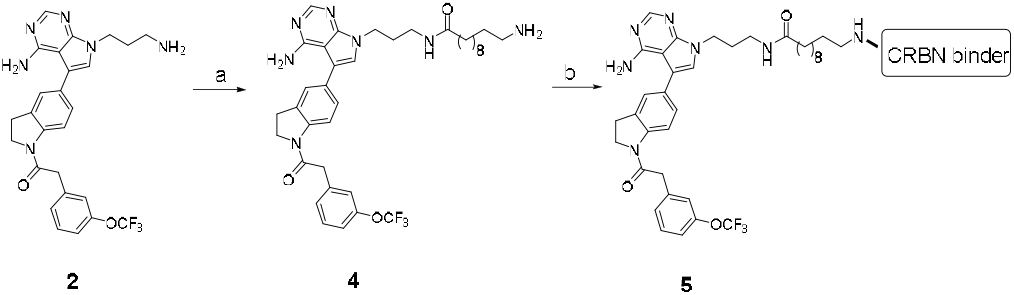
Synthesis Route of compound 5a-f^a^. ^a^Reagents and conditions: (a) (i) HATU, DIPEA, DMF, rt; (ii) TFA, DCM, rt; (b) HATU, DIPEA, DMF, rt.

As shown in Scheme 3, the Sonogashira coupling reaction of ethyl 4-iodobenzoate, followed by the removal of the Boc-protecting group under acidic conditions, afforded the key intermediate **6. 7** was synthesized via Borch reductive amination of compound **6**, followed by hydrolysis. Starting from 5-bromo-2-fluoropyridine, nucleophilic substitution with tert-butyl 2,6-diazaspiro[3.3]heptane-2-carboxylate, followed by deprotection, yielded intermediate **8**. Subsequently, the Sonogashira coupling reaction of **8**, tert-butyl 4-(5-bromopyridin-2-yl)piperazine-1-carboxylate or tert-butyl (6-bromopyridin-3-yl)carbamate, followed by hydrolysis, yielded compounds **9-11**, respectively. Intermediate **12** was obtained through the nucleophilic aromatic substitution reaction of 5-bromo-2-fluoropyrimidine, which was then transformed into the target compound **13** via Suzuki coupling.

**Scheme 3.**
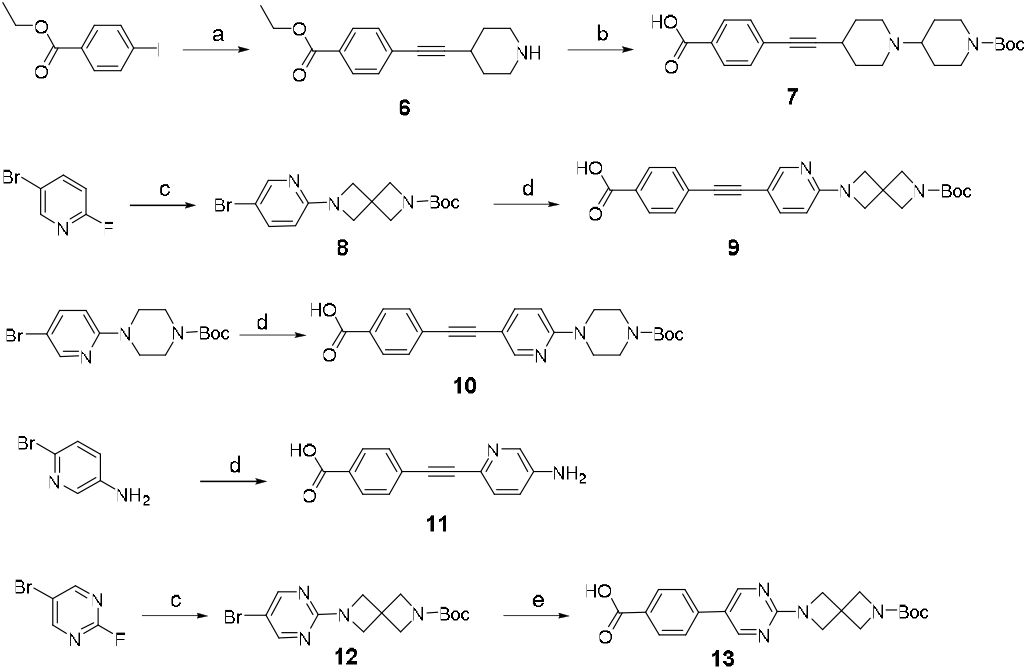
Synthesis Routes of linkers 7, 9-11 and 13^a^. ^a^Reagents and conditions: (a) (i) Pd(PPh_3_)_2_Cl_2_, CuI, TEA, DMF,100°C; (ii) TFA, DCM, rt; (b) NaBH(OAc)_3_, AcOH, DCE, rt; (ii) LiOH, THF/H_2_O = 5/1, rt; (c) tert-butyl 2,6-diazaspiro[3.3]heptane-2-carboxylate, TEA, DMF,100°C; (d) (i) methyl 4-ethynylbenzoate, Pd(PPh_3_)_2_Cl_2_, CuI, TEA, DMF,100°C; (ii) LiOH, THF/H_2_O = 5/1, rt; (e) (i) Pd(dppf)_2_Cl_2_, K_2_CO_3_, dioxane/H_2_O = 10/1,100°C; (ii) LiOH, THF/H_2_O = 5/1, rt.

As illustrated in Scheme 4, the target compounds **15** and **18** were synthesized by coupling the CRBN ligand with the key intermediates **14** and **17**, respectively, derived from the starting materials **2** or **16** with different linkers, followed by deprotection.

**Scheme 4.**
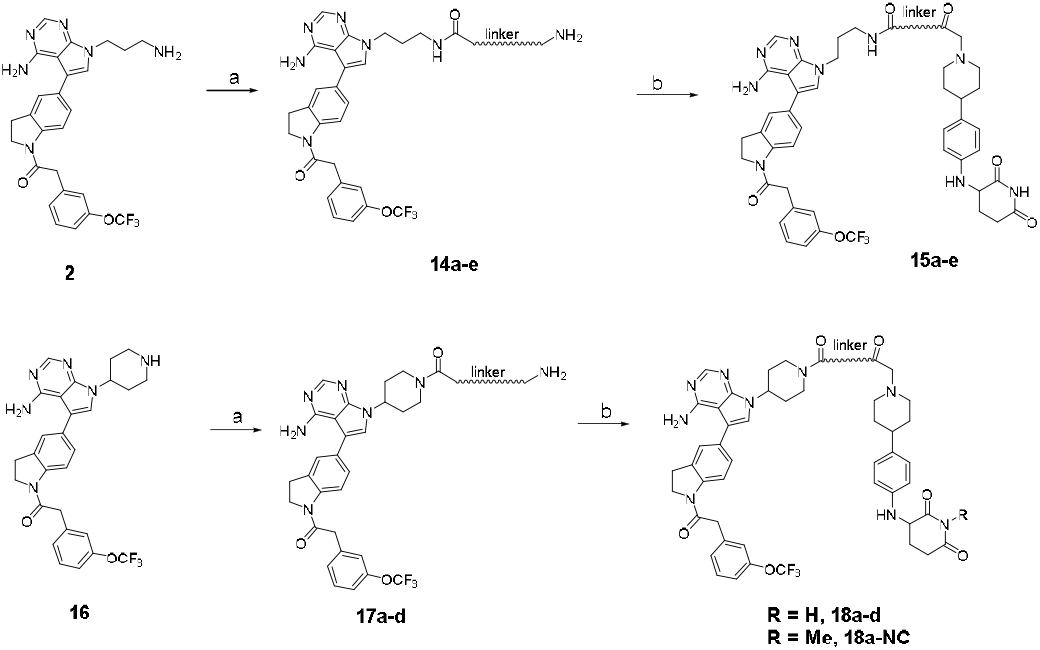
Synthesis Routes of Compounds 15 and 18^a^. ^a^Reagents and conditions: (a) (i) HATU, DIPEA, DMF, rt; (ii) TFA, DCM, rt; (b) HATU, DIPEA, DMF, rt.

### Identification of Pomalidomide-based PROTACs as Potent RIPK1 Degraders

Our investigation began with the design and synthesis of a series of PROTACs utilizing pomalidomide as the CRBN binder and a type II RIPK1 inhibitor as the RIPK1 warhead^39^. Homology modeling suggested an ethyl group orientation conducive to solvent exposure, facilitating an optimal exit vector for PROTAC linkage.^18^ We systematically screened various aliphatic linker lengths ranging from 3 to 10 carbon atoms to maximize the pairing of the RIPK1−PROTAC−CRBN ternary complex (Figure 2a). Evaluation of RIPK1 degradation potency in a stable Jurkat cell line expressing nLuc-RIPK1 revealed significant variability across different linker lengths.

**Figure 2.**
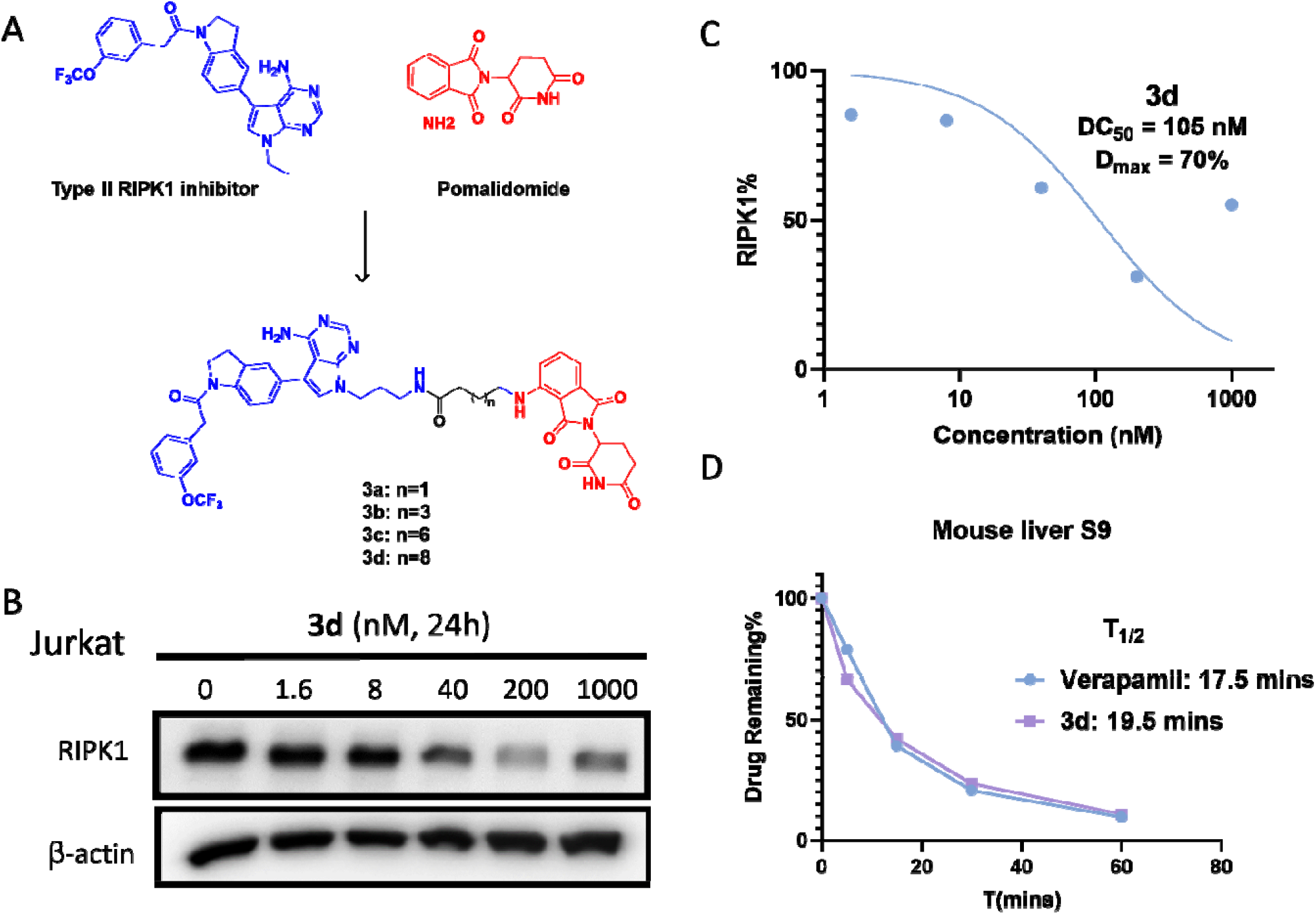
Degradation potency and *in vitro* metabolic stability of pomalidomide-based RIPK1 PROTACs. (A) Design of pomalidomide-based RIPK1 PROTACs. (B) Western blot of endogenous RIPK1 levels of WT Jurkat cells treated with indicated concentration of **3d** for 24h. (C) Dose-response curve of **3d** for RIPK1 degradation in WT Jurkat cells. (D) Metabolic stability of **3d** and Verapamil in mouse liver S9 fraction

As shown in Table 1, compounds **3a** and **3b**, featuring 3 and 5 carbon atom linkers, respectively, exhibited minimal degradation of nLuc-RIPK1 at 1 μM concentration. Compound **3c**, with an 8-carbon linker, displayed moderate degradation activity, reducing 37% of nLuc-RIPK1 at the same concentration. Intriguingly, compound **3d**, with longer linker, demonstrated further improvement in degradation potency, resulting in reduction of 73% of nLuc-RIPK1. Further evaluation of **3d** for degrading endogenous RIPK1 in WT Jurkat cells via Western blot analysis revealed a significant reduction in RIPK1 levels at 200 nM concentration with concentration for 50% degradation (DC_50_) and maximum degradation (D_max_) values of 105 nM and 70%, respectively (Figure 2b-c). However, a diminished degradation effect was observed at 1 μM concentrations (Figure 2b), indicative of the “hook effect.” Assessment of the metabolic stability of **3d** in mouse liver S9 fraction in the presence of NADPH showed rapid metabolism, with a half-life (T_1/2_) of 19.5 mins, similar to the positive control, Verapamil (Figure 2d).

**Table 1.** Assessment of Degradation Potency of Pomalidomide-based Degraders with Varied Linker Lengths.

| No. | n | nLuc-RIPK1 level @1 μM<sup>a</sup> |
| --- | --- | --- |
| **3a** | 2 | 96% |
| **3b** | 4 | 98% |
| **3c** | 7 | 63% |
| **3d** | 9 | 27% |

<sup>a</sup>nLuc-RIPK1 level in Jurkat cells. Data are the mean of triplicate determinations.

### Optimization of CRBN Ligand

Significant efforts have been devoted to the exploration of new CRBN ligands, aiming to identify both “thalidomide-like” molecules and “non-thalidomide-like” chemotypes.^27^ In pursuit of developing CRBN-based RIPK1 degraders with enhanced degradation potency and metabolic stability, we utilized compound **3d** as a template RIPK1 degrader and synthesized a series of new RIPK1 degraders employing reported CRBN ligands.

As shown in Table 2 and Figure S1, evaluation of these compounds revealed notable differences in RIPK1 degradation activity. Compound **3d** exhibited potent RIPK1 degradation, with a DC_50_ value of 14.0 nM and achieving a D_max_ of 75% RIPK1 in nLuc-RIPK1 Jurkat cells. Similarly, compound **5a**, containing the thalidomide moiety but differing in linkage attachment site, demonstrated robust RIPK1 degradation activity, with a DC_50_ value of 13.8 nM and achieving a D_max_ of 72% RIPK1. In contrast, replacing the thalidomide moiety with a thalidomide-like CRBN ligand (N-linked degron)^36^ yielded compound **5b**, which exhibited superior degradation activity for nLuc-RIPK1, with a DC_50_ value of 6.3 nM and a D_max_ of 90%. Conversely, compounds **5c-f**, incorporating other CRBN binders, showed diminished degradation activities, achieving maximum degradation levels of only 30-39% of RIPK1. Collectively, compound **5b**, featuring the N-linked degron, demonstrated the most potent degradation activit**y** and warrants further optimization.

**Table 2.**
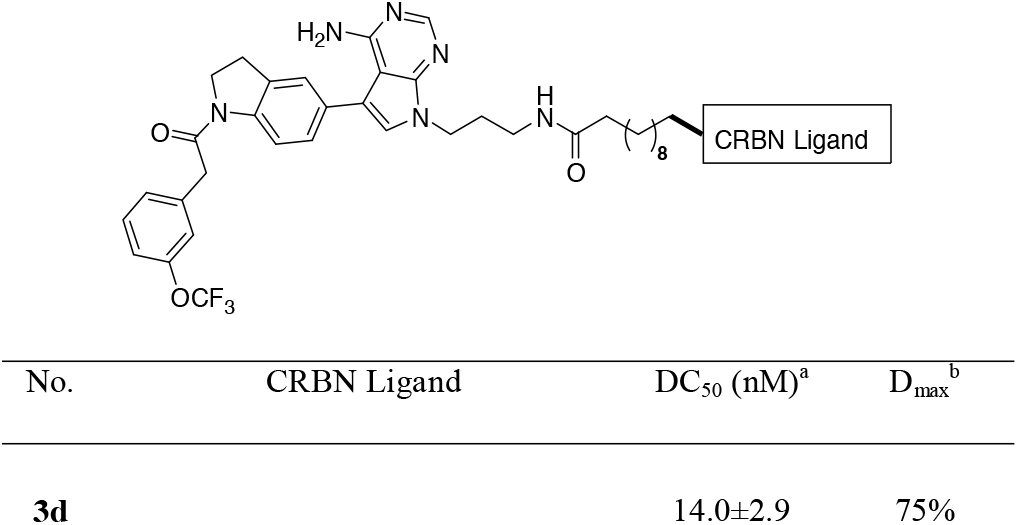
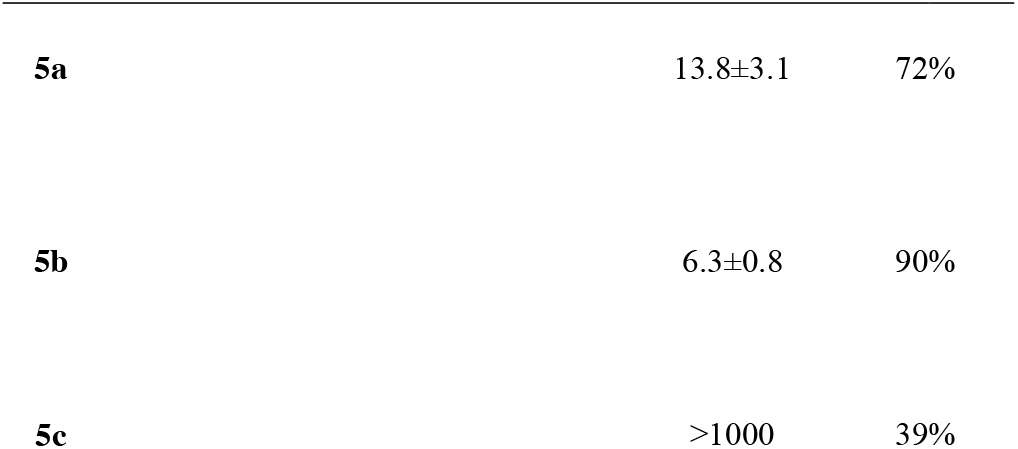
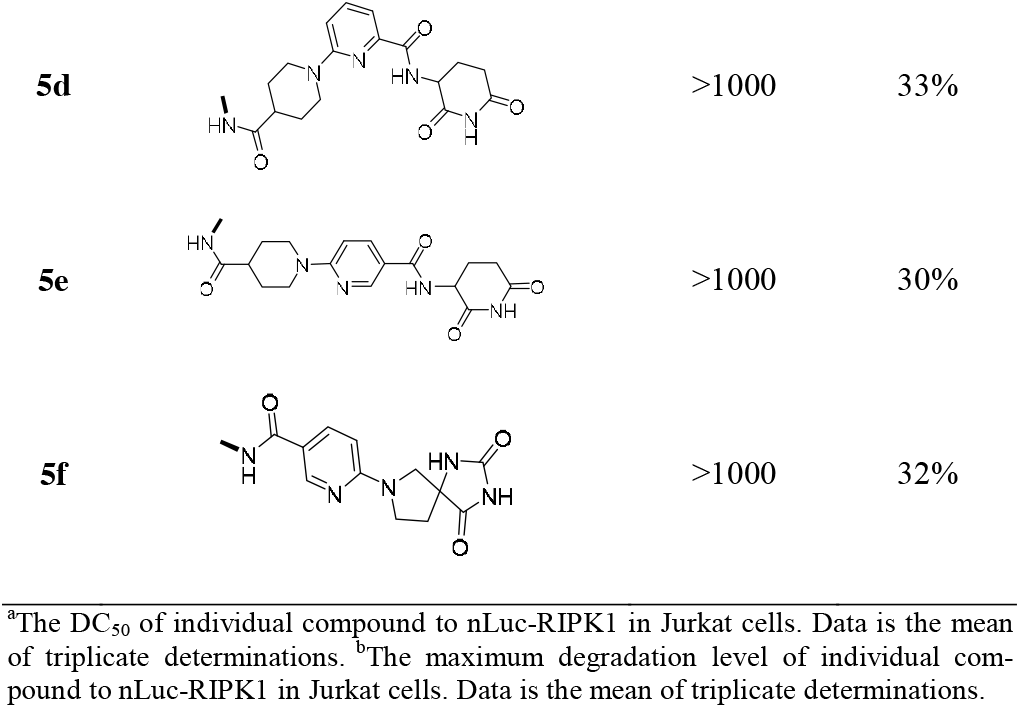
Exploration of CRBN Ligand in RIPK1 Degraders.

### Optimization of the Linker in 5b

The impact of linker composition and rigidity on the potency and pharmacokinetic (PK) properties of PROTACs has been well recognized.^40–44^ Therefore, we systematically investigated the influence of linker rigidity on both the potency of RIPK1 degradation and the metabolic stability in mouse liver S9 fraction.

As depicted in Table 3 and Figure S1, the replacement of aliphatic chains with rigid linkers (compounds **15a-e**) yielded potent degradation activities, with DC_50_ values ranging from 3.7 to 29.9 nM. Moreover, compounds **15a-e** achieved over 85% RIPK1 degradation in Jurkat cells. Metabolic stability assessment revealed that compounds **15a-b**, incorporating relatively flexible linkers, exhibited half-lives (T_1/2_) of 11.3 and 23.6 mins, respectively, with no improvement compared to **5b**. In contrast, compounds **15c-e**, featuring more rigid linkers, demonstrated prolonged half-lives in liver S9 fraction compared to **5b**. Remarkably, compound **15e** exhibited the highest stability in mouse liver S9 fraction, with a T_1/2_ exceeding 120 mins, underscoring the potential of rigid linkers to enhance the metabolic stability of degraders.

**Table 3.** Impact of Linkers on Degradation Potency and Metabolic Stability.

| No. | Linker | DC<sub>50</sub> (nM)<sup>a</sup> | D<sub>max</sub><sup>b</sup> | T<sub>1/2</sub> (mins)<sup>c</sup> |
| --- | --- | --- | --- | --- |
| **5b** |  | 6.3±0.8 | 90% | 21.5 |
| **15a** |  | 29.9±6.9 | 87% | 11.3 |
| **15b** |  | 10.6±1.2 | 85% | 23.6 |
| **15c** |  | 3.7±0.5 | 93% | 49.3 |
| **15d** |  | 6.5±1.0 | 90% | 97.6 |
| **15e** |  | 6.9±2.2 | 88% | >120 |

<sup>a</sup>The DC<sub>50</sub> of individual compound to nLuc-RIPK1 in Jurkat cells. Data are the mean of triplicate determinations. <sup>b</sup>The maximum degradation level of individual compound to nLuc-RIPK1 in Jurkat cells. Data are the mean of triplicate determinations. <sup>c</sup>The half-life time of individual compound incubated with mouse liver S9 fraction in the presence of NADPH. Data are the mean of duplicate determinations.

### Further Refinement of Exit Vector and Linker in RIPK1 Degraders

To further enhance the stability of degraders, we employed a strategy of incorporating both a cyclic exit vector and a rigid linker into the design of RIPK1 degraders, resulting in a new series of compounds. As depicted in Table 4 and Figure S1, all degraders (**18a-c**) bearing these optimized moieties exhibited highly potent RIPK1 degradation activities, with DC_50_ values of 2.5, 4.8, and 5.3 nM, respectively. These compounds effectively degraded over 80% of RIPK1. However, compound **18d**, featuring a shorter linker, displayed decreased degradation potency, with a DC_50_ value of 122 nM, suggesting that a short linker may impede the formation of the ternary complex.

**Table 4.** Exploration of RIPK1 Degraders with New Exit Vector and Rigid Linker.

| No. | Linker | DC<sub>50</sub> (nM)<sup>a</sup> | D<sub>max</sub><sup>b</sup> | T<sub>1/2</sub> (mins)<sup>c</sup> |
| --- | --- | --- | --- | --- |
| **5b** |  | 6.3±0.8 | 90% | 21.5 |
| **LD5095 (18a)** |  | 2.5±0.5 | 91% | >120 |
| **18b** |  | 4.8±0.6 | 85% | >120 |
| **18c** |  | 9.3±1.4 | 80% | >120 |
| **18d** |  | 122±25 | 62% | NT<sup>d</sup> |

<sup>a</sup>The DC<sub>50</sub> of individual compound to nLuc-RIPK1 in Jurkat cells. Data are the mean of triplicate determinations. <sup>b</sup>The maximum degradation level of individual compound to nLuc-RIPK1 in Jurkat cells. Data are the mean of triplicate determinations. <sup>c</sup>The half-life time of individual compound incubated with mouse liver S9 fraction in the presence of NADPH. Data are the mean of duplicate determinations. <sup>d</sup>Not test.

To assess their potential for enhanced metabolic stability, we evaluated the *in vitro* stability of compounds **18a-c** in mouse liver S9 fraction. As anticipated, all these degraders displayed substantially prolonged half-lives (T_1/2_ > 120 mins) in mouse liver S9 fraction, indicating significant improvement in metabolic stability.

### Evaluation of Pharmacokinetic Profile of CRBN-based RIPK1 Degraders

To comprehensively assess the *in vivo* stability of the optimized compounds, namely **15c, 15d, 15e**, and **18a**, we conducted pharmacokinetic studies in mice (n=3) following intravenous administration at a dose of 1.0 mg/kg. As summarized in Table 5 and Figure S2, compounds 15c-e displayed clearance rates of 15.7, 12.3, and 8.11 mL/min/kg, respectively, along with corresponding area under the curve (AUC_0-last) values of 1091, 1369, and 2114 h·ng/mL. Notably, compound 18a (LD5095) exhibited the most favorable pharmacokinetic properties, featuring a prolonged plasma half-life (t1/2) of 21.2 hours and an AUC_0-last of 2903 h·ng/mL. Given the superior IV profile of 18a, we further evaluated its oral bioavailability in mice (n=3) at a dose of 20 mg/kg (Figure S6, Table S1). Following oral administration, 18a reached a peak plasma concentration (C_max_) of 304 ng/mL at a T_max_ of 2.7 hours, with a terminal half-life of 7.3 hours. The oral exposure, evidenced by an AUC_0-inf of 4795 h·ng/mL, confirms that 18a possesses sufficient drug-like properties and metabolic stability for oral dosing. These results underscore the significantly improved pharmacokinetic profiles of optimized degraders with rigid linkers and cyclic exit vectors, positioning 18a as a promising candidate for further *in vivo* efficacy and pharmacodynamic evaluations.

**Table 5.** Pharmacokinetic Parameters of CRBN Based RIPK1 Degraders in Mice ^a^.

|  | Cl<sub>plasma</sub> (mL/min/kg) | T<sub>1/2</sub> (h) | C<sub>max</sub> (ng/mL) | AUC<sub>0–last</sub> (h·ng/mL) | Vd (L/kg) |
| --- | --- | --- | --- | --- | --- |
| **15c** | 15.7±3.8 | 8.1±0.2 | 907±222 | 1091±253 | 4.5±1.1 |
| **15d** | 12.3±2.2 | 7.0±2.6 | 734±125 | 1369±256 | 4.5±0.4 |
| 15e | 8.1±1.8 | 3.4±0.2 | 1507±35<br>3 | 2114±474 | 1.6±0.3 |
| LD5095 | 4.9±0.6 | 21.2±1.8 | 2007±46<br>2 | 2903±316 | 6.7±0.3 |

<sup>a</sup> Mice were treated with compounds via intravenous administration (1 mg/kg, n=3).

### Evaluation of LD5095 in Various Cancer Cell Lines

Due to its potent degradation activity and superior pharmacokinetic properties, we designated **18a** as **LD5095** for further evaluation as a lead compound. To assess the degradation potency of **LD5095** against endogenous RIPK1 in cells, we conducted a western blot assay to evaluate its RIPK1 degradation activities in WT Jurkat, MOLM14, and U937 cell lines. Treatment with **LD5095** in a dose-dependent manner resulted in the degradation of endogenous RIPK1 after 24 hours in all three cell lines, with DC_50_ values of 1.4, 1.2, and 2.3 nM, respectively. Moreover, **LD5095** achieved maximal RIPK1 degradation at 200 nM, reaching D_max_ values of 94% in Jurkat cells, 95% in MOLM14 cells, and 91% in U937 cells (Figure 3a, S3). In addition, the degradation kinetics of RIPK1 induced by **LD5095** were rapid, with complete degradation observed within 4 hours (Figure 3b).

**Figure 3.**
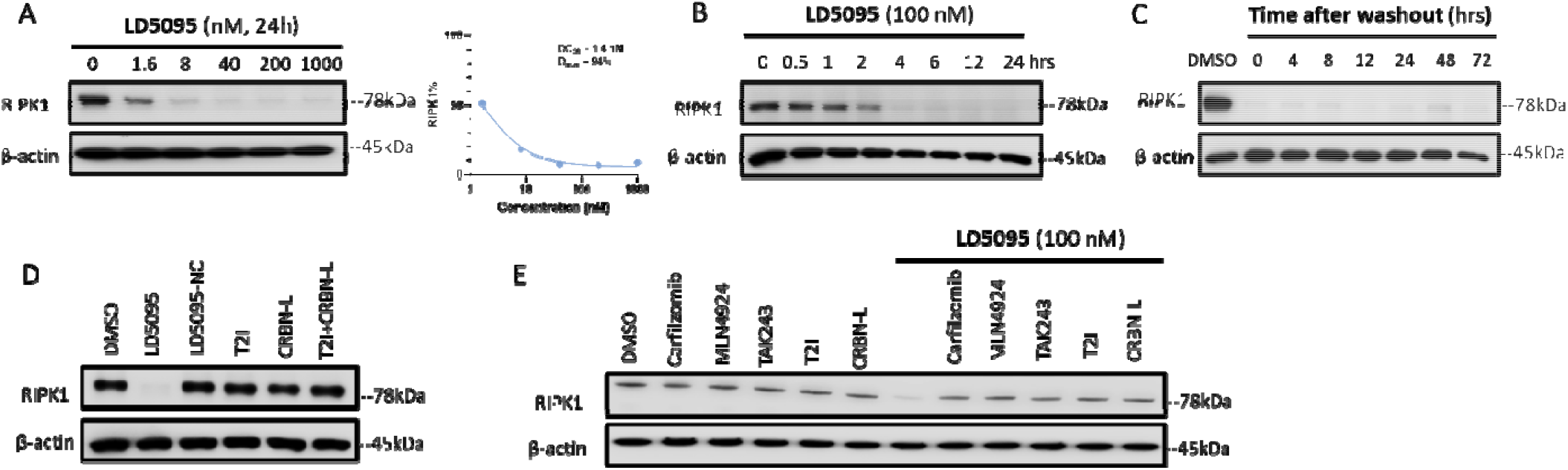
Potency, Kinetics, and Mechanistic Validation of LD5095-Mediated RIPK1 Degradation.(A) Concentration-dependent degradation potency of LD5095 against endogenous RIPK1 in Jurkat cells following 24-hour treatment.(B) Representative Western blot analysis of RIPK1 levels in Jurkat cells treated with 100 nM LD5095 across the indicated time points.(C) Recovery kinetics of RIPK1 levels in Jurkat cells after initial treatment with 100 nM LD5095 for 12 hours, followed by three PBS washes and replenishment with fresh media; cells were collected at the indicated time points.(D) Representative Western blot of RIPK1 levels following 24-hour treatment with LD5095 (100 nM), the negative control LD5095-NC (100 nM), the RIPK1 warhead T2I (100 nM), the CRBN ligand (100 nM), or a combination of T2I and the CRBN ligand. (E) Representative Western blot analysis of RIPK1 levels under competitive or pathway-inhibitory conditions where Jurkat cells were pre-treated for 2 hours with either a proteasome inhibitor (Carfilzomib, 300 nM), a neddylation inhibitor (MLN4924, 1 μM), a ubiquitin-activating enzyme (UAE) inhibitor (TAK-243, 250 nM), or an excess of either the RIPK1 warhead (T2I, 1 μM) or the CRBN ligand (CRBN-L, 1 μM), and subsequently treated with DMSO or LD5095 (100 nM) for 4 hours prior to collection.

### Mechanism Validation of LD5095 *in vitro*

Treatment with the negative control (LD5095-NC, featuring a methylated glutarimide), the RIPK1 warhead (T2I) alone, the CRBN ligand (CRBN-L) alone, or the combination of T2I and CRBN-L did not impact RIPK1 levels. This confirms that LD5095 functions as a bone fide bifunctional PROTAC, requiring the formation of a stable ternary complex to induce target degradation (Figure 3d). Furthermore, **LD5095**-mediated RIPK1 degradation was effectively abolished when cells were pre-treated with the ubiquitin-activating enzyme (UAE) inhibitor TAK-243, the neddylation inhibitor MLN-4924, or the proteasome inhibitor carfilzomib. The rescue of RIPK1 levels by these agents, along with the inhibition observed upon competition with 10-fold excess T2I or 100-fold excess CRBN-L, provides definitive evidence that **LD5095** operates through a standard PROTAC mechanism of action (Figure 3e). In addition, quantitative PCR (qPCR) analysis revealed that **LD5095** did not alter the mRNA expression of RIPK1 (Figure S7). By blocking the proteasome, a modest increase of ubiquitinated RIPK1 in the carfilzomib-induced strong ubiquitination background was observed with **LD5095** treatment (Figure S8). These results collectively confirm that **LD5095**-induced degradation is strictly dependent on the specific recruitment of the CRBN E3 ligase and the subsequent activation of the entire ubiquitin-proteasome system (UPS) cascade in Jurkat cells.

### LD5095 Exhibits Prolonged Cellular Activity After Washout

To investigate the cellular retention of **LD5095**, we monitored RIPK1 levels in Jurkat cells after extensive washout of **LD5095**. As shown in Figure 3c, even after 48 and 72 hours following the 12-hour treatment and subsequent washout, minimal resynthesis of RIPK1 protein was observed. This sustained degradation stands in marked contrast to the kinetic profile of the VHL-ligand based RIPK1 degrader **LD5097**, which we have previously characterized^23^. Following washout, LD5097 permits RIPK1 to re-accumulate relatively quickly, reaching ∼50% of baseline levels within 12 hours. Given the long endogenous half-life of RIPK1 (Table S2), this rebound is unlikely to reflect rapid de novo resynthesis; rather, it indicates that LD5097 is cleared from cells shortly after washout, so that active RIPK1 degradation is not maintained. The prolonged activity of **LD5095**, by contrast, indicates that it is retained intracellularly for an extended period, exhibiting a significantly longer residence time than LD5097 and thereby sustaining RIPK1 degradation well after washout.

### LD5095 is a Highly Selective RIPK1 Degrader

The RIPK1 binder utilized in our RIPK1 degraders is recognized as a typical type II kinase inhibitor, exhibiting binding activity to several off-target kinases, including TrkA, Flt1, Flt4, Ret, Met, Mer, Fak, FGFR1, and MLK1.^39^ To evaluate the specificity of **LD5095**, we conducted mass spectrometry (MS)-based analysis of the entire cellular proteome of MOLM14 cells, a cell line known to express the entire kinome.^45^ Following treatment with either **LD5095** (100 nM) or DMSO for 6 hours, we successfully detected ∼ 17,000 proteome isoforms. Our results revealed that **LD5095** degrades RIPK1 with high specificity, as indicated by the red dot in Figure 4, with no observed degradation of off-target kinases (Table S1). This finding is consistent with previous studies suggesting that PROTACs with potentially promiscuous target protein binders may achieve enhanced selectivity through protein-protein interactions involving the E3 ligase. ^45–47^ Using a “dirty” warhead to achieve specific degradation has been well documented in the literature.^48^ Unlike small-molecule inhibitors, effective targeted protein degradation requires the formation of a productive ternary complex among the target protein, the PROTAC molecule, and the recruited E3 ligase. Degradation selectivity is therefore determined not only by warhead binding affinity, but also by additional factors, including linker length and composition, the relative orientation of the target protein and E3 ligase, and the stability and cooperativity of the ternary complex. We speculate that although **LD5095** may bind to other kinases, it does not promote productive ternary complex formation or subsequent ubiquitination of these off-target proteins, resulting in minimal off-target degradation. Collectively, these features contribute to the excellent degradation selectivity observed for **LD5095**.

**Figure 4.**
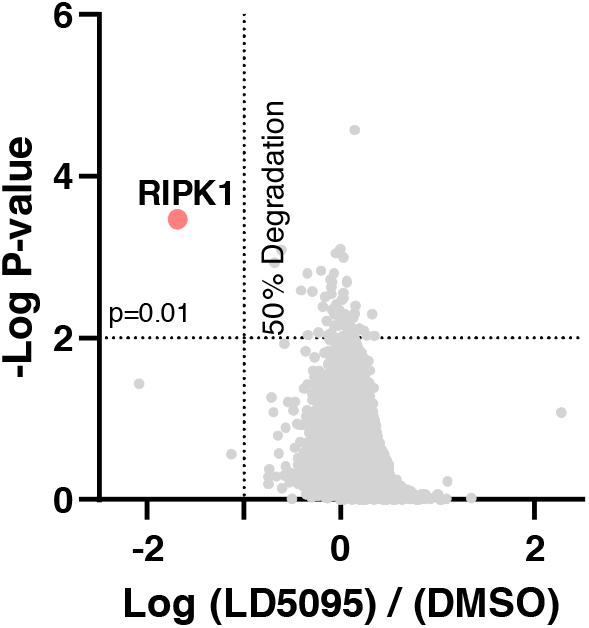
Proteome profiling of **LD5095** induced protein degradation in MOLM14 cells. MOLM14 cells were treated with **LD5095** (100 nM) or DMSO for 6 hrs (n=3). In total, ∼17,000 proteome isoforms were quantified in the proteomics experiment. RIPK1 (red dot) and TIGD4 are two proteins showing >50% degr**ad**ation with P < 0.01.

### LD5095 Sensitizes Cells to TNF-_α_-Mediated Apoptosis Without Intrinsic Cytotoxicity

Genetic ablation of RIPK1 is well-documented to sensitize cells to TNFα -induced apoptosis. To evaluate the pharmacological impact of LD5095, we assessed its effect on Jurkat cell viability and apoptosis in the presence and absence of extrinsic stimuli. Crucially, treatment with LD5095 as a monotherapy did not induce detectable cytotoxicity or growth inhibition (Figure S5), consistent with the role of RIPK1 as a context-dependent signaling scaffold rather than an essential survival protein. However, when combined with TNF-α, LD5095 dramatically reduced cell viability, effectively mirroring the phenotype observed in RIPK1 knockout models (Figure 5a-b). Flow cytometry analysis further confirmed that LD5095 significantly enhanced TNFα -induced apoptosis (Figure 5c). This synergistic effect was characterized by the robust induction of apoptotic markers, including the cleavage of Caspase-3/7 and PARP (Figure 5d). The observation that this cell death was fully reversed by the pan-caspase inhibitor Z-VAD-FMK confirms that the loss of viability is driven specifically by apoptotic pathways (Figure 5b-d). Collectively, these findings demonstrate that LD5095 is a potent and selective sensitizer that primes cells for TNFα -mediated programmed cell death without compromising baseline cellular integrity.

**Figure 5.**
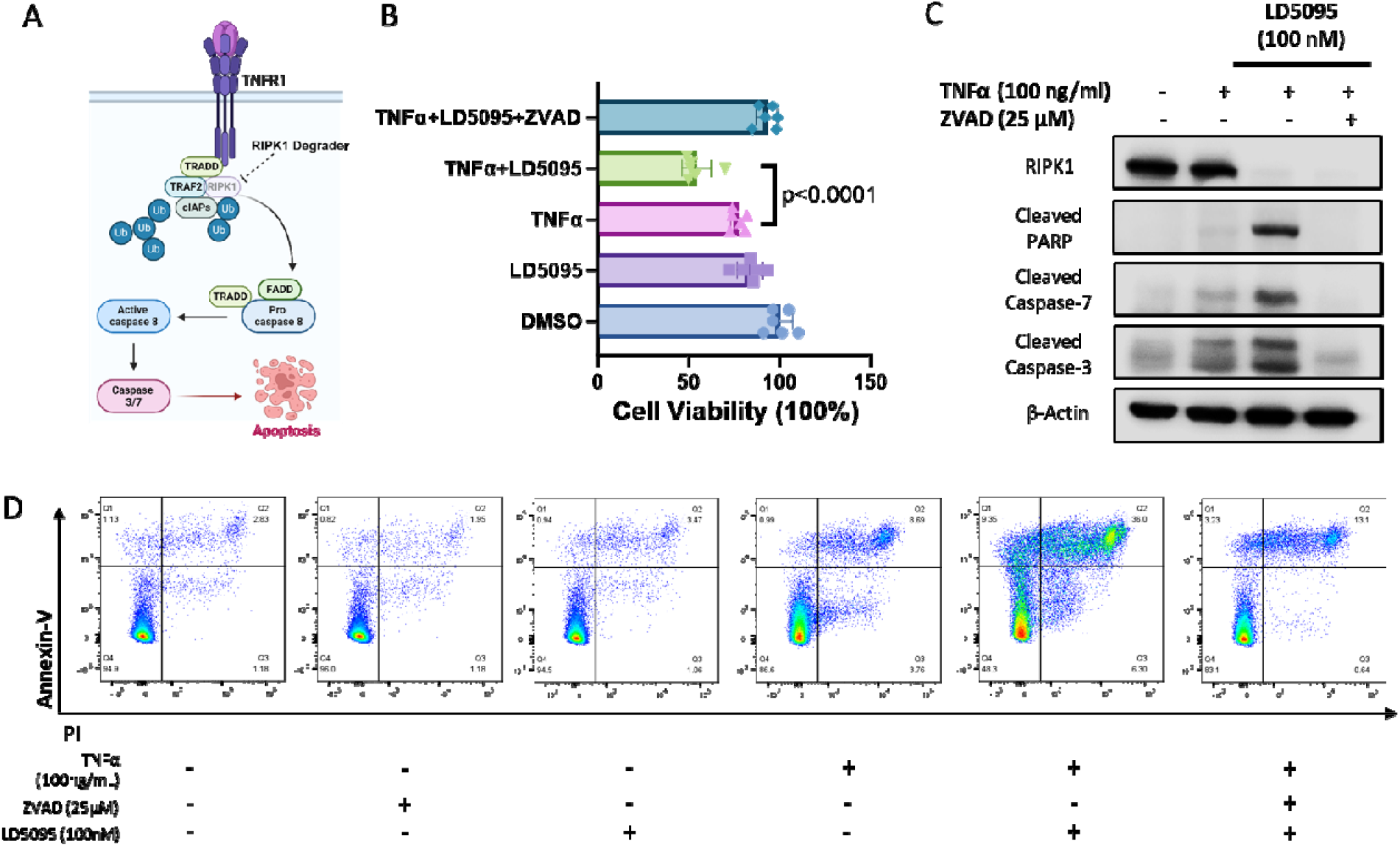
**LD5095** sensitizes Jurkat cells to TNFα-mediated apoptosis. (A) Schematic representation of the role of RIPK1 in cell death pathways. (B) Cell viability assay of Jurkat cells following 72 hours of treatment with the indicated compounds. Treatments were: **LD5095** (100 nM), TNFα (100 ng/mL), and Z-VAD-FMK (25 μM). Data are presented as the mean±SD (n=6 biological repeats). (C) Western blot analysis of apoptosis markers in Jurkat cells after 72 hours of treatment. Blots show the expression levels of cleaved caspase-3, cleaved caspase-7, and cleaved PARP. (D) Representative flow cytometry dot plots illustrating apoptosis detection using Annexin V and Propidium Iodide (PI) staining. Quadrant definitions are as follows: Viable cells (Annexin V-/PI-), early apoptotic cells (Annexin V+/PI-), and late apoptotic cells (Annexin V+/PI+).

### Assessment of the Pharmacodynamic Property of LD5095

Finally, we assessed the pharmacodynamic effect of **LD5095** in Jurkat subcutaneous xenograft tumors in mice. As shown in Figure 6, single administration of our first-generation RIPK1 degrader, **LD4172**, failed to reduce RIPK1 protein levels in tumor tissues. In contrast, a single intravenous dose of **LD5095** (10 mg/kg) resulted in significant and sustained target depletion, reducing RIPK1 protein levels by >55%, 52%, and 85% at 6, 24, and 144 hours, respectively. Notably, robust degradation (85%) persisted even at the 144-hour (6-day) time point, the longest interval tested. This sustained pharmacodynamic response supports the feasibility of a weekly or biweekly dosing regimen in future clinical applications.

**Figure 6.**
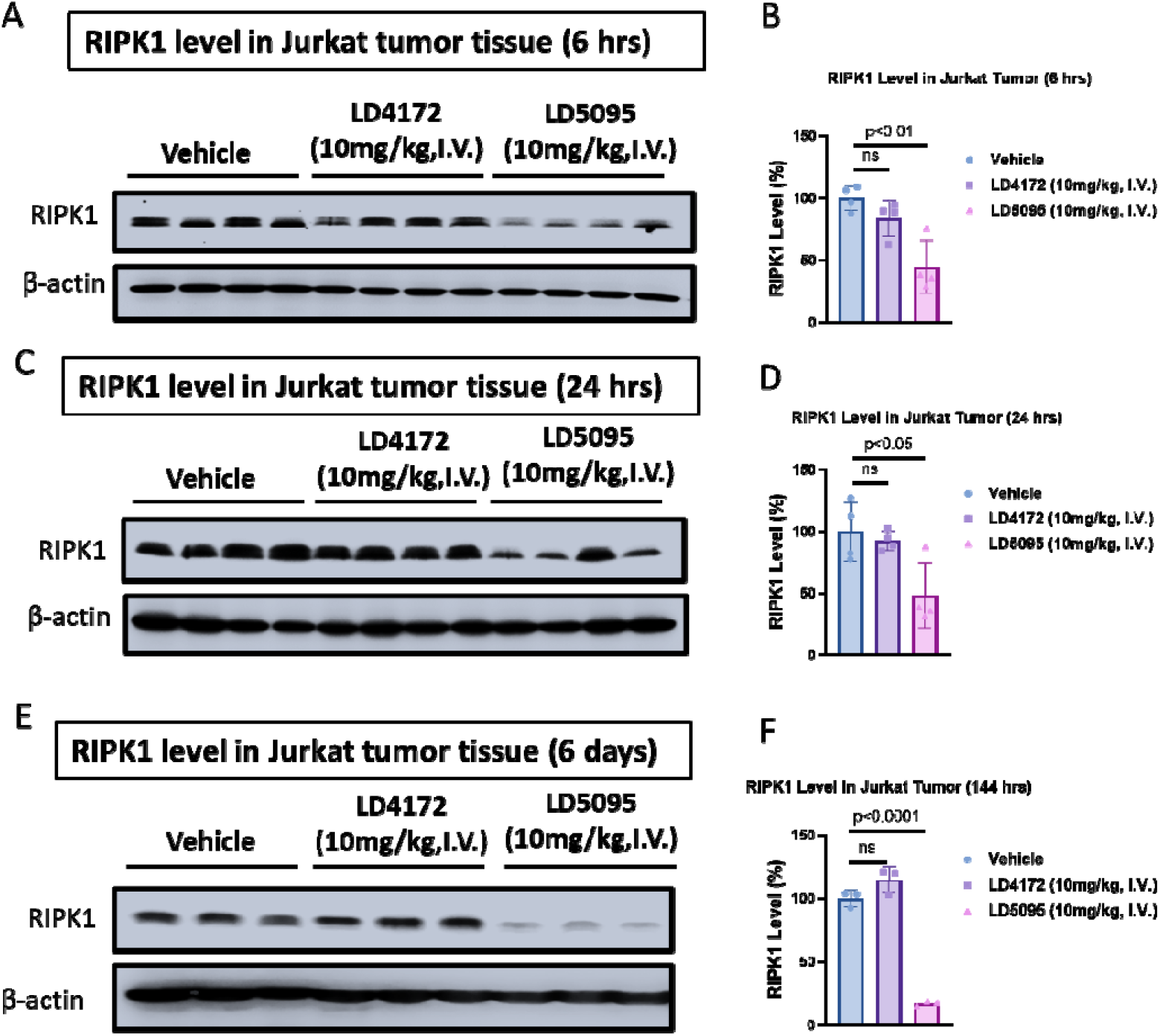
Figure 6. Pharmacodynamic (PD) evaluation of **LD5095** on RIPK1 protein levels in a Jurkat subcutaneous xenograft mouse model. Representative Western blot analyses (A, C, E) and corresponding densitometric quantifications (B, D, F) of RIPK1 protein are shown following single intravenous (IV) administration of vehicle, **LD4172**, or **LD5095** to tumor-bearing nude mice. Tumor tissues were collected at the specified time points and analyzed by Western blotting. Panels A and B show results 6 hours after a 10 mg/kg dose (n=4 per group); Panels C and D show results 24 hours after a 10 mg/kg dose (n=4 per group); and panels E and F show results 144 hours after a 10 mg/kg dose (n=3 per group). Data in panels B, D, and F are presented as the mean ±standard deviation (SD). ns indicates no statistical significance.

### Species-Dependent Degradation Activity of LD5095

Despite demonstrating potent and selective activity in human cellular models, **LD5095** exhibited negligible activity in murine B16F10 cells (Figure S4). This marked species discrepancy is a well-documented phenomenon for Cereblon-based degraders, including clinically approved IMiDs like pomalidomide and lenalidomide^49–51^, and has been similarly observed with our previously reported degrader, **LD5097**^23^. This species-specific activity profile currently precludes the evaluation of **LD5095** in standard immunocompetent murine syngeneic models as we demonstrated previously^18^.

## DISCUSSION AND CONCLUSION

In summary, we have successfully designed, synthesized, and evaluated a series of CRBN-recruiting RIPK1 degraders, representing the first such class to be reported. Optimization of the linker architecture, exit vector, and CRBN ligand led to the discovery of **LD5095** as a lead compound with high potency, achieving DC50 values of 1.4 nM, 1.2 nM, and 2.3 nM in Jurkat, MOLM-14, and U937 cancer cell lines, respectively. Mechanistic validation using a comprehensive suite of controls—including the standalone RIPK1 warhead (T2I), CRBN ligand (CRBNL), and inhibitors of the E1-E3-proteasome cascade (TAK-243, MLN4924, and Carfilzomib)—definitively confirmed that **LD5095** induces RIPK1 degradation through specific ternary complex formation and the ubiquitin-proteasome system (UPS). Notably, **LD5095** demonstrated prolonged cellular activity with minimal protein recovery observed 72 hours post-washout and exhibited high proteomic selectivity for RIPK1. A defining feature of LD5095 is its context-dependent biological activity.

While LD5095 significantly enhances TNF-α-mediated apoptosis, it does not induce significant growth inhibition as a monotherapy in cancer cell lines or human primary cells, including B cells, T cells, and PBMCs. This lack of acute cytotoxicity in unstimulated cells reflects the specialized role of RIPK1 as a signaling scaffold rather than an essential survival protein. This profile suggests a favorable therapeutic window, where **LD5095** may “prime” the tumor microenvironment for immunogenic cell death (ICD) without compromising the viability of healthy, non-transformed tissues.

The pharmacological profile of **LD5095** further supports its translational potential. Following intravenous (IV) administration, LD5095 exhibited excellent metabolic stability characterized by low clearance (Cl_plasma = 4.86 mL/min/kg) and a prolonged terminal half-life (t_1/2 =_ 21.2 h). Upon oral administration at 20 mg/kg, the compound achieved a C_max_ of 304.3 ng/mL with a t_1/2_ of approximately 7.3 hours. Furthermore, our PD study revealed that single administration effectively reduced RIPK1 levels in Jurkat xenografts for over 6 days. This sustained effect is likely a result of the synergy between efficient cellular degradation, extended intracellular retention, and the molecule’s high intrinsic metabolic stability.

We did not employ a cycloheximide (CHX) chase to measure RIPK1 turnover, because the endogenous half-life of RIPK1 has already been quantified by dynamic SILAC metabolic labeling across multiple human primary cells and cell lines, ranging from roughly 40 to 75 hours (Table S2). In contrast to a CHX chase— which globally arrests translation and becomes cytotoxic over the multi-day window that a protein with such a long half-life would demand—dynamic SILAC reports turnover under physiological, synthesis-competent conditions, rendering an additional CHX experiment both redundant and technically ill-suited for this target. Importantly, this relatively slow basal turnover means that once LD5095 depletes RIPK1, restoration to baseline is inherently slow. This intrinsic property, together with the prolonged intracellular retention and high metabolic stability of LD5095, accounts for the durable suppression of RIPK1 observed for over 6 days in Jurkat xenografts following a single dose.

The lack of activity in standard murine models mirrors observations with our previously reported degrader, LD5097. This species discrepancy appears correlated with linker rigidity; while the flexible-linker PROTAC LD4172 retains potency across species^18^, the rigid-linker architecture of **LD5095** imposes stricter conformational constraints that may be sensitive to human-murine ortholog variations. To accurately validate its ability to potentiate antitumor immunity, future studies will utilize humanized immune system mouse models (e.g., huCD34^+^ engrafted mice). This approach will allow for the evaluation of **LD5095**-mediated immune activation and synergy with immune checkpoint inhibitors (ICIs) within a fully humanized context. Overall, **LD5095** emerges as a promising candidate for cancer treatment and a valuable tool for further elucidating RIPK1 biology.

## EXPERIMENTAL SECTION

### Chemistry

All reagents and solvents employed were purchased commercially and used as received. Reagents were purified prior to use unless otherwise stated. Column chromatography was carried out on a Yamazen Smart Flash EPCLC W-Prep 2XY or Agilent 1260 Infinity Preparative LC System. ^1^H NMR and ^13^C NMR spectral data were recorded in CDCl_3_, acetone-*d*_*6*_, DMSO-*d*_*6*_, CD_3_OD on a Varian Palo Alto 400MHz NMR spectrometer and Chemical shifts (δ) were reported in parts per million (ppm), and the signals were described as brs (broad singlet), d (doublet), dd (doublet of doublet), m (multiple), q (quarter), s (singlet), and t (triplet). Coupling constants (J values) were given in Hz. Low-resolution mass spectra (ESI) was obtained using Agilent LC-MS 1200 series (6130 single quad). High-resolution mass spectra (ESI) was obtained using ThermoFisher Orbitrap Fusion Lumos Tribrid Mass Spectrometer. All final compounds had purity >95% determined by using High Pressure Liquid Chromatography (HPLC) using an Agilent Eclipse plus-C18 column eluting with a mixture of MeCN/Water (V:V = 80:20 plus 0.1% FA)

### General Procedures for the preparation of compound (3)

To a solution of 3-(4-fluoro-1,3-dioxo-2,3-dihydro-1H-inden-2-yl)piperidine-2,6-dione (50 mg, 0.18 mmol) in DMF (10 mL) was added amine (0.22 mmol) and DIPEA (51 µL, 0.36 mmol). The mixture was stirred at 90 °C for 2 hours. The reaction was concentrated and then was purified by silica gel flash column chromatography to give the desired compound. The compound was then dissolved in DCM (2 mL) and TFA (0.5 mL). The mixture was stirred at room temperature for 7 hours before it was concentrated under reduced pressure to give compound **1** without further purification.

The solution of compound **1** (0.1 mmol), **2** (62 mg, 0.12 mmol) in DMF (3 mL) was added HATU (45 mg, 0.12 mmol) and DIPEA (70 µL, 0.5 mmol). The mixture was stirred at room temperature for 1 hour. The reaction was concentrated and then was purified by reverse phase preparative HPLC to provide compound **3**.

*N-(3-(4-amino-5-(1-(2-(3-(trifluoromethoxy)phenyl)acetyl)indolin-5-yl)-7H-pyrrolo[2,3-d]pyrimidin-7-yl)propyl)-4-((2-(2,6-dioxopiperidin-3-yl)-1,3-dioxoisoindolin-4-yl)amino)butanamide (****3a***): ^1^H NMR (400 MHz, DMSO-*d*_*6*_) δ 11.06 (s, 1H), 8.07 (d, *J* = 8.1 Hz, 2H), 7.91 (s, 1H), 7.51 (t, *J* = 7.7 Hz, 1H), 7.44 (t, *J* = 7.8 Hz, 1H), 7.29 (d, *J* = 12.8 Hz, 4H), 7.21 (dd, *J* = 14.2, 8.5 Hz, 2H), 7.07 (d, *J* = 8.5 Hz, 1H), 6.96 (d, J = 7.0 Hz, 1H), 6.59 (brs, 1H), 6.03 (s, 1H), 5.01 (dd, *J* = 12.7, 5.0 Hz, 1H), 4.20 (t, *J* = 8.0 Hz, 2H), 4.12 (s, 2H), 3.93 (s, 2H), 3.27 (d, *J* = 9.0 Hz, 2H), 3.20 (d, *J* = 7.2 Hz, 2H), 3.01 (d, *J* = 5.6 Hz, 2H), 2.84 (t, *J* = 13.3 Hz, 1H), 2.54 (d, *J* = 16.5 Hz, 2H), 2.14 (t, *J* = 6.7 Hz, 2H), 2.02 – 1.93 (m, 1H), 1.91 – 1.82 (m, 2H), 1.81 – 1.70 (m, 2H). ^13^C NMR (100 MHz, DMSO-*d*_*6*_) δ 173.2, 172.0, 170.5, 169.2, 168.8, 167.7, 157.62, 151.9, 150.4, 148.6, 146.7, 142.1, 138.4, 136.6, 133.2, 132.6, 130.4, 130.3, 126.4 (q, *J* = 222 Hz), 122.6, 119.4, 117.5, 116.5, 115.4, 110.7, 109.5, 100.3, 48.9, 48.2, 42.1, 42.0, 41.6, 36.4, 32.9, 31.4, 30.2, 27.9, 25.1, 22.5. Purity: 98.8%. MS (ESI): m/z 852.3 [M+H]^+^. HRMS (ESI): m/z Calcd for (C_43_H_41_F_3_N_9_O_7_) ([M+H]^+^): 852.3081; found: 852.3068.

*N-(3-(4-amino-5-(1-(2-(3-(trifluoromethoxy)phenyl)acetyl)indolin-5-yl)-7H-pyrrolo[2,3-d]pyrimidin-7-yl)propyl)-6-((2-(2,6-dioxopiperidin-3-yl)-1,3-dioxoisoindolin-4-yl)amino)hexanamide (****3b***): ^1^H NMR (400 MHz, DMSO-*d*_*6*_) δ 11.06 (s, 1H), 8.07 (d, *J* = 10.5 Hz, 2H), 7.83 (s, 1H), 7.51 (t, *J* = 7.8 Hz, 1H), 7.44 (t, *J* = 7.9 Hz, 1H), 7.29 (d, *J* = 13.4 Hz, 4H), 7.21 (dd, *J* = 14.9, 8.3 Hz, 2H), 7.03 (d, *J* = 8.6 Hz, 1H), 6.96 (d, *J* = 7.0 Hz, 1H), 6.48 (s, 1H), 6.02 (brs, 1H), 5.05 – 4.95 (m, 1H), 4.20 (t, *J* = 8.0 Hz, 2H), 4.11 (s, 2H), 3.93 (s, 2H), 3.28 (s, 2H), 3.25 – 3.20 (m, 2H), 2.99 (d, *J* = 5.6 Hz, 2H), 2.83 (t, *J* = 13.4 Hz, 1H), 2.54 (d, *J* = 17.2 Hz, 2H), 2.04 (t, *J* = 6.9 Hz, 2H), 2.01 – 1.93 (m, 1H), 1.90 – 1.80 (m, 2H), 1.52 (s, 4H), 1.29 (d, *J* = 6.8 Hz, 2H). ^13^C NMR (100 MHz, DMSO-*d*_*6*_) δ 173.2, 172.4, 170.5, 169.3, 168.8, 167.7, 157.6, 151.9, 150.4, 146.8, 142.1, 138.4, 136.6, 133.2, 132.5, 130.4, 130.3, 126.4 (q, *J* = 222 Hz), 123.6, 122.6, 119.4, 117.5, 116.5, 115.4, 110.7, 109.4, 100.3, 48.9, 48.2, 42.1, 41.6, 36.3, 35.8, 31.4, 30.3, 28.9, 27.9, 26.3, 25.4, 22.5. Purity: >99.0%. MS (ESI): m/z 880.3 [M+H]^+^. HRMS (ESI): m/z Calcd for (C_45_H_45_F_3_N_9_O_7_) ([M+H]^+^): 880.3394; found: 880.3383.

*N-(3-(4-amino-5-(1-(2-(3-(trifluoromethoxy)phenyl)acetyl)indolin-5-yl)-7H-pyrrolo[2,3-d]pyrimidin-7-yl)propyl)-9-((2-(2,6-dioxopiperidin-3-yl)-1,3-dioxoisoindolin-4-yl)amino)nonanamide (****3c***): ^1^H NMR (400 MHz, DMSO-*d*_*6*_) δ 11.07 (s, 1H), 8.07 (d, *J* = 8.8 Hz, 2H), 7.81 (s, 1H), 7.52 (t, *J* = 7.7 Hz, 1H), 7.44 (t, *J* = 7.8 Hz, 1H), 7.29 (d, *J* = 12.3 Hz, 3H), 7.24 – 7.16 (m, 2H), 7.02 (d, *J* = 8.7 Hz, 1H), 6.97 (d, *J* = 6.9 Hz, 1H), 6.47 (s, 1H), 6.03 (brs, 1H), 5.01 (dd, *J* = 12.7, 5.0 Hz, 1H), 4.20 (t, *J* = 8.0 Hz, 2H), 4.10 (d, *J* = 6.1 Hz, 2H), 3.93 (s, 2H), 3.20 (d, *J* = 6.9 Hz, 4H), 3.00 (d, *J* = 5.6 Hz, 2H), 2.84 (t, *J* = 13.3 Hz, 1H), 2.54 (d, *J* = 16.6 Hz, 2H), 2.01 (t, *J* = 6.9 Hz, 3H), 1.90 – 1.79 (m, 2H), 1.50 (d, *J* = 6.9 Hz, 2H), 1.45 (d, *J* = 7.2 Hz, 2H), 1.23 (d, *J* = 12.0 Hz, 8H). ^13^C NMR (100 MHz, DMSO-*d*_*6*_) δ 173.2, 172.5, 170.5, 169.3, 168.8, 167.7, 157.6, 151.9, 150.4, 148.6, 146.8, 142.1, 138.4, 136.6, 133.2, 132.5, 130.4, 130.3, 126.4 (q, *J* = 222 Hz), 123.6, 122.6, 119.4, 117.5, 116.5, 115.4, 110.7, 109.4, 48.9, 48.2, 42.2, 42.0, 41.6, 36.3, 35.8, 31.4, 30.3, 29.2, 29.1, 29.0, 27.9, 26.7, 25.6, 22.5. Purity: 95.2%. MS (ESI): m/z 922.4 [M+H]^+^. HRMS (ESI): m/z Calcd for (C_48_H_51_F_3_N_9_O_7_) ([M+H]^+^): 922.3864; found: 922.3854.

*N-(3-(4-amino-5-(1-(2-(3-(trifluoromethoxy)phenyl)acetyl)indolin-5-yl)-7H-pyrrolo[2,3-d]pyrimidin-7-yl)propyl)-11-((2-(2,6-dioxopiperidin-3-yl)-1,3-dioxoisoindolin-4-yl)amino)undecanamide (****3d***): ^1^H NMR (400 MHz, DMSO-*d*_*6*_) δ 11.07 (s, 1H), 8.07 (d, *J* = 8.7 Hz, 2H), 7.81 (s, 1H), 7.52 (t, *J* = 7.6 Hz, 1H), 7.44 (t, *J* = 7.8 Hz, 1H), 7.29 (d, *J* = 12.3 Hz, 3H), 7.24 – 7.17 (m, 2H), 7.03 (d, *J* = 8.6 Hz, 1H), 6.97 (d, *J* = 7.0 Hz, 1H), 6.47 (s, 1H), 6.04 (brs, 1H), 5.01 (d, *J* = 7.9 Hz, 1H), 4.20 (t, *J* = 8.0 Hz, 2H), 4.11 (s, 2H), 3.93 (s, 2H), 3.24 – 3.17 (m, 4H), 2.99 (d, *J* = 5.4 Hz, 2H), 2.85 (s, 1H), 2.70 – 2.50 (m, 2H), 2.00 (d, *J* = 6.6 Hz, 3H), 1.89 – 1.80 (m, 2H), 1.47 (m, 4H), 1.21 (m, 12H). ^13^C NMR (100 MHz, DMSO-*d*_*6*_) δ 173.2, 172.5, 170.5, 169.3, 168.8, 167.7, 151.9, 150.4, 148.6, 146.8, 142.1, 138.4, 136.6, 133.2, 132.6, 130.4, 130.3, 129.4, 126.4 (q, *J* = 222 Hz), 123.6, 122.6, 119.4, 117.5, 116.6, 115.4, 110.7, 109.4, 100.3, 48.9, 48.2, 42.2, 42.0, 41.6, 36.3, 35.9, 31.4, 30.3, 29.4, 29.3, 29.2, 29.0, 27.9, 26.7, 25.6, 22.5. Purity: 97.8%. MS (ESI): m/z 950.4 [M+H]^+^. HRMS (ESI): m/z Calcd for (C_50_H_55_F_3_N_9_O_7_) ([M+H]^+^): 950.4177; found: 950.4170.

### General Procedures for the preparation of 11-amino-N-(3-(4-amino-5-(1-(2-(3-(trifluoromethoxy)phenyl)acetyl)indolin-5-yl)-7H-pyrrolo[2,3-d]pyrimidin-7-yl)propyl)undecanamide (4)

To a solution of **2** (200 mg, 0.4 mmol), 9-((tert-butoxycarbonyl)amino)nonanoic acid (132 mg, 0.5 mmol) in DMF (10 mL) was added HATU (480 mg, 0.6 mmol) and DIPEA (172 µL, 1.2 mmol). The mixture was stirred at room temperature for 1 hour and was concentrated and then was purified by silica gel flash column chromatography to give the desired compound. The compound was then dissolved in DCM (5 mL) and TFA (1 mL). The mixture was stirred at room temperature for 3 hours before it was concentrated under reduced pressure to give compound **4** without further purification (119 mg, 43% yield, two steps). ^1^H NMR (400 MHz, DMSO-*d*_*6*_) δ 8.54 – 8.46 (m, 1H), 7.92 (s, 5H), 7.70 (d, *J* = 10.6 Hz, 1H), 7.50 – 7.44 (m, 1H), 7.31 (d, *J* = 12.4 Hz, 2H), 7.26 (s, 2H), 4.24 (s, 2H), 3.98 (s, 1H), 3.09 (d, *J* = 4.4 Hz, 2H), 3.03 (d, *J* = 5.5 Hz, 2H), 2.72 (s, 2H), 2.03 (t, *J* = 6.7 Hz, 2H), 1.97 – 1.85 (m, 2H), 1.50 (d, *J* = 6.9 Hz, 2H), 1.45 (s, 2H), 1.28 – 1.24 (m, 8H), 1.23 (s, 4H). MS (ESI): m/z 694.4 [M+H]^+^.

### General Procedures for the preparation of compound (5)

To a solution of **4** (28 mg, 0.04 mmol) and CRBN ligand acid (0.05 mmol) in DMF (3 mL) was added HATU (22 mg, 0.06 mmol) and DIPEA (34 µL, 0.24 mmol). The mixture was stirred at room temperature for 1 hour. The reaction was concentrated and then was purified by reverse phase preparative HPLC to provide the title compound **5**.

*N-(3-(4-amino-5-(1-(2-(3-(trifluoromethoxy)phenyl)acetyl)indolin-5-yl)-7H-pyrrolo[2,3-d]pyrimidin-7-yl)propyl)-11-(2-((2-(2,6-dioxopiperidin-3-yl)-1,3-dioxoisoindolin-5-yl)amino)acetamido)undecanamide (****5a***): ^1^H NMR (400 MHz, DMSO-*d*_*6*_) δ 11.03 (s, 1H), 8.08 (d, *J* = 9.0 Hz, 2H), 7.97 (s, 1H), 7.80 (s, 1H), 7.54 (d, *J* = 8.2 Hz, 1H), 7.44 (t, *J* = 7.8 Hz, 1H), 7.29 (d, *J* = 12.9 Hz, 4H), 7.22 (d, *J* = 11.5 Hz, 2H), 6.89 (s, 1H), 6.81 (d, *J* = 8.3 Hz, 1H), 6.03 (brs, 1H), 5.04 – 4.94 (m, 1H), 4.20 (t, *J* = 8.0 Hz, 2H), 4.11 (s, 2H), 4.07 (d, *J* = 5.2 Hz, 1H), 3.93 (s, 2H), 3.77 (s, 2H), 3.19 (t, *J* = 7.8 Hz, 2H), 3.13 (d, *J* = 5.3 Hz, 2H), 3.00 (s, 2H), 2.84 (t, *J* = 13.2 Hz, 1H), 2.53 (m, 2H), 2.00 (t, *J* = 7.1 Hz, 2H), 1.95 (d, *J* = 12.0 Hz, 1H), 1.90 – 1.80 (m, 2H), 1.41 (m, 2H), 1.32 (m, 2H), 1.15 (m, 12H). ^13^C NMR (100 MHz, DMSO-*d*_*6*_) δ 173.2, 172.5, 170.5, 169.0, 168.9, 168.0, 167.5, 157.6, 154.6, 151.9, 150.4, 148.6, 142.1, 138.4, 134.3, 130.5 (q, *J* = 188 Hz), 130.4, 130.3, 125.3, 123.6, 122.6, 119.4, 117.2, 116.5, 115.4, 100.3, 49.0, 48.2, 46.3, 42.0, 41.6, 38.9, 36.3, 35.9, 31.4, 30.3, 29.5, 29.4, 29.3, 29.2, 29.1, 27.9, 26.7, 25.7, 22.6. Purity: 95.4%. MS (ESI): m/z 1007.4 [M+H]^+^. HRMS (ESI): m/z Calcd for (C_52_H_58_F_3_N_10_O_8_) ([M+H]^+^): 1007.4391; found: 1007.4383.

*N-(3-(4-amino-5-(1-(2-(3-(trifluoromethoxy)phenyl)acetyl)indolin-5-yl)-7H-pyrrolo[2,3-d]pyrimidin-7-yl)propyl)-11-(2-(4-(4-((2,6-dioxopiperidin-3-yl)amino)phenyl)piperidin-1-yl)acetamido)undecanamide (****5b***): ^1^H NMR (400 MHz, DMSO-*d*_*6*_) δ 10.78 (s, 1H), 8.12 (d, *J* = 9.3 Hz, 2H), 7.84 (s, 1H), 7.69 (s, 1H), 7.48 (t, *J* = 7.8 Hz, 1H), 7.33 (d, *J* = 13.9 Hz, 4H), 7.29 – 7.21 (m, 2H), 6.94 (d, *J* = 7.6 Hz, 2H), 6.60 (d, *J* = 7.9 Hz, 2H), 6.07 (brs, 1H), 5.65 (d, *J* = 7.1 Hz, 1H), 4.24 (t, *J* = 8.2 Hz, 3H), 4.15 (s, 2H), 3.97 (s, 2H), 3.23 (t, *J* = 8.0 Hz, 2H), 3.17 (s, 1H), 3.05 (dd, *J* = 15.0, 6.5 Hz, 4H), 2.89 (s, 2H), 2.84 (d, *J* = 10.6 Hz, 2H), 2.78 – 2.68 (m, 1H), 2.57 (m, 1H), 2.35 – 2.24 (m, 1H), 2.09 (d, *J* = 8.6 Hz, 3H), 2.04 (t, *J* = 7.3 Hz, 2H), 1.85 (m, 3H), 1.64 (s, 4H), 1.47 (s, 2H), 1.39 (m, 2H), 1.22 (m, 12H). ^13^C NMR (100 MHz, DMSO-*d*_*6*_) δ 174.1, 173.5, 172.5, 169.5, 168.8, 157.6, 151.9, 150.4, 148.6, 146.4, 142.1, 138.4, 134.4, 130.5 (q, *J* = 209 Hz), 130.4, 130.3, 127.5, 125.3, 123.6, 122.6, 119.4, 116.6, 115.4, 113.0, 100.3, 62.1, 54.5, 53.0, 48.2, 42.0, 41.6, 40.8, 38.5, 36.3, 35.9, 33.7, 31.1, 30.3, 29.6, 29.4, 29.3, 29.2, 29.1, 27.9, 26.7, 25.7, 25.1. Purity: 97.6%. MS (ESI): m/z 1021.5 [M+H]^+^. HRMS (ESI): m/z Calcd for (C_55_H_68_F_3_N_10_O_6_) ([M+H]^+^): 1021.5275; found: 1021.5258.

*N-(3-(4-amino-5-(1-(2-(3-(trifluoromethoxy)phenyl)acetyl)indolin-5-yl)-7H-pyrrolo[2,3-d]pyrimidin-7-yl)propyl)-11-(2-(4-(2,6-dioxopiperidin-3-yl)phenoxy)acetamido)undecanamide (****5c***): ^1^H NMR (400 MHz, DMSO-*d*_*6*_) δ 10.78 (s, 1H), 8.08 (d, *J* = 9.3 Hz, 2H), 8.00 (s, 1H), 7.81 (s, 1H), 7.44 (t, *J* = 7.8 Hz, 1H), 7.29 (d, *J* = 12.2 Hz, 4H), 7.25 – 7.17 (m, 2H), 7.10 (d, *J* = 8.4 Hz, 2H), 6.86 (d, *J* = 8.5 Hz, 2H), 6.03 (brs, 1H), 4.40 (s, 2H), 4.20 (t, *J* = 8.1 Hz, 2H), 4.15 – 4.07 (m, 2H), 3.93 (s, 2H), 3.75 (dd, *J* = 11.3, 4.5 Hz, 1H), 3.19 (t, *J* = 8.0 Hz, 2H), 3.13 (d, *J* = 5.2 Hz, 1H), 3.08 – 3.02 (m, 2H), 3.00 (d, *J* = 5.8 Hz, 2H), 2.60 (m, 1H), 2.11 (m, 1H), 2.01 (t, *J* = 7.2 Hz, 2H), 1.96 (d, *J* = 8.4 Hz, 1H), 1.90 – 1.81 (m, 2H), 1.44 (m, 2H), 1.36 (m, 2H), 1.18 (m, 12H). ^13^C NMR (100 MHz, DMSO-*d*_*6*_) δ 174.8, 173.8, 172.5, 168.8, 167.8, 157.6, 157.1, 151.9, 150.4, 148.6, 142.1, 138.4, 132.2, 130.5 (q, *J* = 188 Hz), 130.4, 130.3, 129.9, 125.3, 123.6, 122.6, 119.4, 116.6, 115.4, 114.9, 100.3, 67.4, 48.2, 46.9, 42.0, 41.6, 38.6, 36.3, 35.9, 31.8, 30.3, 29.5, 29.4, 29.3, 29.2, 29.1, 27.9, 26.7, 26.4, 25.7. Purity: 96.5%. MS (ESI): m/z 939.4 [M+H]^+^. HRMS (ESI): m/z Calcd for (C_50_H_58_F_3_N_8_O_7_) ([M+H]^+^): 939.4381; found: 939.4370.

*6-(4-((11-((3-(4-amino-5-(1-(2-(3-(trifluoromethoxy)phenyl)acetyl)indolin-5-yl)-7H-pyrrolo[2,3-d]pyrimidin-7-yl)propyl)amino)-11-oxoundecyl)carbamoyl)piperidin-1-yl)-N-(2,6-dioxopiperidin-3-yl)picolinamide (****5d***): ^1^H NMR (400 MHz, DMSO-*d*_*6*_) δ 10.90 (s, 1H), 8.76 (d, *J* = 8.2 Hz, 1H), 8.11 (d, *J* = 8.7 Hz, 2H), 7.85 (s, 1H), 7.78 (s, 1H), 7.66 (t, *J* = 7.9 Hz, 1H), 7.48 (t, *J* = 7.9 Hz, 1H), 7.33 (d, *J* = 12.3 Hz, 4H), 7.28 – 7.21 (m, 3H), 7.04 (d, *J* = 8.6 Hz, 1H), 6.07 (brs, 1H), 4.73 (t, *J* = 8.8 Hz, 1H), 4.42 (d, J = 12.3 Hz, 2H), 4.24 (t, *J* = 8.0 Hz, 2H), 4.14 (d, *J* = 6.3 Hz, 2H), 3.97 (s, 2H), 3.23 (t, *J* = 7.9 Hz, 2H), 3.17 (s, 1H), 3.02 (d, *J* = 3.5 Hz, 4H), 2.82 (d, *J* = 12.7 Hz, 2H), 2.53 (s, 1H), 2.35 (d, *J* = 11.4 Hz, 1H), 2.23 (m, 1H), 2.05 (t, *J* = 7.1 Hz, 2H), 1.93 – 1.85 (m, 2H), 1.73 (d, *J* = 11.9 Hz, 2H), 1.55 (d, *J* = 11.9 Hz, 2H), 1.47 (s, 2H), 1.35 (s, 2H), 1.22 (m, 14H). ^13^C NMR (100 MHz, DMSO-*d*_*6*_) δ 174.3, 173.4, 172.7, 172.5, 168.9, 164.6, 158.1, 157.6, 151.9, 150.4, 148.6, 147.9, 142.1, 139.0, 138.4, 130.5 (q, *J* = 189 Hz), 130.4, 130.3, 125.3, 123.6, 122.6, 119.4, 116.6, 115.4, 111.0, 110.7, 100.3, 49.8, 48.2, 44.8, 42.5, 42.0, 41.7, 38.7, 36.3, 35.9, 31.4, 30.3, 29.5, 29.4, 29.3, 29.2, 29.1, 28.4, 27.9, 26.7, 25.7, 24.4. Purity: >99.0%. MS (ESI): m/z 1036.5 [M+H]^+^. HRMS (ESI): m/z Calcd for (C_54_H_65_F_3_N_11_O_7_) ([M+H]^+^): 1036.5021; found: 1036.5010.

*5-(4-((11-((3-(4-amino-5-(1-(2-(3-(trifluoromethoxy)phenyl)acetyl)indolin-5-yl)-7H-pyrrolo[2,3-d]pyrimidin-7-yl)propyl)amino)-11-oxoundecyl)carbamoyl)piperidin-1-yl)-N-(2,6-dioxopiperidin-3-yl)picolinamide (****5e***): ^1^H NMR (400 MHz, DMSO-*d*_*6*_) δ 10.81 (s, 1H), 8.57 (s, 1H), 8.49 (d, *J* = 8.2 Hz, 1H), 8.08 (d, *J* = 9.5 Hz, 2H), 7.90 (d, *J* = 8.7 Hz, 1H), 7.81 (s, 1H), 7.73 (s, 1H), 7.44 (t, *J* = 7.8 Hz, 1H), 7.31 (s, 1H), 7.28 (s, 3H), 7.25 – 7.17 (m, 2H), 6.83 (d, *J* = 9.2 Hz, 1H), 6.04 (brs, 1H), 4.71 (s, 1H), 4.37 (d, *J* = 12.8 Hz, 2H), 4.20 (t, *J* = 7.9 Hz, 2H), 4.11 (s, 2H), 3.93 (s, 2H), 3.20 (t, *J* = 8.0 Hz, 2H), 3.00 – 2.93 (m, 4H), 2.85 (s, 2H), 2.76 – 2.66 (m, 1H), 2.51 (s, 1H), 2.35 (s, 1H), 2.00 (d, *J* = 7.0 Hz, 2H), 1.88 – 1.83 (m, 2H), 1.67 (d, *J* = 12.6 Hz, 2H), 1.45 (d, *J* = 11.9 Hz, 4H), 1.31 (s, 2H), 1.18 (m, 14H). ^13^C NMR (100 MHz, DMSO-*d*_*6*_) δ 174.2, 173.5, 172.8, 172.5, 168.9, 165.2, 160.1, 157.6, 151.9, 150.5, 148.6, 142.2, 138.4, 136.9, 130.5 (q, *J* = 187 Hz), 130.4, 125.3, 123.6, 122.6, 119.4, 117.8, 116.6, 115.4, 105.8, 100.3, 49.7, 48.3, 44.5, 42.5, 42.0, 41.6, 38.7, 36.3, 35.9, 31.4, 30.3, 29.6, 29.4, 29.3, 29.2, 29.1, 28.4, 27.9, 26.7, 25.7, 24.8. Purity: 95.0%. MS (ESI): m/z 1036.5 [M+H]^+^. HRMS (ESI): m/z Calcd for (C_54_H_65_F_3_N_11_O_7_) ([M+H]^+^): 1036.5021; found: 1036.5012.

*N-(11-((3-(4-amino-5-(1-(2-(3-(trifluoromethoxy)phenyl)acetyl)indolin-5-yl)-7H-pyrrolo[2,3-d]pyrimidin-7-yl)propyl)amino)-11-oxoundecyl)-6-(2,4-dioxo-1,3,7-triazaspiro[4*.*4]nonan-7-yl)nicotinamide (****5f***): ^1^H NMR (400 MHz, DMSO-*d*_*6*_) δ 10.81 (s, 1H), 8.54 (s, 1H), 8.44 (s, 1H), 8.14 (s, 1H), 8.08 (d, *J* = 9.1 Hz, 2H), 7.90 (d, *J* = 8.8 Hz, 1H), 7.81 (s, 1H), 7.44 (t, *J* = 7.7 Hz, 1H), 7.29 (d, *J* = 12.4 Hz, 4H), 7.25 – 7.17 (m, 2H), 6.46 (d, *J* = 8.8 Hz, 1H), 6.03 (brs, 1H), 4.20 (t, *J* = 7.9 Hz, 2H), 4.11 (s, 2H), 3.93 (s, 2H), 3.67 (d, *J* = 11.8 Hz, 2H), 3.56 (d, *J* = 11.6 Hz, 2H), 3.21 – 3.13 (m, 4H), 2.99 (d, *J* = 5.4 Hz, 2H), 2.32 (m, 1H), 2.11 (d, *J* = 6.0 Hz, 1H), 2.00 (d, *J* = 6.8 Hz, 2H), 1.85 (s, 2H), 1.43 (s, 4H), 1.19 (m, 12H). ^13^C NMR (100 MHz, DMSO-*d*_*6*_) δ 176.7, 172.5, 168.9, 165.3, 157.9, 157.6, 156.6, 151.9, 150.4, 148.6, 148.4, 142.1, 138.4, 136.44, 130.5 (q, *J* = 187 Hz), 130.4, 130.3, 125.3, 123.6, 122.6, 119.4, 118.6, 116.6, 115.4, 105.8, 100.3, 66.7, 55.2, 48.2, 45.8, 42.0, 41.6, 36.3, 35.9, 35.3, 30.3, 29.7, 29.4, 29.3, 29.2, 29.1, 27.9, 26.9, 25.7. Purity: >99.0%. MS (ESI): m/z 952.4 [M+H]^+^. HRMS (ESI): m/z Calcd for (C_49_H_57_F_3_N_11_O_6_) ([M+H]^+^): 952.4445; found: 952.4434.

### General Procedures for the preparation of ethyl 4-(piperidin-4-ylethynyl)benzoate (6)

A solution of ethyl 4-iodobenzoate (552 mg, 2.0 mmol) and tert-butyl 4-ethynylpiperidine-1-carboxylate (460 mg, 2.2 mmol) in DMF (20 mL) was added Pd(PPh_3_)_2_Cl_2_ (140 mg, 0.2 mmol), CuI (38 mg, 0.2 mmol) and TEA (0.58 mL, 4.0 mmol). The resulting suspension was stirred for 8 hours at 100 °C. The mixture was cooled to room temperature and was poured into 60 mL water and the compound was extracted with EtOAc. The organic phase was washed with brine and then dried over MgSO_4_, filtered, and concentrated under reduced pressure. The residue was purified by flash chromatography to afford desired product (515 mg, 72%). MS (ESI): m/z 357.1 [M+H]^+^.

The above product was dissolved in DCM (10 mL) and the solution was added 1 mL TFA. The mixture was stirred for 2 hours at room temperature. The mixture was concentrated under reduced pressure to give compound **6** (326 mg, 88%). ^1^H NMR (400 MHz, DMSO-*d*_*6*_) δ 8.51 (m, 1H), 7.90 (d, *J* = 8.0 Hz, 2H), 7.52 (d, *J* = 7.9 Hz, 2H), 4.28 (q, *J* = 6.9 Hz, 2H), 3.20 (s, 2H), 2.96 (d, *J* = 33.8 Hz, 3H), 2.01 (d, *J* = 11.9 Hz, 2H), 1.74 (d, *J* = 10.1 Hz, 2H), 1.28 (t, *J* = 7.0 Hz, 3H). MS (ESI): m/z 257.1 [M+H]^+^.

### General Procedures for the preparation of 4-((1’-(tert-butoxycarbonyl)-[1,4’-bipiperidin]-4-yl)ethynyl)benzoic acid (7)

To **6** (100 mg, 0.4 mmol) in DCE (10 mL) was added tert-butyl 4-oxopiperidine-1-carboxylate (100 mg, 0.5 mmol) and 1 drop of AcOH. The resulting mixture was stirred at room temperature for 30 mins followed by adding NaBH(OAc)_3_ (170 mg, 0.8 mmol). The reaction mixture was stirred at 50 °C overnight. The reaction mixture was quenched by adding water and extracted with EtOAc. The organic phase was washed with brine and then dried over MgSO_4_, filtered, and concentrated in vacuo. The residue was purified by flash chromatography to afford desired compound (88 mg, 52%). MS (ESI): m/z 427.2 [M+H]^+^.

The above product (88 mg, 0.2 mmol) and LiOH (10 mg, 0.4 mmol) were dissolved in THF (5 mL) and H_2_O (1 mL). This solution was stirred at room temperature overnight. The reaction was concentrated and then was purified by silica gel flash column chromatography to give the title compound **7** (69 mg, 84%). ^1^H NMR (400 MHz, DMSO-*d*_*6*_) δ 7.85 (d, *J* = 7.8 Hz, 2H), 7.42 (d, *J* = 8.1 Hz, 2H), 3.93 (s, 2H), 2.67 (d, *J* = 33.2 Hz, 4H), 2.39 (s, 1H), 2.31 (s, 2H), 1.81 (s, 2H), 1.66 (d, *J* = 12.0 Hz, 2H), 1.61 – 1.49 (m, 2H), 1.35 (s, 9H), 1.26 – 1.18 (m, 3H). MS (ESI): m/z 413.2 [M+H]^+^.

### General Procedures for the preparation of tert-butyl 6-(5-bromopyridin-2-yl)-2,6-diazaspiro[3.3]heptane-2-carboxylate (8)

The mixture of tert-butyl 2,6-diazaspiro[3.3]heptane-2-carboxylate (400 mg, 2.0 mmol) and 5-bromo-2-fluoropyridine (300 mg, 1.7 mmol) in anhydrous DMF (15 mL) was added TEA (0.5 mL, 3.4 mmol) at room temperature. The mixture was stirred at 100 °C for 12 hours. After cooling to room temperature, the mixture was poured into 20 mL water and the compound was extracted with EtOAc. The organic phase was washed with brine and then dried over MgSO_4_, filtered, and concentrated under reduced pressure. The residue was purified by flash chromatography to afford compound **8** (480 mg, 80%). ^1^H NMR (400 MHz, CDCl_3_) δ 8.13 (s, 1H), 7.50 (dd, *J* = 8.8, 2.3 Hz, 1H), 6.17 (d, *J* = 8.8 Hz, 1H), 4.08 (s, 8H), 1.42 (s, 9H). MS (ESI): m/z 354.1, 356.1 [M+H]^+^.

### General Procedures for the preparation of 4-((6-(6-(tert-butoxycarbonyl)-2,6-diazaspiro[3.3]heptan-2-yl)pyridin-3-yl)ethynyl)benzoic acid (9)

A solution of compound 8 (100 mg, 0.28 mmol) and methyl 4-ethynylbenzoate (57 mg, 0.33 mmol) in DMF (10 mL) was added Pd(PPh_3_)_2_Cl_2_ (20 mg, 0.028 mmol), CuI (10 mg, 0.056 mmol) and TEA (0.4 mL, 2.8 mmol). The resulting suspension was stirred for 8 hours at 100 °C. The mixture was cooled to room temperature and was poured into 15 mL water and the compound was extracted with EtOAc. The organic phase was washed with brine and then dried over MgSO_4_, filtered, and concentrated under reduced pressure. The residue was purified by flash chromatography to afford desired product (79 mg, 63%). MS (ESI): m/z 448.2 [M+H]^+^.

The above product (79 mg, 0.18 mmol) and LiOH (7 mg, 0.36 mmol) were dissolved in THF (5 mL) and H_2_O (1 mL). This solution was stirred at room temperature overnight. The reaction was concentrated and then was purified by silica gel flash column chromatography to give the title compound (70 mg, 91%). ^1^H NMR (400 MHz, DMSO-*d*_*6*_) δ 8.25 (s, 1H), 7.90 (d, *J* = 7.6 Hz, 2H), 7.63 (d, *J* = 7.8 Hz, 1H), 7.56 (d, *J* = 7.8 Hz, 2H), 6.36 (d, *J* = 8.6 Hz, 1H), 4.09 (s, 4H), 4.00 (s, 4H), 1.34 (s, 9H). MS (ESI): m/z 420.1 [M+H]^+^.

Compound **10, 11** were prepared according to the procedure described above for compound **9**.

*4-((6-(4-(tert-butoxycarbonyl)piperazin-1-yl)pyridin-3-yl)ethynyl)benzoic acid (****10***): ^1^H NMR (400 MHz, CDCl_3_) δ 8.35 (s, 1H), 7.99 (d, *J* = 8.2 Hz, 2H), 7.60 (d, *J* = 8.7 Hz, 1H), 7.54 (d, *J* = 8.2 Hz, 2H), 6.59 (d, *J* = 8.9 Hz, 1H), 3.59 (d, *J* = 4.7 Hz, 4H), 3.54 (d, *J* = 4.9 Hz, 4H), 1.47 (s, 9H). MS (ESI): m/z 408.2 [M+H]^+^.

*4-((5-((tert-butoxycarbonyl)amino)pyridin-2-yl)ethynyl)benzoic acid (****11***): ^1^H NMR (400 MHz, DMSO-*d*_*6*_) δ 8.11 (s, 1H), 7.88 (d, *J* = 16.4 Hz, 2H), 7.54 (s, 2H), 7.50 (s, 1H), 6.48 (s, 1H), 6.42 (s, 1H). MS (ESI): m/z 239.1 [M+H]^+^.

### General Procedures for the preparation of 4-(6-(6-(tert-butoxycarbonyl)-2,6-diazaspiro[3.3]heptan-2-yl)pyridin-3-yl)benzoic acid (13)

The mixture of tert-butyl 2,6-diazaspiro[3.3]heptane-2-carboxylate (400 mg, 2.0 mmol) and 5-bromo-2-fluoropyrimidine (300 mg, 1.7 mmol) in anhydrous DMF (15 mL) was added TEA (0.5 mL, 3.4 mmol) at room temperature . The mixture was stirred at 100 °C for 12 hours. After cooling to room temperature, the mixture was poured into 20 mL water and the compound was extracted with EtOAc. The organic phase was washed with brine and then dried over MgSO_4_, filtered, and concentrated under reduced pressure. The residue was purified by flash chromatography to afford compound 12 (500 mg, 84%). ^1^H NMR (400 MHz, CDCl_3_) δ 8.29 (s, 2H), 4.20 (s, 4H), 4.09 (s, 4H), 1.43 (s, 9H). MS (ESI): m/z 355.1, 357.1 [M+H]^+^.

A solution of **12** (100 mg, 0.3 mmol), (4-(methoxycarbonyl)phenyl)boronic acid (72 mg, 0.4 mmol), K_2_CO_3_ (83 mg, 0.6 mmol) and Pd(dppf)_2_Cl_2_.CH_2_Cl_2_ (25 mg, 0.03 mmol) in 1,4-dioxane (10 mL) and water (1 mL) was degassed and fulfilled with N_2_ for three times. Then the mixture was stirred at 100 °C for 12 hours before it was quenched with water. The resulting mixture was extracted with EtOAc, the combined organic phases were dried over anhydrous Na_2_SO_4_ and concentrated under reduced pressure. The residue was purified by silica gel flash column chromatography to give the product **13**. ^1^H NMR (400 MHz, DMSO-*d*_*6*_) δ 8.73 (s, 2H), 7.94 (d, *J* = 6.8 Hz, 2H), 7.74 (d, *J* = 8.3 Hz, 2H), 4.19 (s, 4H), 4.01 (s, 4H), 1.34 (s, 9H). MS (ESI): m/z 396.2 [M+H]^+^.

### General Procedures for the preparation of compound (14)

To a solution of **2** (50 mg, 0.1 mmol) and carboxylic acid (0.12 mmol) in DMF (5 mL) was added HATU (57 mg, 0.15 mmol) and DIPEA (86 µL, 0.6 mmol). The mixture was stirred at room temperature for 1 hour. The reaction was concentrated and then was purified by silica gel flash column chromatography to give the desired compound.

The compound was then dissolved in DCM (2 mL) and was added TFA (0.5 mL). The mixture was stirred at room temperature for 5 hours before it was concentrated under reduced pressure. Crude product **14** was used directly in the next step.

*N-(3-{4-amino-5-[1-(2-{3-[(trifluoromethyl)oxy]phenyl}acetyl)-2,3-dihydro-1H-indol-5-yl]pyrrolo[2,3-d]pyrimidin-7-yl}propyl)-4-{[1-(hexahydropyridin-4-yl)hexahydropyridin-4-yl]ethynyl}benzamide (****14a***): MS (ESI): m/z 805.4 [M+H]^+^.

*N-(3-{4-amino-5-[1-(2-{3-[(trifluoromethyl)oxy]phenyl}acetyl)-2,3-dihydro-1H-indol-5-yl]pyrrolo[2,3-d]pyrimidin-7-yl}propyl)-4-(hexahydropyridin-4-ylethynyl)benzamide (****14b***): MS (ESI): m/z 722.3 [M+H]^+^.

*N-(3-{4-amino-5-[1-(2-{3-[(trifluoromethyl)oxy]phenyl}acetyl)-2,3-dihydro-1H-indol-5-yl]pyrrolo[2,3-d]pyrimidin-7-yl}propyl)-4-{[6-(hexahydropyridin-4-yl)pyridin-3-yl]ethynyl}benzamide (****14c***): MS (ESI): m/z 799.3 [M+H]^+^.

*N-(3-{4-amino-5-[1-(2-{3-[(trifluoromethyl)oxy]phenyl}acetyl)-2,3-dihydro-1H-indol-5-yl]pyrrolo[2,3-d]pyrimidin-7-yl}propyl)-4-{[6-(2,6-diazaspiro[3*.*3]heptan-2-yl)pyridin-3-yl]ethynyl}benzamide (****14d***): MS (ESI): m/z 812.3 [M+H]^+^.

*4-[(5-aminopyridin-2-yl)ethynyl]-N-(3-{4-amino-5-[1-(2-{3-[(trifluoromethyl)oxy]phenyl}acetyl)-2,3-dihydro-1H-indol-5-yl]pyrrolo[2,3-d]pyrimidin-7-yl}propyl)benzamide (****14e***): MS (ESI): m/z 731.2 [M+H]^+^.

### General Procedures for the preparation of compound (15)

To a solution of **14** (0.04 mmol) and 2-(4-(4-((2,6-dioxopiperidin-3-yl)amino)phenyl)piperidin-1-yl)acetic acid (18 mg, 0.05 mmol) in DMF (3 mL) was added HATU (28 mg, 0.07 mmol) and DIPEA (34 µL, 0.24 mmol). The mixture was stirred at room temperature for 1 hour. The reaction was concentrated and then was purified by reverse phase preparative HPLC to provide the title compound **15**.

*N-(3-(4-amino-5-(1-(2-(3-(trifluoromethoxy)phenyl)acetyl)indolin-5-yl)-7H-pyrrolo[2,3-d]pyrimidin-7-yl)propyl)-4-((1’-(2-(4-(4-((2,6-dioxopiperidin-3-yl)amino)phenyl)piperidin-1-yl)acetyl)-[1,4’-bipiperidin]-4-yl)ethynyl)benzamide (****15a***): ^1^H NMR (400 MHz, DMSO-*d*_*6*_) δ 10.74 (s, 1H), 8.55 (s, 1H), 8.23 (s, 2H), 8.11 – 8.06 (m, 2H), 7.77 (d, *J* = 7.5 Hz, 2H), 7.41 (d, *J* = 6.6 Hz, 3H), 7.32 (s, 1H), 7.27 (s, 2H), 7.23 – 7.17 (m, 2H), 6.90 (d, *J* = 7.6 Hz, 2H), 6.56 (d, *J* = 7.5 Hz, 2H), 6.03 (brs, 1H), 5.60 (s, 1H), 4.34 (d, *J* = 10.9 Hz, 2H), 4.19 (d, *J* = 7.1 Hz, 8H), 4.08 (d, *J* = 11.3 Hz, 2H), 3.93 (s, 2H), 3.21 (d, *J* = 8.7 Hz, 6H), 3.00 (d, *J* = 13.1 Hz, 1H), 2.85 (s, 2H), 2.71 (d, *J* = 12.6 Hz, 2H), 2.31 (s, 4H), 2.01 (d, *J* = 7.1 Hz, 4H), 1.82 (s, 4H), 1.70 (m, 4H), 1.54 (d, *J* = 10.4 Hz, 4H). ^13^C NMR (100 MHz, DMSO-*d*_*6*_) δ 174.1, 173.5, 168.8, 167.6, 165.9, 157.6, 151.9, 150.4, 146.4, 142.1, 138.4, 134.3, 134.0, 130.5 (q, *J* = 211 Hz), 130.4, 127.8, 127.6, 126.1, 125.3, 123.6, 122.6, 119.4, 116.5, 115.4, 113.0, 100.3, 96.0, 61.7, 53.0, 48.2, 42.1, 41.6, 40.8, 37.2, 33.8, 32.2, 31.1, 30.1, 27.9, 25.1. Purity: >99.0%. MS (ESI): m/z 1132.5 [M+H]^+^. HRMS (ESI): m/z Calcd for (C_63_H_69_F_3_N_11_O_6_) ([M+H]^+^): 1132.5384; found: 1132.5378.

*N-(3-(4-amino-5-(1-(2-(3-(trifluoromethoxy)phenyl)acetyl)indolin-5-yl)-7H-pyrrolo[2,3-d]pyrimidin-7-yl)propyl)-4-((1-(2-(4-(4-((2,6-dioxopiperidin-3-yl)amino)phenyl)piperidin-1-yl)acetyl)piperidin-4-yl)ethynyl)benzamide (****15b***): ^1^H NMR (400 MHz, DMSO-*d*_*6*_) δ 10.78 (s, 1H), 8.61 (s, 1H), 8.32 (s, 1H), 8.17 – 8.08 (m, 2H), 7.82 (d, *J* = 7.4 Hz, 2H), 7.48 (t, *J* = 7.7 Hz, 3H), 7.34 (d, *J* = 20.0 Hz, 4H), 7.25 (dd, *J* = 14.8, 8.3 Hz, 3H), 6.94 (d, *J* = 7.6 Hz, 2H), 6.60 (d, *J* = 7.4 Hz, 2H), 6.07 (brs, 1H), 5.64 (s, 1H), 4.23 (d, *J* = 6.3 Hz, 4H), 3.97 (s, 2H), 3.87 (s, 4H), 3.37 (s, 1H), 3.29 – 3.21 (m, 4H), 3.20 – 3.09 (m, 4H), 2.96 (s, 1H), 2.89 (d, *J* = 9.6 Hz, 2H), 2.73 (t, *J* = 12.4 Hz, 1H), 2.57 (m, 1H), 2.30 (d, *J* = 11.1 Hz, 1H), 2.09 (s, 1H), 2.05 (d, *J* = 9.5 Hz, 4H), 1.90 (s, 1H), 1.85 (d, *J* = 11.1 Hz, 2H), 1.75 – 1.60 (m, 4H), 1.50 (dd, *J* = 48.6, 36.0 Hz, 4H). ^13^C NMR (100 MHz, DMSO-*d*_*6*_) δ 174.1, 173.5, 168.9, 167.9, 165.9, 157.6, 151.9, 150.5, 148.6, 146.4, 142.1, 138.4, 134.4, 134.1, 130.5 (q, *J* = 187 Hz), 130.4, 130.3, 127.8, 127.3, 125.9, 125.3, 123.6, 122.6, 121.8, 119.4, 116.5, 115.4, 113.0, 100.3, 95.2, 81.3, 61.8, 54.2, 53.0, 48.2, 44.4, 42.2, 41.7, 40.9, 37.2, 33.9, 32.3, 31.6, 31.1, 30.1, 27.9, 27.4, 25.1. Purity: >99.0%. MS (ESI): m/z 1049.5 [M+H]^+^. HRMS (ESI): m/z Calcd for (C_58_H_60_F_3_N_10_O_6_) ([M+H]^+^): 1049.4649; found: 1049.4659.

*N-(3-(4-amino-5-(1-(2-(3-(trifluoromethoxy)phenyl)acetyl)indolin-5-yl)-7H-pyrrolo[2,3-d]pyrimidin-7-yl)propyl)-4-((6-(4-(2-(4-(4-((2,6-dioxopiperidin-3-yl)amino)phenyl)piperidin-1-yl)acetyl)piperazin-1-yl)pyridin-3-yl)ethynyl)benzamide (****15c***): ^1^H NMR (400 MHz, DMSO-*d*_*6*_) δ 10.78 (s, 1H), 8.64 (s, 1H), 8.49 (s, 1H), 8.34 (s, 1H), 8.14 (s, 1H), 8.11 (d, *J* = 8.2 Hz, 1H), 7.86 (d, *J* = 7.9 Hz, 2H), 7.71 (d, *J* = 8.8 Hz, 1H), 7.59 (d, *J* = 8.0 Hz, 2H), 7.48 (t, *J* = 7.9 Hz, 1H), 7.36 (d, *J* = 10.1 Hz, 2H), 7.31 (s, 2H), 7.28 – 7.20 (m, 2H), 6.94 (t, *J* = 9.6 Hz, 3H), 6.60 (d, *J* = 8.0 Hz, 2H), 6.08 (brs, 1H), 5.65 (d, *J* = 6.9 Hz, 1H), 4.24 (s, 4H), 3.97 (s, 2H), 3.68 (d, *J* = 14.3 Hz, 4H), 3.58 (d, *J* = 7.2 Hz, 4H), 3.25 (d, *J* = 9.8 Hz, 6H), 3.20 (s, 1H), 2.92 (d, *J* = 10.1 Hz, 2H), 2.78 – 2.66 (m, 1H), 2.59 (s, 1H), 2.34 (m, 1H), 2.13 – 2.01 (m, 5H), 1.84 (d, *J* = 12.1 Hz, 1H), 1.69 (d, *J* = 11.1 Hz, 2H), 1.56 (d, *J* = 11.2 Hz, 2H). ^13^C NMR (100 MHz, DMSO-*d*_*6*_) δ 174.1, 173.5, 168.9, 168.3, 165.9, 158.0, 157.6, 151.9, 151.2, 150.5, 148.6, 146.4, 142.1, 140.4, 138.4, 134.4, 134.2, 130.5 (q, *J* = 184 Hz), 130.4, 130.2, 127.9, 127.3, 125.8, 125.3, 122.6, 119.4, 116.5, 115.4, 113.0, 107.3, 107.1, 100.3, 90.2, 89.7, 61.7, 54.1, 53.0, 48.2, 45.1, 44.4, 41.6, 40.8, 37.2, 33.9, 31.1, 30.1, 27.9, 25.1. Purity: >99.0%. MS (ESI): m/z 1127.5 [M+H]^+^. HRMS (ESI): m/z Calcd for (C_62_H_62_F_3_N_12_O_6_) ([M+H]^+^): 1127.4867; found: 1127.4915.

*N-(3-(4-amino-5-(1-(2-(3-(trifluoromethoxy)phenyl)acetyl)indolin-5-yl)-7H-pyrrolo[2,3-d]pyrimidin-7-yl)propyl)-4-((6-(6-(2-(4-(4-((2,6-dioxopiperidin-3-yl)amino)phenyl)piperidin-1-yl)acetyl)-2,6-diazaspiro[3*.*3]heptan-2-yl)pyridin-3-yl)ethynyl)benzamide (****15d***): ^1^H NMR (400 MHz, DMSO-*d*_*6*_) δ 10.74 (s, 1H), 8.58 (s, 1H), 8.25 (s, 1H), 8.13 – 8.04 (m, 2H), 7.82 (d, *J* = 7.6 Hz, 2H), 7.63 (d, *J* = 8.4 Hz, 1H), 7.54 (d, *J* = 7.5 Hz, 2H), 7.44 (t, *J* = 7.6 Hz, 1H), 7.33 (s, 1H), 7.29 (d, *J* = 13.6 Hz, 3H), 7.25 – 7.17 (m, 2H), 6.92 (d, *J* = 7.4 Hz, 2H), 6.57 (d, *J* = 7.5 Hz, 2H), 6.38 (d, *J* = 8.4 Hz, 1H), 6.03 (brs, 1H), 5.61 (d, *J* = 7.0 Hz, 1H), 4.39 (s, 2H), 4.20 (s, 6H), 4.13 (s, 4H), 4.04 (s, 2H), 3.93 (s, 2H), 3.21 (dd, *J* = 18.2, 7.6 Hz, 4H), 3.13 (s, 1H), 2.94 (s, 2H), 2.84 (s, 2H), 2.70 (t, *J* = 12.4 Hz, 1H), 2.53 (m, 1H), 2.27 (d, *J* = 12.6 Hz, 1H), 2.02 (d, *J* = 7.8 Hz, 3H), 1.81 (d, *J* = 10.6 Hz, 1H), 1.66 – 1.49 (m, 4H). ^13^C NMR (100 MHz, DMSO-*d*_*6*_) δ 174.1, 173.5, 169.7, 168.9, 165.9, 159.1, 157.6, 151.9, 151.5, 150.5, 148.6, 146.4, 142.1, 139.8, 138.4, 134.5, 134.1, 130.5 (q, *J* = 180 Hz), 130.4, 130.3, 127.4, 125.8, 125.3, 123.7, 122.6, 119.4, 116.5, 115.4, 113.0, 107.1, 106.0, 100.3, 90.4, 89.7, 61.1, 60.6, 60.0, 58.2, 54.3, 53.0, 48.2, 42.2, 41.6, 41.0, 37.2, 33.8, 33.7, 31.1, 30.1, 27.9, 25.1. Purity: >99.0%. MS (ESI): m/z 1139.5 [M+H]^+^. HRMS (ESI): m/z Calcd for (C_63_H_62_F_3_N_12_O_6_) ([M+H]^+^): 1139.4867; found: 1139.4851.

*N-(3-(4-amino-5-(1-(2-(3-(trifluoromethoxy)phenyl)acetyl)indolin-5-yl)-7H-pyrrolo[2,3-d]pyrimidin-7-yl)propyl)-4-((5-(2-(4-(4-((2,6-dioxopiperidin-3-yl)amino)phenyl)piperidin-1-yl)acetamido)pyridin-2-yl)ethynyl)benzamide (****15e***): ^1^H NMR (400 MHz, DMSO-*d*_*6*_) δ 10.71 (s, 1H), 10.12 (s, 1H), 8.85 (s, 1H), 8.62 (s, 1H), 8.18 (d, *J* = 8.5 Hz, 1H), 8.11 (s, 1H), 8.08 (d, *J* = 8.0 Hz, 1H), 7.86 (d, *J* = 8.1 Hz, 2H), 7.67 – 7.59 (m, 3H), 7.43 (d, *J* = 7.8 Hz, 1H), 7.33 (s, 1H), 7.28 (s, 3H), 7.25 – 7.16 (m, 3H), 6.95 (d, *J* = 8.0 Hz, 2H), 6.59 (d, *J* = 8.2 Hz, 2H), 6.05 (brs, 1H), 5.63 (d, *J* = 7.4 Hz, 1H), 4.20 (d, *J* = 7.2 Hz, 6H), 3.93 (s, 2H), 3.23 (d, *J* = 8.3 Hz, 2H), 3.14 (d, *J* = 11.9 Hz, 2H), 2.92 (d, *J* = 9.8 Hz, 2H), 2.71 (t, *J* = 12.2 Hz, 1H), 2.54 (m, 1H), 2.31 (s, 1H), 2.20 (t, *J* = 10.6 Hz, 2H), 2.03 (d, *J* = 6.7 Hz, 3H), 1.89 – 1.79 (m, 1H), 1.69 (d, *J* = 22.0 Hz, 4H). ^13^C NMR (100 MHz, DMSO-*d*_*6*_) δ 174.2, 173.5, 170.0, 168.9, 165.8, 157.6, 151.9, 150.5, 148.6, 146.4, 142.2, 142.0, 138.4, 136.5, 135.6, 135.0, 134.5, 131.8, 130.5 (q, *J* = 202 Hz), 130.4, 130.3, 128.0, 127.6, 126.8, 125.3, 124.7, 123.7, 122.7, 119.4, 116.6, 115.5, 113.1, 100.3, 91.2, 87.4, 62.6, 54.5, 53.0, 48.3, 42.2, 41.6, 40.8, 37.3, 33.6, 31.2, 30.1, 27.9, 25.1. Purity: 95.4%. MS (ESI): m/z 1058.4 [M+H]^+^. HRMS (ESI): m/z Calcd for (C_58_H_55_F_3_N_11_O_6_) ([M+H]^+^): 1058.4289; found: 1058.4269.

### General Procedures for the preparation of compound (17)

To a solution of **16** (50 mg, 0.1 mmol) and carboxylic acid (0.12 mmol) in DMF (5 mL) was added HATU (57 mg, 0.15 mmol) and DIPEA (86 µL, 0.6 mmol). The mixture was stirred at room temperature for 1 hour. The reaction was concentrated and then was purified by silica gel flash column chromatography to give the desired compound.

The compound was then dissolved in DCM (2 mL) and was added TFA (0.5 mL). The mixture was stirred at room temperature for 5 hours before it was concentrated under reduced pressure. Crude product **17** was used directly in the next step.

1-*[5-(4-amino-7-{1-[(4-{[6-(2,6-diazaspiro[3*.*3]heptan-2-yl)pyridin-3-yl]ethynyl}phenyl)carbonyl]hexahydropyridin-4-yl}pyrrolo[2,3-d]pyrimidin-5-yl)-2,3-dihydro-1H-indol-1-yl]-2-{3-[(trifluoromethyl)oxy]phenyl}ethan-1-one (****17a***): MS (ESI): m/z 838.3 [M+H]^+^.

2-*1-(5-{4-amino-7-[1-({4-[2-(2,6-diazaspiro[3*.*3]heptan-2-yl)pyrimidin-5-yl]phenyl}carbonyl)hexahydropyridin-4-yl]pyrrolo[2,3-d]pyrimidin-5-yl}-2,3-dihydro-1H-indol-1-yl)-2-{3-[(trifluoromethyl)oxy]phenyl}ethan-1-one (****17b***): MS (ESI): m/z 815.3 [M+H]^+^.

3-*1-(5-{4-amino-7-[1-({4-[(5-aminopyridin-2-yl)ethynyl]phenyl}carbonyl)hexahydropyridin-4-yl]pyrrolo[2,3-d]pyrimidin-5-yl}-2,3-dihydro-1H-indol-1-yl)-2-{3-[(trifluoromethyl)oxy]phenyl}ethan-1-one (****17c***): MS (ESI): m/z 757.3 [M+H]^+^.

*1-{5-[4-amino-7-(1-{[4-(piperazin-1-yl)phenyl]carbonyl}hexahydropyridin-4-yl)pyrrolo[2,3-d]pyrimidin-5-yl]-2,3-dihydro-1H-indol-1-yl}-2-{3-[(trifluoromethyl)oxy]phenyl}ethan-1-one (****17d***): MS (ESI): m/z 725.3 [M+H]^+^.

Compound **18** was prepared according to the procedure described above for compound **15**.

*3-((4-(1-(2-(6-(5-((4-(4-(4-amino-5-(1-(2-(3-(trifluoromethoxy)phenyl)acetyl)indolin-5-yl)-7H-pyrrolo[2,3-d]pyrimidin-7-yl)piperidine-1-carbonyl)phenyl)ethynyl)pyridin-2-yl)-2,6-diazaspiro[3*.*3]heptan-2-yl)-2-oxoethyl)piperidin-4-yl)phenyl)amino)piperidine-2,6-dione (****LD5095*** (***18a***)). ^1^H NMR (400 MHz, DMSO-*d*_*6*_) δ 10.78 (s, 1H), 8.28 (s, 1H), 8.12 (d, *J* = 12.0 Hz, 2H), 7.66 (d, *J* = 8.6 Hz, 1H), 7.60 – 7.53 (m, 3H), 7.48 (dd, *J* = 10.9, 8.1 Hz, 3H), 7.33 (d, *J* = 12.7 Hz, 3H), 7.26 (s, 2H), 6.96 (d, *J* = 7.8 Hz, 2H), 6.60 (d, *J* = 8.0 Hz, 2H), 6.41 (d, *J* = 8.7 Hz, 1H), 6.10 (s, 1H), 5.65 (d, *J* = 7.1 Hz, 1H), 4.93 (s, 1H), 4.65 (s, 1H), 4.43 (m, 2H), 4.24 (d, *J* = 7.6 Hz, 3H), 4.16 (m, 4H), 4.07 (m, 3H), 3.97 (s, 2H), 3.69 (s, 1H), 3.24 (t, *J* = 7.8 Hz, 2H), 2.97 (s, 2H), 2.88 (d, *J* = 9.0 Hz, 2H), 2.73 (t, *J* = 12.2 Hz, 1H), 2.57 (m, 1H), 2.29 (m, 1H), 2.14 – 1.99 (m, 6H), 1.85 (d, *J* = 9.4 Hz, 2H), 1.70 – 1.51 (m, 4H). ^13^C NMR (100 MHz, DMSO-*d*_*6*_) δ 174.1, 173.5, 169.7, 168.9, 168.8, 159.1, 157.6, 151.8, 515.5, 150.1, 148.6, 146.4, 142.1, 139.8, 138.4, 136.1, 134.5, 130.5 (q, *J* = 196 Hz), 130.3, 127.6, 127.4, 125.3, 124.3, 122.6, 120.9, 119.4, 116.5, 115.9, 113.1, 107.2, 106.0, 100.3, 89.6, 61.1, 60.6, 60.0, 58.2, 54.3, 53.0, 51.1, 48.3, 41.6, 40.9, 33.8, 33.7, 31.1, 27.9, 25.1. Purity: >99.0%. MS (ESI): m/z 1165.5 [M+H]^+^. HRMS (ESI): m/z Calcd for (C_65_H_64_F_3_N_12_O_6_) ([M+H]^+^): 1165.5024; found: 1165.4994.

*3-((4-(1-(2-(6-(5-(4-(4-(4-amino-5-(1-(2-(3-(trifluoromethoxy)phenyl)acetyl)indolin-5-yl)-7H-pyrrolo[2,3-]pyrimidin-7-yl)piperidine-1-carbonyl)phenyl)pyrimidin-2-yl)-2,6-diazaspiro[3*.*3]heptan-2-yl)-2-oxoethyl)piperidin-4-yl)phenyl)amino)piperidine-2,6-dione (****18b***): ^1^H NMR (400 MHz, DMSO-*d*_*6*_) δ 8.71 (s, 2H), 8.09 (d, *J* = 11.8 Hz, 2H), 7.69 (d, *J* = 7.8 Hz, 2H), 7.51 (d, *J* = 6.1 Hz, 2H), 7.44 (d, *J* = 7.7 Hz, 1H), 7.30 (d, *J* = 12.9 Hz, 4H), 7.23 (s, 2H), 6.92 (d, *J* = 7.3 Hz, 2H), 6.57 (d, *J* = 8.2 Hz, 1H), 6.51 – 6.39 (m, 1H), 6.06 (brs, 1H), 5.61 (d, *J* = 7.1 Hz, 1H), 4.89 (s, 1H), 4.41 (s, 2H), 4.22 (s, 8H), 4.05 (s, 2H), 3.94 (s, 2H), 3.20 (t, *J* = 8.0 Hz, 2H), 2.94 (s, 2H), 2.84 (s, 2H), 2.67 (m, 1H), 2.56 (s, 1H), 2.26 (s, 1H), 2.17 (t, *J* = 7.0 Hz, 1H), 2.03 (s, 5H), 1.88 – 1.77 (m, 2H), 1.61 (s, 4H). ^13^C NMR (100 MHz, DMSO-*d*_*6*_) δ 174.2, 173.8, 173.5, 169.7, 169.1, 168.9, 162.1, 157.6, 156.3, 151.7, 150.1, 148.6, 146.4, 142.1, 138.4, 136.5, 135.2, 134.5, 130.5 (q, *J* = 192 Hz), 130.3, 128.0, 127.4, 125.8, 125.3, 122.6, 122.2, 120.9, 119.4, 116.5, 115.9, 113.0, 112.6, 100.3, 61.1, 60.4, 60.0, 58.2, 54.3, 53.0, 51.1, 48.2, 41.6, 41.0, 33.8, 33.6, 31.6, 31.1, 27.9, 25.2. Purity: 98.6%. MS (ESI): m/z 1142.5 [M+H]^+^. HRMS (ESI): m/z Calcd for (C_62_H_63_F_3_N_13_O_6_) ([M+H]^+^): 1142.4976; found: 1142.4993.

*N-(6-((4-(4-(4-amino-5-(1-(2-(3-(trifluoromethoxy)phenyl)acetyl)indolin-5-yl)-7H-pyrrolo[2,3-d]pyrimidin-7-yl)piperidine-1-carbonyl)phenyl)ethynyl)pyridin-3-yl)-2-(4-(4-((2,6-dioxopiperidin-3-yl)amino)phenyl)piperidin-1-yl)acetamide (****18c***): ^1^H NMR (400 MHz, DMSO-*d*_*6*_) δ 10.79 (s, 1H), 10.17 (s, 1H), 8.88 (s, 1H), 8.27 (s, 1H), 8.21 (d, *J* = 8.5 Hz, 1H), 8.13 (d, *J* = 12.1 Hz, 2H), 7.66 (t, *J* = 10.6 Hz, 3H), 7.55 (d, *J* = 5.7 Hz, 3H), 7.48 (t, *J* = 7.8 Hz, 1H), 7.35 (s, 1H), 7.32 (s, 1H), 7.27 (s, 2H), 6.99 (d, *J* = 7.8 Hz, 2H), 6.62 (d, *J* = 7.9 Hz, 2H), 6.10 (brs, 1H), 5.66 (s, 1H), 4.94 (s, 1H), 4.66 (s, 2H), 4.25 (t, *J* = 7.6 Hz, 4H), 3.97 (s, 2H), 3.24 (t, *J* = 8.2 Hz, 2H), 3.18 (d, *J* = 10.7 Hz, 2H), 2.96 (d, *J* = 9.8 Hz, 2H), 2.74 (t, *J* = 12.1 Hz, 1H), 2.58 (m, 1H), 2.35 (s, 1H), 2.24 (t, *J* = 10.4 Hz, 2H), 2.08 (s, 4H), 1.86 (d, *J* = 8.7 Hz, 2H), 1.73 (m, 4H). ^13^C NMR (100 MHz, DMSO-*d*_*6*_) δ 174.2, 173.5, 170.3, 169.0, 168.6, 157.7, 151.8, 150.1, 148.6, 146.4, 142.2, 142.0, 138.4, 137.0, 136.6, 135.6, 134.5, 132.0, 130.5 (q, *J* = 192 Hz), 130.4, 127.9, 127.7, 126.8, 125.3, 123.1, 122.7, 119.4, 116.6, 116.0, 113.1, 100.3, 90.5, 87.4, 62.6, 54.5, 53.0, 51.1, 48.3, 41.6, 40.9, 33.6, 31.1, 27.9, 25.2. Purity: 98.5%. MS (ESI): m/z 1084.4 [M+H]^+^. HRMS (ESI): m/z Calcd for (C_60_H_57_F_3_N_11_O_6_) ([M+H]^+^): 1084.4445; found: 1084.4436.

*3-((4-(1-(2-(4-(4-(4-(4-amino-5-(1-(2-(3-(trifluoromethoxy)phenyl)acetyl)indolin-5-yl)-7H-pyrrolo[2,3-d]pyrimidin-7-yl)piperidine-1-carbonyl)phenyl)piperidin-1-yl)-2-oxoethyl)piperidin-4-yl)phenyl)amino)piperidine-2,6-dione (****18d***): ^1^H NMR (400 MHz, DMSO-*d*_*6*_) δ 10.78 (s, 1H), 8.24 (s, 1H), 8.12 (d, *J* = 12.5 Hz, 2H), 7.53 (s, 1H), 7.48 (t, *J* = 7.7 Hz, 1H), 7.35 (t, *J* = 12.2 Hz, 5H), 7.26 (s, 2H), 7.00 (d, *J* = 7.6 Hz, 2H), 6.94 (d, *J* = 7.4 Hz, 2H), 6.60 (d, *J* = 7.3 Hz, 2H), 6.09 (brs, 1H), 5.64 (s, 1H), 4.90 (s, 1H), 4.24 (s, 4H), 3.97 (s, 2H), 3.73 (s, 2H), 3.60 (s, 2H), 3.22 (m, 10H), 2.92 (d, *J* = 8.9 Hz, 2H), 2.78 – 2.65 (m, 1H), 2.56 (m, 1H), 2.32 (s, 1H), 2.08 (d, *J* = 10.7 Hz, 5H), 1.93 (s, 2H), 1.84 (d, *J* = 11.9 Hz, 1H), 1.69 (d, *J* = 10.6 Hz, 2H), 1.56 (d, *J* = 11.6 Hz, 2H). ^13^C NMR (100 MHz, DMSO-*d*_*6*_) δ 174.1, 173.5, 169.7, 168.9, 168.0, 157.6, 152.0, 151.7, 150.1, 148.6, 146.4, 142.1, 138.4, 134.4, 130.5 (q, *J* = 190 Hz), 130.4, 129.0, 127.3, 126.0, 125.3, 122.6, 120.9, 119.4, 116.5, 115.8, 114.7, 113.0, 61.6, 54.1, 53.0, 51.4, 48.5, 48.3, 47.8, 45.2, 41.7, 41.3, 40.8, 33.8, 32.2, 31.1, 27.9, 25.1. Purity: 99.0%. MS (ESI): m/z 1052.5 [M+H]^+^. HRMS (ESI): m/z Calcd for (C_57_H_61_F_3_N_11_O_6_) ([M+H]^+^): 1052.4758; found: 1052.4739.

*3-((4-(1-(2-(6-(5-((4-(4-(4-amino-5-(1-(2-(3-(trifluoromethoxy)phenyl)acetyl)indolin-5-yl)-7H-pyrrolo[2,3-d]pyrimidin-7-yl)piperidine-1-carbonyl)phenyl)ethynyl)pyridin-2-yl)-2,6-diazaspiro[3*.*3]heptan-2-yl)-2-oxoethyl)piperidin-4-yl)phenyl)amino)-1-methylpiperidine-2,6-dione (****LD5095-NC***): ^1^H NMR (400 MHz, DMSO-*d*_*6*_) δ 8.53 (s, 1H), 8.36 (s, 1H), 8.12 (d, *J* = 11.2 Hz, 2H), 7.74 (d, *J* = 8.5 Hz, 1H), 7.59 (d, *J* = 7.3 Hz, 2H), 7.55 – 7.46 (m, 4H), 7.33 (d, *J* = 13.1 Hz, 3H), 7.26 (s, 2H), 7.08 (d, *J* = 7.3 Hz, 2H), 6.94 (d, *J* = 8.3 Hz, 1H), 6.64 (d, *J* = 7.6 Hz, 2H), 6.09 (brs, 1H), 5.76 (d, *J* = 6.8 Hz, 1H), 4.93 (m, 1H), 4.60 (s, 2H), 4.25 (m, 3H), 4.12 (d, *J* = 10.1 Hz, 1H), 3.97 (s, 2H), 3.83 (d, *J* = 10.5 Hz, 2H), 3.73 (s, 2H), 3.65 (m, 4H), 3.59 (s, 3H), 3.26 (d, *J* = 14.3 Hz, 2H), 3.17 (s, 2H), 3.03 – 2.85 (m, 1H), 2.70 (d, *J* = 13.5 Hz, 2H), 2.57 (m, 1H), 2.07 (m, 8H), 1.89 (d, *J* = 13.1 Hz, 4H). ^13^C NMR (100 MHz, DMSO-*d*_*6*_) δ 174.2, 173.6, 169.0, 168.7, 166.4, 162.7, 157.8, 157.6, 151.8, 151.1, 150.1, 148.6, 147.0, 142.2, 140.5, 138.4, 136.2, 131.7, 130.5 (q, *J* = 201 Hz), 130.4, 127.7, 125.3, 124.2, 122.7, 121.8, 120.9, 119.4, 116.5, 115.9, 113.0, 107.6, 107.1, 100.3, 89.7, 89.3, 64.5, 61.7, 52.9, 51.1, 48.2, 44.9, 44.5, 44.1, 41.6, 41.2, 38.4, 31.2, 27.9, 27.4, 25.1. Purity: 98.8%. MS (ESI): m/z 1179.5 [M+H]^+^. HRMS (ESI): m/z Calcd for (C_66_H_66_F_3_N_12_O_6_) ([M+H]^+^): 1179.5180; found: 1179.5144.

### Degradation Evaluation of nLuc-RIPK1 in Jurkat cells

Jurkat cells expressing nLuc-RIPK1 were seeded into 96-well white plates (Corning) at 1□×□10^4^ cells/well in 99□μL of DMEM/FCS, followed by overnight incubation. Cells were then treated with either DMSO or compound as indicated. After 24 hours, 100 uL of the Nano-Glo Luciferase assay system (Promega) was added into each well. Plates were incubated for 10□min at room temperature before assaying for luminescence. Luminescence was then measured on a BioTek Synergy H1 plate.

### Metabolic Stability in Mouse Liver S9 Fraction

Mix the 25.5 uL of MgCl_2_ (50 mM), 15 uL of S9 fraction (20 mg/ml) and 186.5 uL PSB with 25.5 uL of NADPH (10 mM) or 25.5 μL of ultrapure water. The final concentration of S9 fraction was 1 mg/mL. The mixture was pre-warmed at 37°C for 10 minutes.

The reaction was started with the addition of 2.55 μL of 100 μM control compound or test compound solutions. Verapamil was used as positive control in this study. The final concentration of control compound and test compounds were 1 μM. The incubation solution was incubated in water batch at 37°C.

Aliquots of 25 µL were taken from the reaction solution at 0, 15, 30, 45, 60 and 90 minutes. The reaction was stopped by the addition of 10 volumes of cold acetonitrile with IS (100 nM imipramine). Samples were centrifuged at 12,000 g for 10 minutes. Aliquot of 100 µL of the supernatant was mixed with 100 µL of ultra-pure water and then used for LC-MS/MS analysis. All calculations were carried out using GraphPad Prism 10.0.

### Western Blotting

Cells were seeded into six-well plates at a density of 5×10^5^ cells/mL in 2 mL of complete culture medium. Following an overnight adaptation period, cells were treated with serially diluted compounds for 24 h. After treatment, whole-cell lysates were prepared using a lysis buffer (1×RIPA supplemented with protease and phosphatase inhibitor cocktail). Protein concentrations in the lysates were measured using the BCA protein assay. Subsequently, equal amounts of protein (20 µg) from each sample were loaded onto a sodium dodecyl sulfate-polyacrylamide gel and separated by electrophoresis (Bio-Rad) at 120 V for 1.5 hours. The separated proteins were then transferred to a polyvinylidene fluoride (PVDF) membrane using a Transblot Turbo system (Bio-Rad). After blocking for 2 h at room temperature in 5% BSA-TBST, the membranes were incubated overnight at 4°C with specific primary antibodies (diluted at 1:1000 in TBST) targeting the proteins of interest, including anti-RIPK1 (3493, Cell Signaling Technology (CST)), anti-cleaved caspase 3 (9661, CST), anti-cleaved caspase7 (8438, CST), anti-cleaved PARP (5625,CST), and anti-β-actin (4970, CST). The membranes were then incubated with horseradish peroxidase-conjugated secondary antibodies (1:1000, 7074, CST) for 1 h at room temperature. Immunoblots were imaged using ECL Prime chemiluminescent western blot detection reagent (R1100, Kindle Biosciences,) and visualized using an Imager (D1001, Kindle Biosciences). All western blots were processed and quantified using ImageJ software, and protein levels were normalized to β-actin loading controls.

### Immunoprecipitation and Western blotting

Wild-type Jurkat cells were harvested at a density of 0.8 × 10□ cells/mL (10 mL; 8 × 10□ cells total) and, following the indicated pre-treatment of 1uM Carfilzomib 0.5h then 100nM LD5095 for 4h, collected by centrifugation (300 × g, 5 min, 4 °C) and washed once with ice-cold PBS. Cells were lysed in 500ul ice-cold lysis buffer containing 50 mM Tris-HCl (pH 7.5), 150 mM NaCl, 1% (v/v) NP-40, 0.1% (w/v) SDS, 1 mM EDTA, 10 mM N-ethylmaleimide, 10 µM PR-619, and 1× protease inhibitor cocktail. Lysates were incubated on ice for 10 min and clarified by centrifugation (16,000 × g, 15 min, 4 °C). Protein concentration was determined by BCA assay, and equal amounts of total protein were used across conditions; an aliquot of each lysate was retained as input.

Lysates were pre-cleared with 25 µL of Protein A magnetic beads (Pierce, Thermo Fisher Scientific) for 60 min at 4 °C. Cleared lysates were then incubated with anti-RIPK1 antibody (Cell Signaling Technology 3493; 1:100) overnight at 4 °C with rotation. Immune complexes were captured by the addition of 25 µL of Protein A magnetic beads for 1h at 4 °C. Beads were washed four times with lysis buffer, and bound proteins were eluted by boiling in 1× Laemmli sample buffer for 5 min.

Eluates and inputs were resolved by SDS-PAGE, transferred to PVDF membranes, and immunoblotted overnight for ubiquitin (Cell Signaling Technology #14049), RIPK1 (Cell Signaling Technology 3493; 1:1000) and beta-actin (Cell Signaling Technology 4970; 1:5000). Signals were detected by ECL substrates (Vazyme E423-01).

### Quantitative PCR (qPCR) Asaay

Wild-type Jurkat cells were seeded in 6-well plates at 0.2 × 10□ cells/mL (2 mL per well; 4 × 10□ cells per well) and treated with 100 nM LD5095 or an equivalent volume of DMSO (vehicle) for 24 h. Total RNA was extracted using the RNeasy Mini Kit (Qiagen) according to the manufacturer’s instructions, and RNA concentration was determined by NanoDrop. Equal amounts of total RNA were used as template across all samples. One-step quantitative RT-PCR was performed using SYBR Green one-step qPCR master mix (Vazyme, CL132) on a LightCycler® 480 Instrument II (Roche) in 384-well plates. The cycling program was: reverse transcription at 55 °C for 15 min; initial denaturation at 95 °C for 30 s; 40 cycles of 95 °C for 10 s and 60 °C for 30 s; followed by melt-curve analysis using the instrument’s default acquisition program to confirm amplification specificity. Data was reported as deltaCt with GAPDH as the reference gene. Data represent n = 3 biological replicates. The primers used were: RIPK1, 5′-TATCCCAGTGCCTGAGACCAAC-3′ (forward) and 5′-GTAGGCTCCAATCTGAATGCCAG-3′ (reverse); GAPDH, 5′-GGAGCGAGATCCCTCCAAAAT-3′ (forward) and 5′-GGCTGTTGTCATACTTCTCATGG-3′ (reverse).

### Cell Viability Assay

Indicated cells (7,500 cells per well) were seeded into 96-well plates containing 150 μL of RPMI medium supplemented with 10% FBS. On the following day, DMSO or drug was added. After 72 hrs, cell viability was measured by Alamar Blue assay. For statistical significance, six replications were tested in each treatment. Relative cell viabilities, referred to DMSO control cells, were plotted using Graphpad Prism.

### Apoptosis Detection Using FITC-conjugated Annexin V/PI

Apoptosis quantification was conducted utilizing a FITC-conjugated Annexin V/PI assay kit (556547, BD Biosciences) and analyzed through flow cytometry. Briefly, 2×10^5^ of Jurkat cells were seeded onto six-well plates and treated as specified for 72 hours at 37°C. Treated and untreated cells were harvested, washed with PBS, and resuspended in 100 µl of binding buffer. Subsequently, cells were stained with PI (50 µg/ml) and FITC-conjugated Annexin V (10 mg/ml) for 15 minutes at room temperature in the dark. After adding another 400 µl of binding buffer, the cells were subjected to LSR II Flow cytometer (BD Biosciences) for analysis, and flow cytometry data were processed using the FlowJo software.

### PK/PD Study in Mice

All *in vivo* studies were performed under an animal protocol (AN-6075) approved by the Institutional Animal Care and Use Committee (IACUC) of Baylor College of Medicine, in accordance with the recommendations in the Guide for the Care and Use of Laboratory Animals of the National Institutes of Health.

For IV injection, drug (Formulation: 0.4 mg/mL in cosolvent which content 10% of DMSO, 30% of PEG400, 5% of Tween80 in PBS; Dosing: 1 mg/kg) were injected into C57BL/6J male mice (n = 3). 10 µL of blood was collected via the tail vein at 2, 5, 15, 30 min, and 1, 2, 4, 8, 24 and 48 h.

The desired serial concentrations of working solutions were achieved by diluting stock solution of analyte with 100% acetonitrile solution. 5 µL of working solutions (1, 2, 4, 10, 20, 100, 200, 1000, 2000 ng/mL) were added to 10 μL of the blank C57BL/6J mouse plasma to achieve calibration standards of 0.5∼1000 ng/mL (0.5, 1, 2, 5, 10, 50, 100, 500, 1000 ng/mL) in a total volume of 15 μL. Four quality control samples at 1 ng/mL, 2 ng/mL, 50 ng/mL and 800 ng/mL for plasma were prepared independently of those used for the calibration curves. These QC samples were prepared on the day of analysis in the same way as calibration standards.

15 µL standards and 15 µL QC samples and 15 µL unknown samples (10 µL plasma with 5 µL blank solution) were added to 200 µL of acetonitrile containing IS mixture for precipitating protein respectively. Then the samples were vortexed for 30 s. After centrifugation at 4 °C, 3900 rpm for 15 min, the supernatant was diluted 3 times with MeCN. 5 µL of supernatant was injected into the LC/MS/MS system for quantitative analysis.

The pharmacokinetic parameters of drug were analyzed using PK Solver Excel® Add-in. The PK trace was fitted into a non-compartmental extravascular model.

To establish Jurkat xenograft tumors, 6-week-old female nude mice (The Jackson Laboratory) were subcutaneously injected with 2.5 ×10^6^ Jurkat cells suspended in a 1:1 mixture with Matrigel. Once tumors reached approximately 100 mm^2^ in size, mice (n = 4 per group) received a single intravenous dose of either the test compound (10 mg/kg) or vehicle control. At indicated time posttreatment, mice were euthanized, and tumor tissues were harvested for Western blot analysis.

## ASSOCIATED CONTENT

### Supporting Information

Supporting Information is available free of charge via the Internet at http://pubs.acs.org.

Fitting curve of tested compounds; Degradation potency of LD5095 in cell lines; Oral PK of LD5095; qPCR analysis; ubiquitination blotting; ^1^H and ^13^C NMR spectra; HPLC analysis (PDF)

Molecular formula strings (CSV).

Proteomics list (XLSX)

Research involving animals was performed in accordance with institutional guidelines as defined by Institutional Animal Care and Use Committee.

## AUTHOR INFORMATION

### Author Contributions

‡D.L. and X.Y. contributed equally.

### Notes

The authors declare the following competing financial interest(s): J.W. is the co-founder of Chemical Biology Probes LLC. J. W. has stock ownership in CoRegen Inc and serves as a consultant for this company. J.W., X.Y. and B.Y. are the co-founders of Fortitude Biomedicines, Inc. and hold equity interest in this company. X.Y., D.L., and J.W. are inventors on a patent covering RIPK1 degraders reported in this work, titled “Novel RIPK1 Kinase-Targeting PROTACs and Methods of Use Thereof”, with the identification number WO2022120118A1. The remaining authors declare no competing interests.

## ACKNOWLEDGMENT

The research was supported in part by National Institute of Health (R01-268518 to J.W.), Cancer Prevention & Research Institute of Texas (CPRIT, RP220480 to J.W.), the seed fund to Center for NextGen Therapeutics, and the Michael E. DeBakey, M.D., Professorship in Pharmacology (to J.W.). This project was supported by the Cytometry and Cell Sorting Core at Baylor College of Medicine with funding from the CPRIT Core Facility Support Award (CPRIT-RP240432) and the NIH (CA125123, OD036336, and OD038251), and the assistance of Joel M. Sederstrom.

## ABBREVIATIONS

AcOH: Acetic acid
CoCl_2_.6H_2_O: Cobalt chloride hexahydrate
CRBN: cereblon
CuI: Copper(I) iodide
DCE: 1,2-dichloroethane
DCM: dichloromethane
DIPEA: N,N-Diisopropylethylamine
DMF: N,N-Dimethylformamide
EtOAc: ethyl acetate
FA: formic acid
HATU: 2-(7-Aza-1H-benzotriazole-1-yl)-1,1,3,3-tetraMethyluroniuM hexafluorophosphate
IV: intravenous
K_2_CO_3_: Potassium carbonate
KI: Potassium iodide
LiOH: Lithium hydroxide
nLuc: NanoLuc luciferase
MeCN: Acetonitrile
MeOH: methanol
N_2_: nitrogen gas
NaBH(OAc)_3_: Sodium triacetoxyborohydride
NADPH: nicotinamide adenine dinucleotide phosphate
NaH: sodium hydride
NF-κB: Nuclear factor kappa-light-chain-enhancer of activated B cells
PD: pharmaco-dynamics
PD: pharmacodynamics
PD-1: programmed cell death protein 1
Pd(PPh_3_)_2_Cl_2_: Bis(triphenylphosphine)palladium(II) chloride
PK: pharmacokinetics
PROTAC: proteolysis targeting chimera
rt: room temperature
TEA: triethylamine
TFA: Trifluoroacetic acid
THF: Tetrahydrofuran
VHL: von Hippel-Lindau Cullin RING E3 ligase
WT: wild-type

---

Insert Table of Contents artwork here

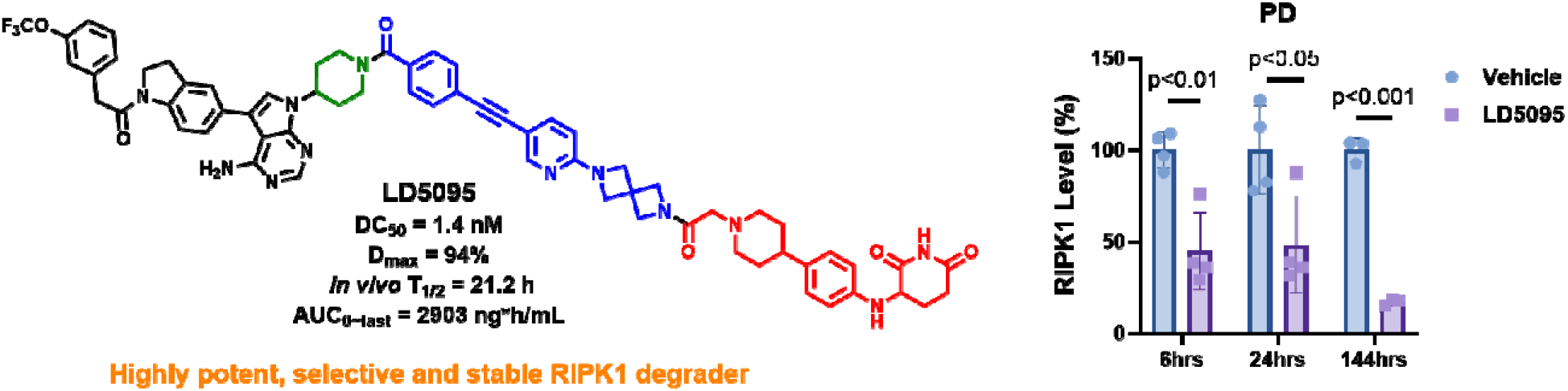

