## supporting info for "Discovery and Development of First-in-Class Cereblon-Recruiting RIPK1 Degraders"

### Supporting Information

#### Table of Contents

**Fitting curve of tested compounds for nLuc-RIPK1 degradation in Jurkat cells**...S2
**Plasma concentration-time curves of tested compounds in C57BL/6J mice** …........S3
**Degradation potency of LD5095 in MOLM14 and U937 cells** ……...………….….... S4
**Degradation potency of LD5095 in B16F10 cells** ……... …... …... …..……………....S4
**Impact of LD5095 on cell viability across tumor and primary Cell Lines**...............S4
**Plasma concentration-time profile of LD5095 following oral administration**.........S5
**Pharmacokinetic parameters of LD5095 in ICR mice**…..……………...…...….....S5
**Quantitative PCR (qPCR) analysis of RIPK1 in Jurkat cells**…..……………….........S5
**Effect of LD5095 on the ubiquitination of RIPK1 in Jurkat cells**…..………….........S6
**Pharmacokinetic Parameters of LD5095 in Mice via oral administration**…..........S7
**RIPK1 SILAC Half-Life Summary**…..……..............................................…..........S7
**Purity of target compounds**…………………………………………..…………....S8
**$^{1}$H NMR, $^{13}$C NMR spectral data of target compounds**……………..…….……..…S9
**HPLC chromatography of representative compounds**…………………….………......S49

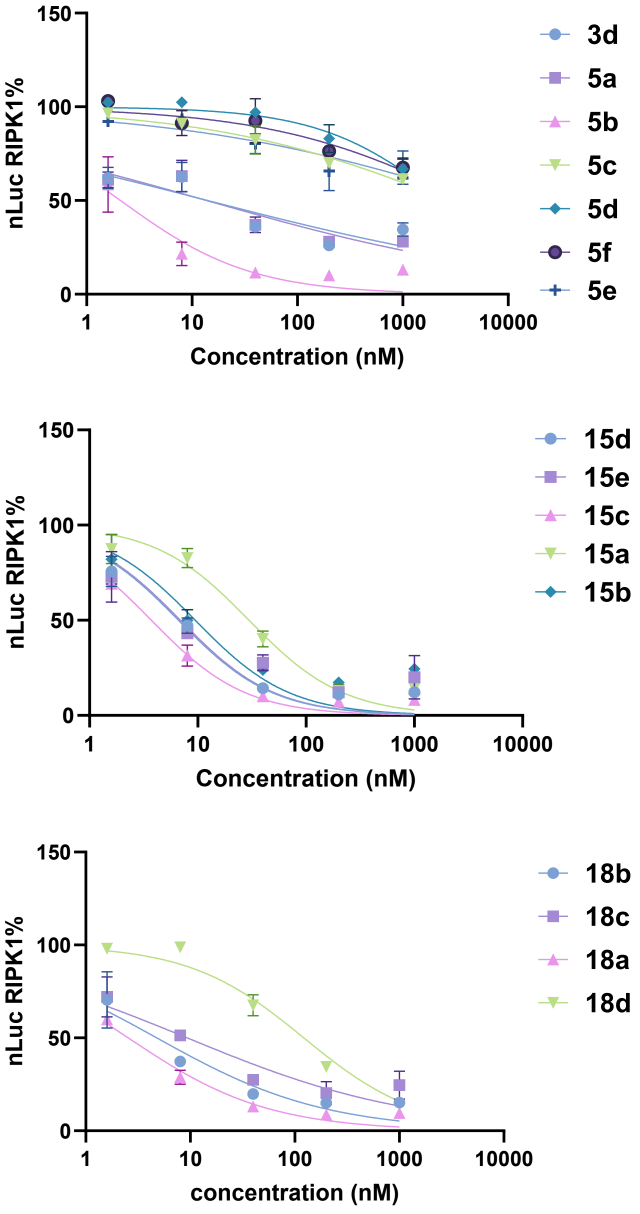


**Figure. S1.** Fitting curve of tested compounds for nLuc-RIPK1 degradation in Jurkat cells.

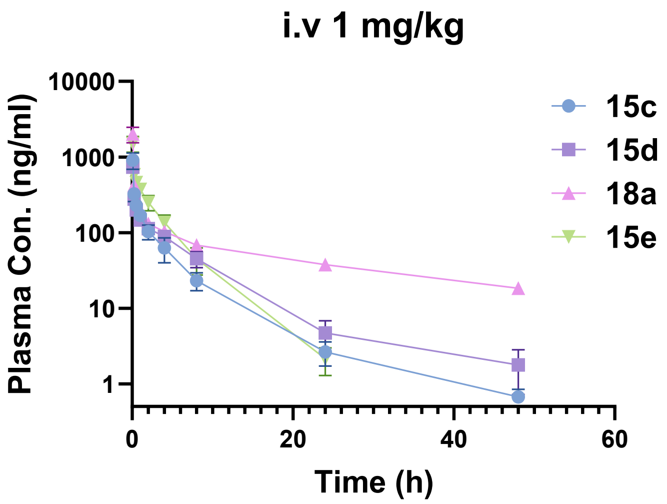


**Figure. S2.** Plasma concentration-time curves of tested compounds in mice.

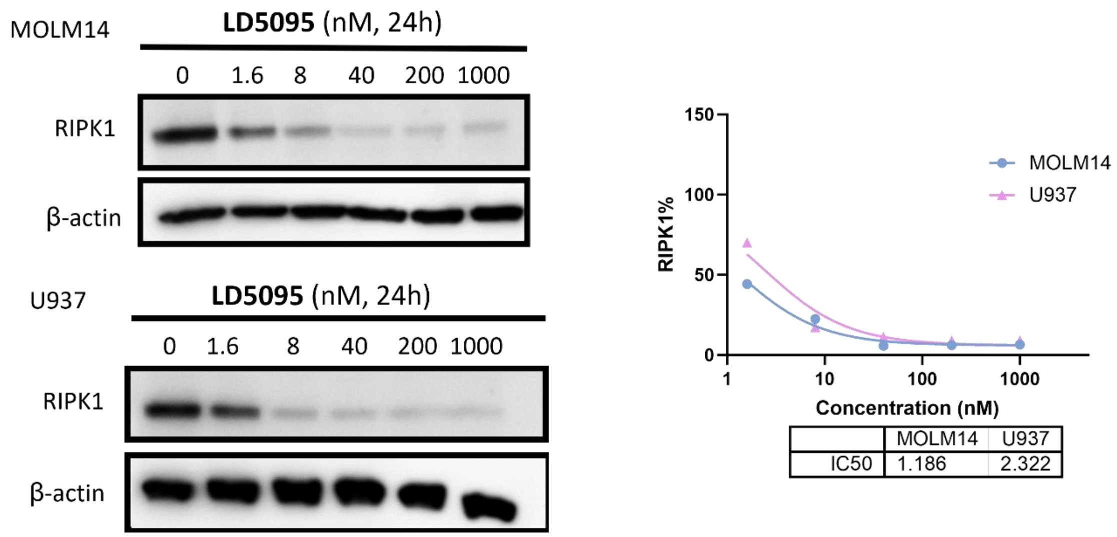


**Figure S3.** Degradation potency of **LD5095** (**18a**) in MOLM14 and U937 cells.

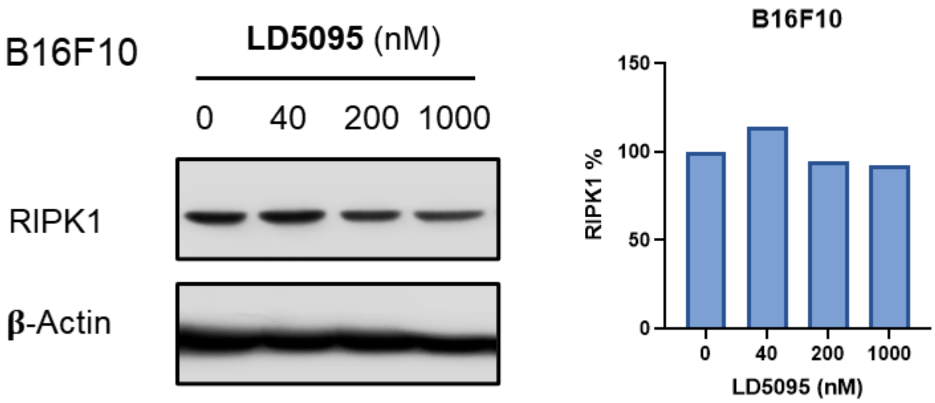


**Figure S4.** Degradation potency of **LD5095** (**18a**) in B16F10 cells

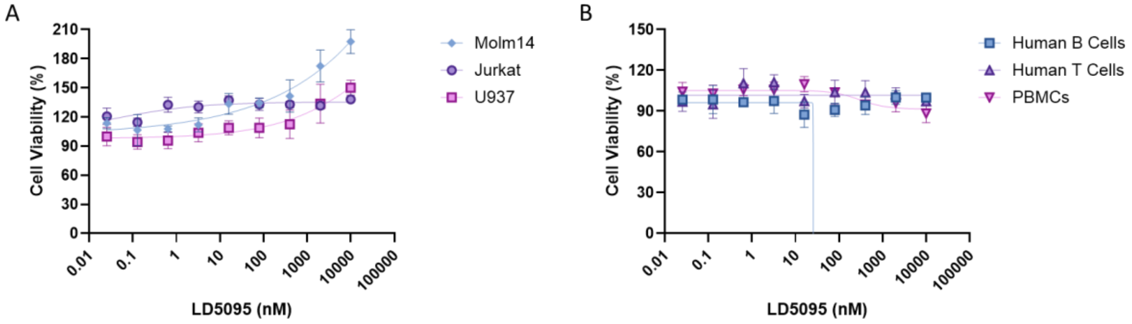


**Figure S5.** Impact of LD5095 on cell viability across tumor and primary cell lines. (A) Cell viability of indicated cancer cell lines following 72-hour treatment with LD5095. For each assay, 4,000

cancer cells were seeded per well in 96-well plates (n = 3 biological replicates). (B) Cell viability of human primary cells following 72-hour treatment with LD5095. For these assays, 20,000 primary cells were seeded per well in 96-well plates (n = 3 biological replicates). Following overnight incubation, cells were treated with a titration of LD5095 starting at 1μM with a 5-fold serial dilution. Cell viability was measured using the CellTiter-Glo assay, with results calculated as a percentage relative to the DMSO-treated control group.

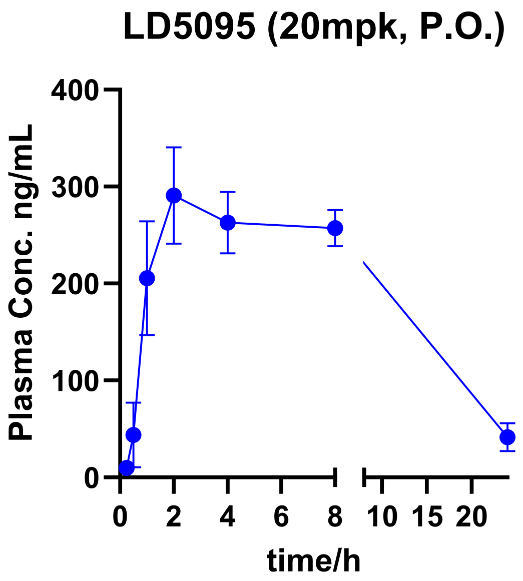


**Figure S6.** Plasma concentration-time profile of LD5095 following oral administration. Pharmacokinetic profile of LD5095 in mice (n = 3) following a single oral dose of 20 mg/kg. Plasma samples were collected at the indicated time points, and LD5095 concentrations were quantified to determine systemic exposure. Data points represent the mean plasma concentration ± standard deviation (SD).

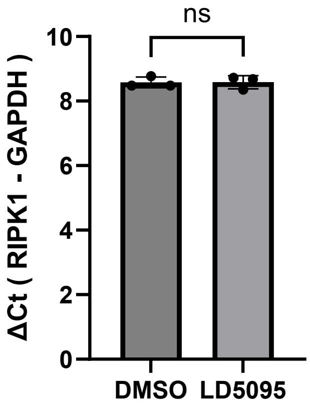


**Figure S7:** Quantitative PCR (qPCR) analysis of RIPK1 in Jurkat cells. Jurkat cells were treated for 24 hours with DMSO or LD5095 (100 nM) prior to collection.

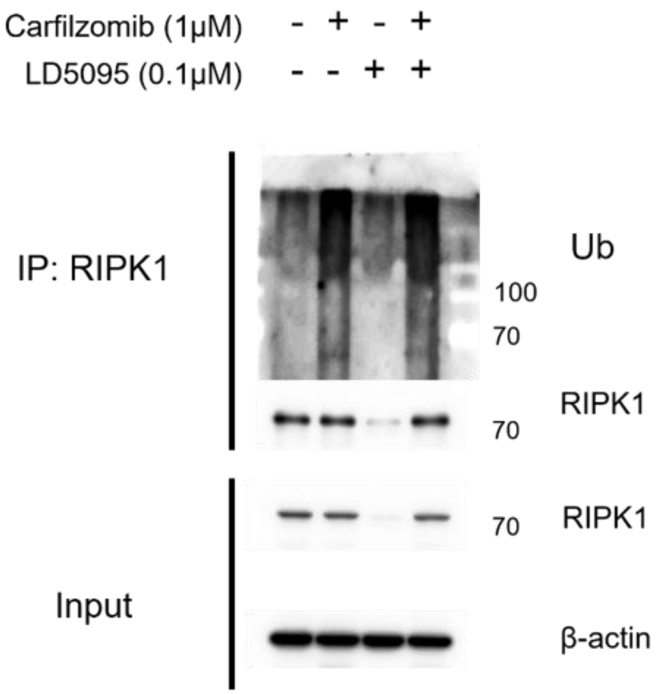


**Figure S8:** Effect of LD5095 on the ubiquitination of RIPK1 in Jurkat cells. Jurkat cells were pre-treated for 0.5 hours with Carfilzomib (1 μM), subsequently treated with DMSO or LD5095 (100 nM) for 4 hours prior to collection.

**Table S1. Pharmacokinetic Parameters of LD5095 in Mice via oral administration (20 mg/kg, n=3)**

| | Cl_plasma (mL/min/kg) | $T_{1/2}$ (h) | $C_{max}$ (ng/mL) | $AUC_{0\text{-}last}$ (h·ng/mL) | Vd (L/kg) |
|---|---|---|---|---|---|
| **LD5095** | 69.7±4.3 | 7.2±1.6 | 304±40 | 4795±298 | 0.043±0.010 |

**Table S2. RIPK1 SILAC Half-Life Summary**

| **System** | **Source** | **RIPK1 half-life** |
|---|---|---|
| Primary human B cells | Mathieson et al. 2018, dynamic SILAC | 46.3, 48.0 h; mean 47.1 h |
| Primary human NK cells | Mathieson et al. 2018, dynamic SILAC | 76.9, 71.1 h; mean 74.0 h |
| Primary human monocytes | Mathieson et al. 2018, dynamic SILAC | 49.0, 39.7 h; mean 44.4 h |
| Mouse embryonic neurons | Mathieson et al. 2018 | not quantified for RIPK1 |
| NIH3T3 mouse fibroblast cell line | Schwanhäusser et al. 2011 | 34.7, 49.0 h; mean 41.3 h |
| HeLa human cell line | Zecha et al. 2018, SILAC-TMT | protein $T_{1/2}$ = 65.0 h; label-turnover $T_{50}$ = 19.9 h |
| Footnote: For the HeLa table specifically: RIPK1 was quantified as Q13546;Q13546-2, with 13 unique peptides, 21.2% sequence coverage, 4 cell-culture replicates, k = 0.01066 /h, and $T_{1/2}$ = 65.0 h. | | |

HPLC analyses were performed on an Agilent 1260 Infinity LC/MS System (column, Agilent Eclipse plus C18, 4.6 mm × 100 mm, 3.5 μm; with the solvent system of MeCN/$H_2O$/FA, 0.7 mL/min; UV wavelength, maximal absorbance at 254 nm; temperature, ambient; injection volume, 5 μL).

Table S3. Purity of target compounds

| Sample | Purity % | Sample | Purity % |
|---|---|---|---|
| **3a** | 98.8 | **15a** | >99.0 |
| **3b** | >99.0 | **15b** | >99.0 |
| **3c** | 95.2 | **15c** | >99.0 |
| **3d** | 97.8 | **15d** | >99.0 |
| **5a** | 95.4 | **15e** | 95.4 |
| **5b** | 97.6 | **18a** | >99.0 |
| **5c** | 96.5 | **18b** | 98.6 |
| **5d** | >99.0 | **18c** | 98.5 |
| **5e** | 95.0 | **18d** | 99.0 |
| **5f** | >99.0 | **LD5095-NC** | 98.8 |

### $^{1}H$ NMR, $^{13}C$ NMR spectra of representative compounds

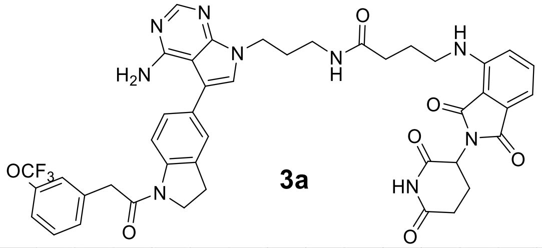


**3a**

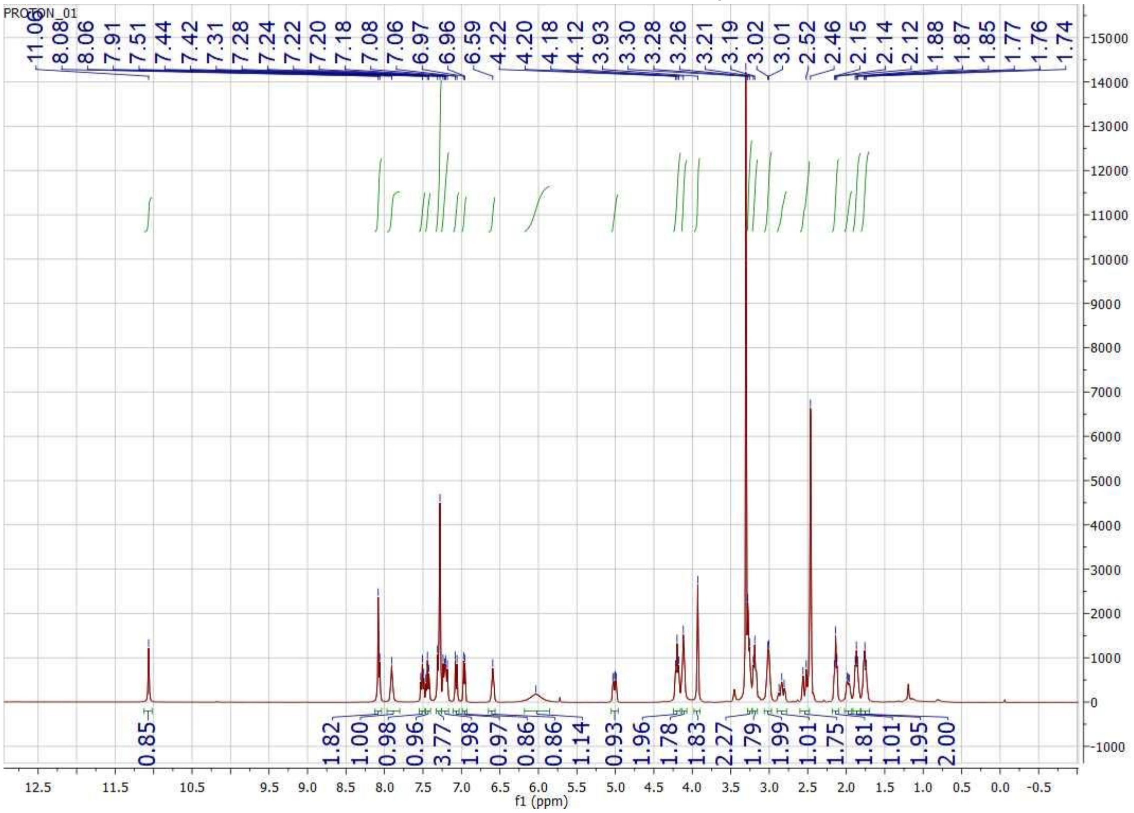

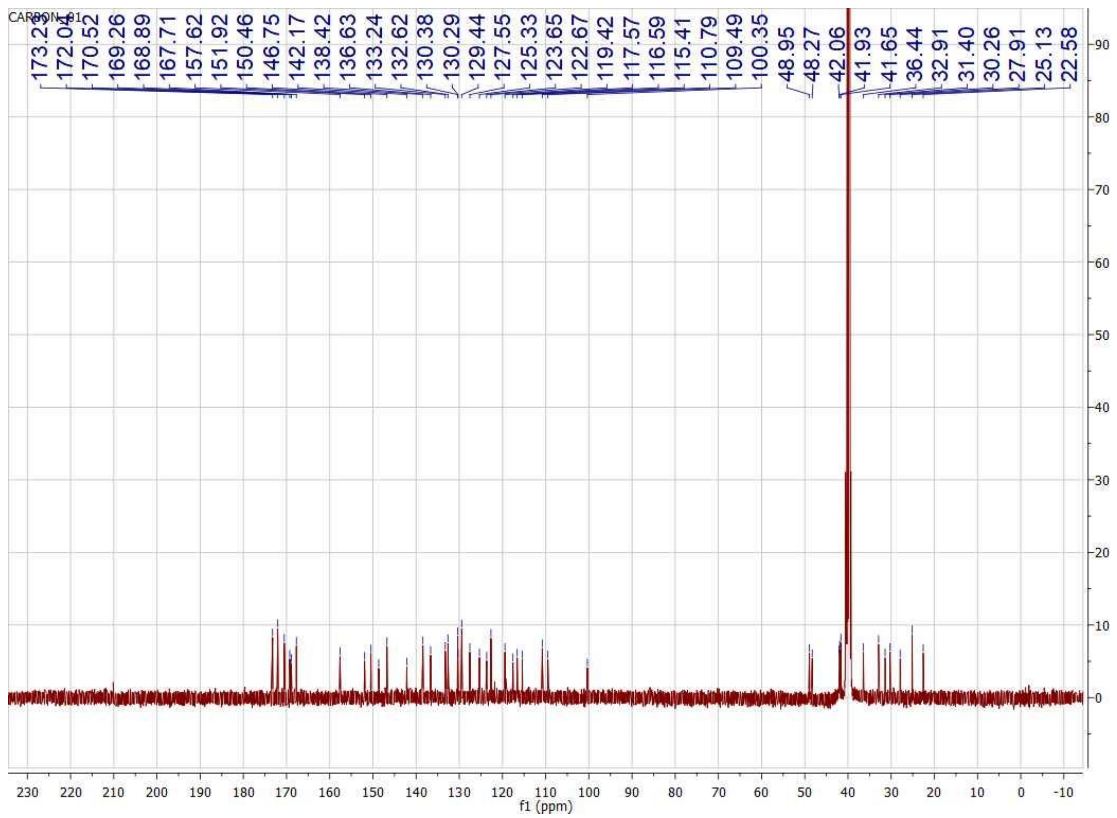

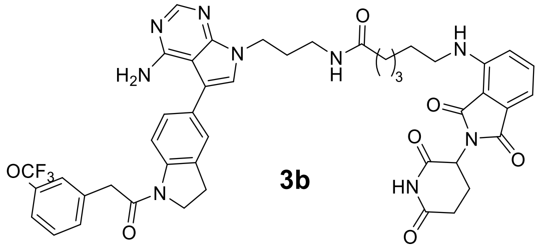

**3b**

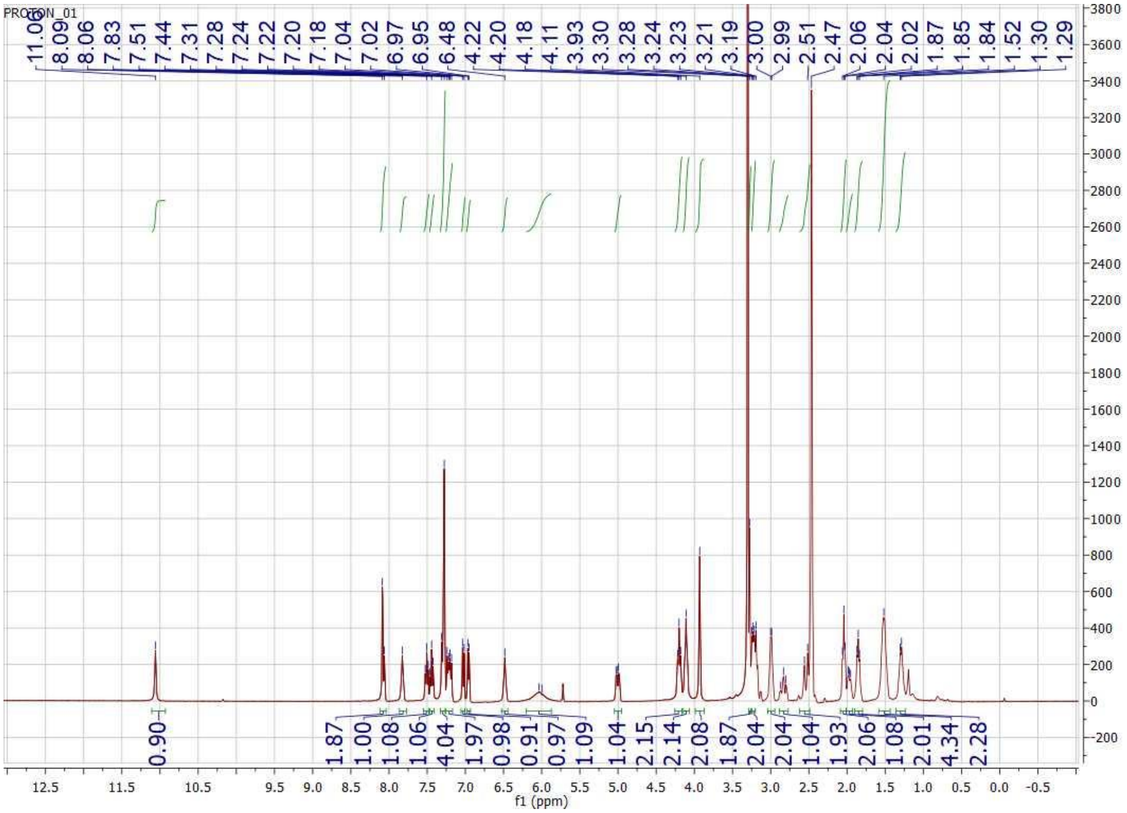
PROTON_01
11.06 8.09 8.06 7.83 7.51 7.44 7.31 7.28 7.24 7.22 7.20 7.18 7.04 7.02 6.97 6.95 6.48 4.22 4.20 4.18 4.11 3.93 3.30 3.28 3.24 3.23 3.21 3.19 3.00 2.99 2.51 2.47 2.06 2.04 2.02 1.87 1.85 1.84 1.52 1.30 1.29
0.90 1.87 1.00 1.08 1.06 4.04 1.97 0.98 0.91 0.97 1.09 1.04 2.15 2.14 2.08 1.87 2.04 2.04 1.04 1.93 2.06 1.08 2.01 4.34 2.28
12.5 11.5 10.5 9.5 9.0 8.5 8.0 7.5 7.0 6.5 6.0 5.5 5.0 4.5 4.0 3.5 3.0 2.5 2.0 1.5 1.0 0.5 0.0 -0.5
f1 (ppm)
3800 3600 3400 3200 3000 2800 2600 2400 2200 2000 1800 1600 1400 1200 1000 800 600 400 200 0 -200

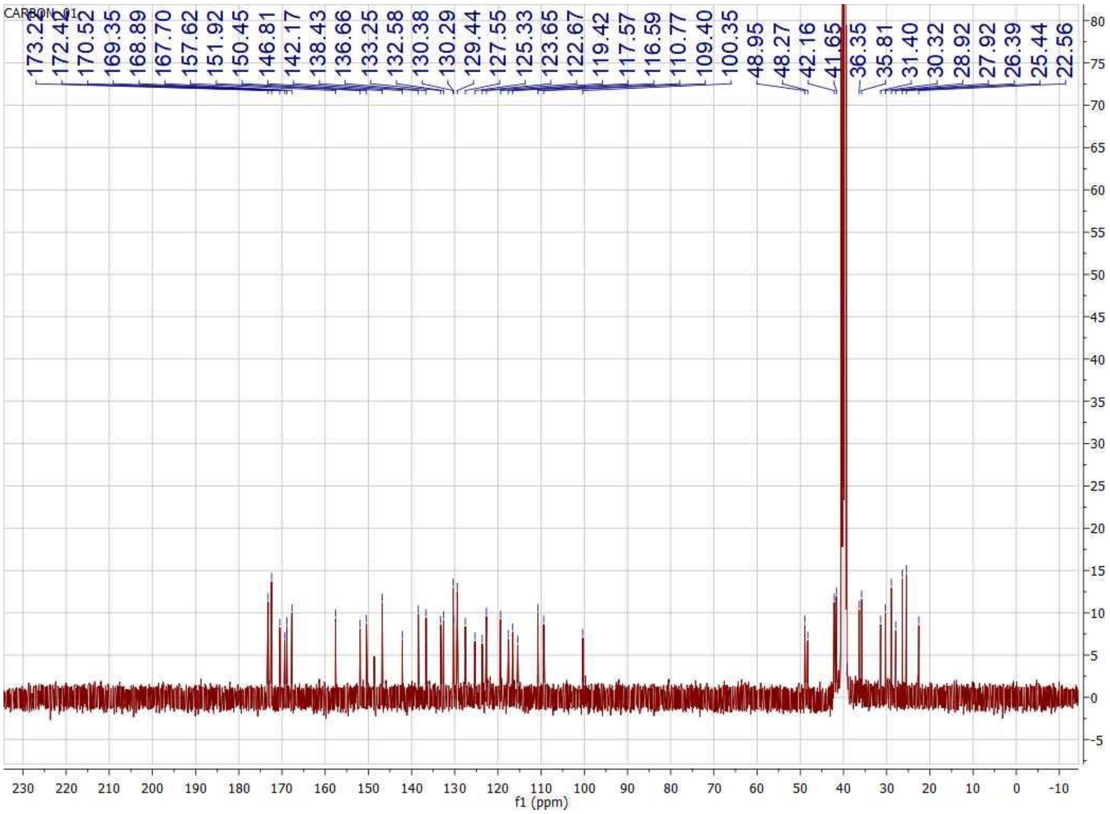

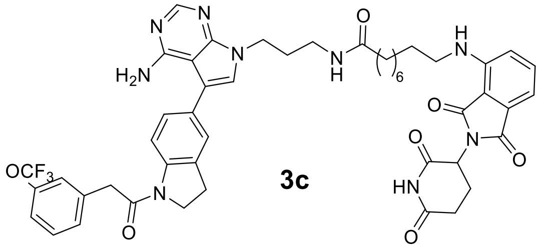
N
N
N
H2N
N
H
O
6
H
N
O
N
O
HN
O
O
OCF3
N
O
3c

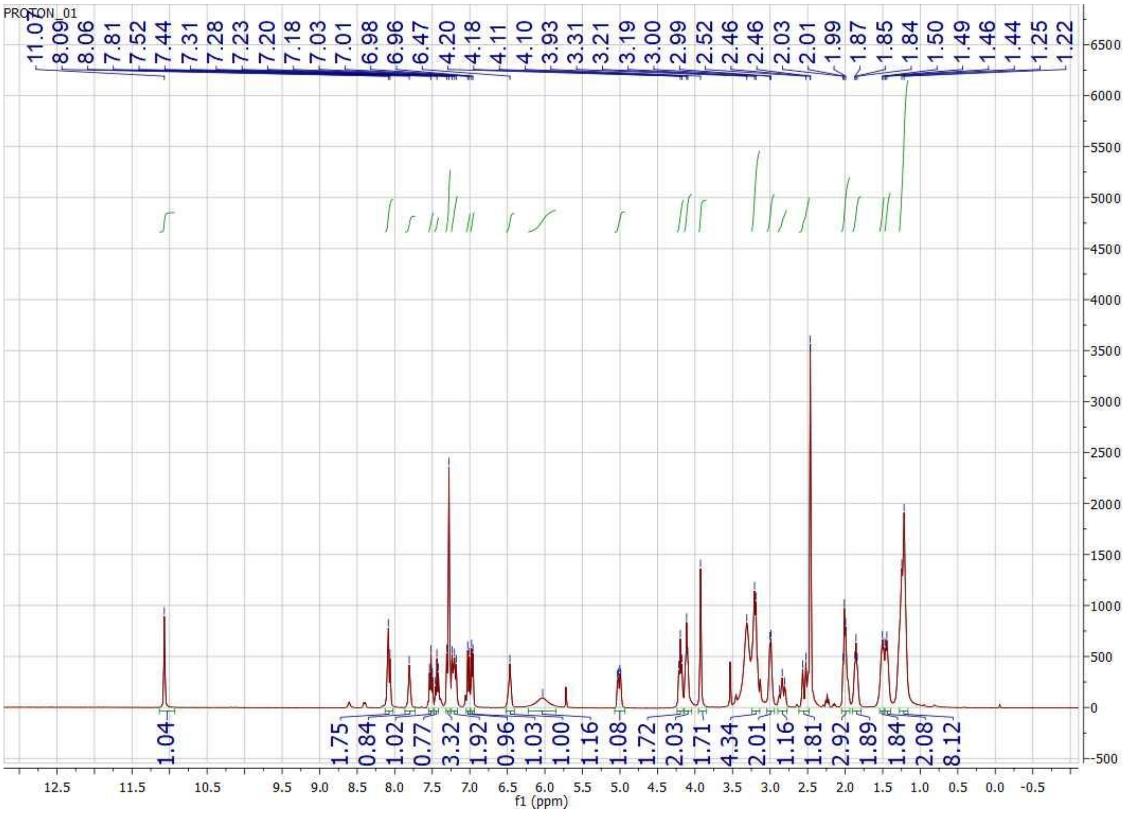
PROTON_01
11.07 8.09 8.06 7.81 7.52 7.44 7.31 7.28 7.23 7.20 7.18 7.03 7.01 6.98 6.96 6.47 4.20 4.18 4.11 4.10 3.93 3.31 3.21 3.19 3.00 2.99 2.52 2.46 2.46 2.03 2.01 1.99 1.87 1.85 1.84 1.50 1.49 1.46 1.44 1.25 1.22
1.04 1.75 0.84 1.02 0.77 3.32 1.92 0.96 1.03 1.00 1.16 1.08 1.72 2.03 1.71 4.34 2.01 1.16 1.81 2.92 1.89 1.84 2.08 8.12
6500 6000 5500 5000 4500 4000 3500 3000 2500 2000 1500 1000 500 0 -500
12.5 11.5 10.5 9.5 9.0 8.5 8.0 7.5 7.0 6.5 6.0 5.5 5.0 4.5 4.0 3.5 3.0 2.5 2.0 1.5 1.0 0.5 0.0 -0.5
f1 (ppm)

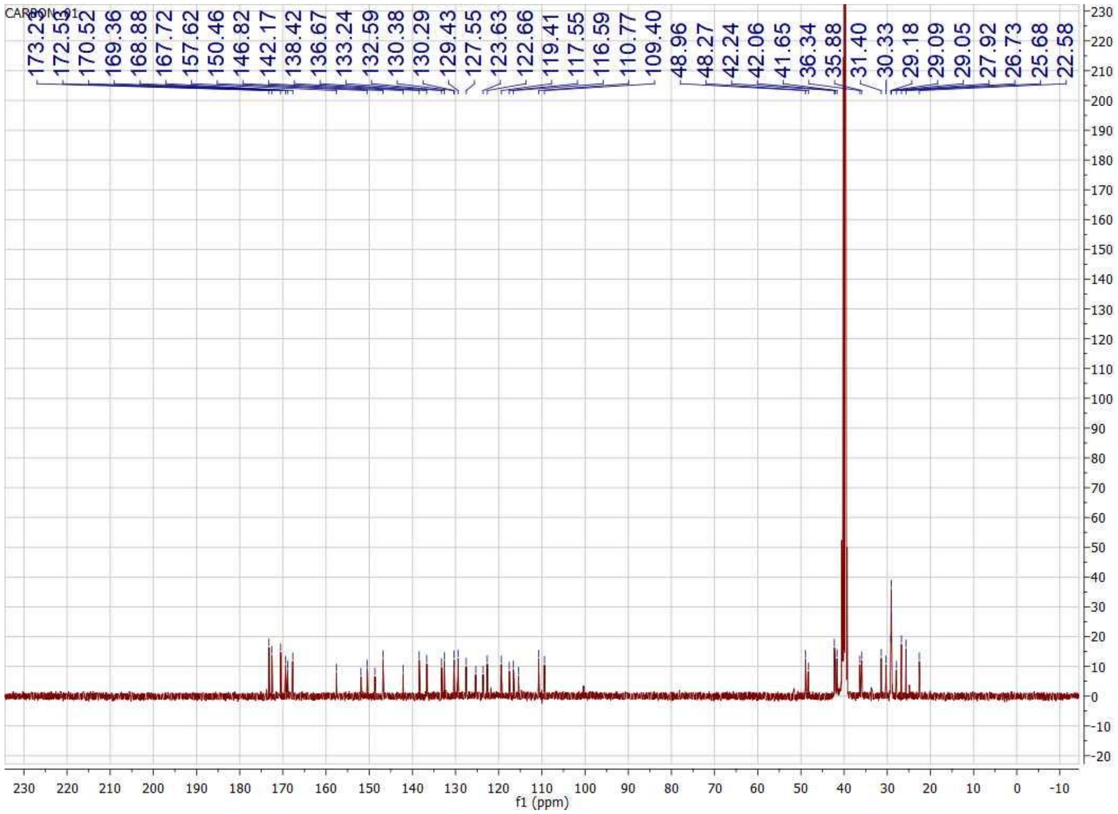
CARBON_01
173.29
172.55
170.52
169.36
168.88
167.72
157.62
150.46
146.82
142.17
138.42
136.67
133.24
132.59
130.38
130.29
129.43
127.55
123.63
122.66
119.41
117.55
116.59
110.77
109.40
48.96
48.27
42.24
42.06
41.65
36.34
35.88
31.40
30.33
29.18
29.09
29.05
27.92
26.73
25.68
22.58
f1 (ppm)

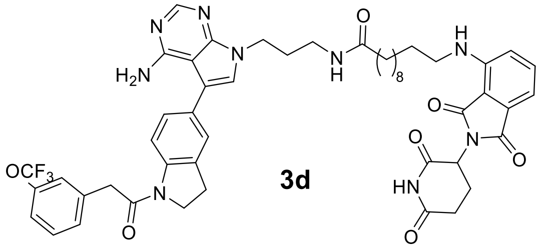

N
N
H$_2$N
N
N
H
O
8
H
N
OCF$_3$
O
N
O
N
O
O
HN
O
O


**3d**

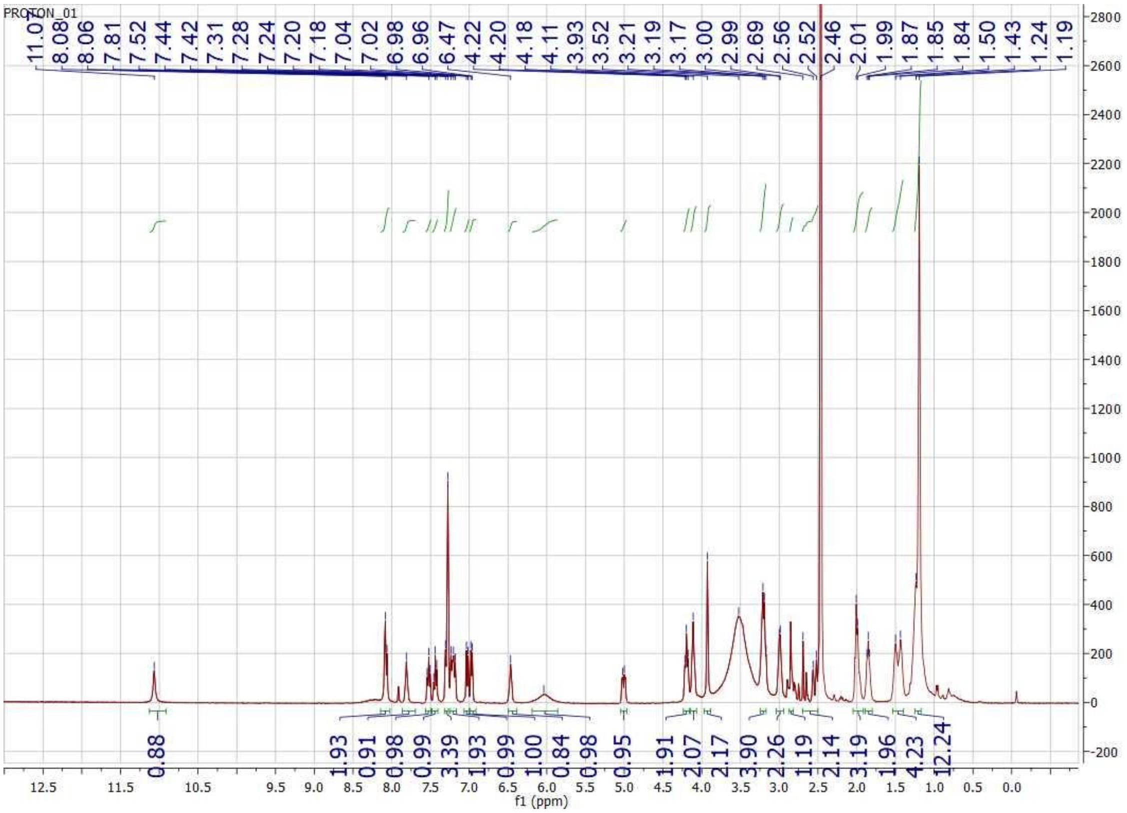

PROTON_01
11.07 8.08 8.06 7.81 7.52 7.44 7.42 7.31 7.28 7.24 7.20 7.18 7.04 7.02 6.98 6.96 6.47 4.22 4.20 4.18 4.11 3.93 3.52 3.21 3.19 3.17 3.00 2.99 2.69 2.56 2.52 2.46 2.01 1.99 1.87 1.85 1.84 1.50 1.43 1.24 1.19
0.88 1.93 0.91 0.98 0.99 3.39 1.93 0.99 1.00 0.84 0.98 0.95 1.91 2.07 2.17 3.90 2.26 1.19 2.14 3.19 1.96 4.23 12.24
f1 (ppm)

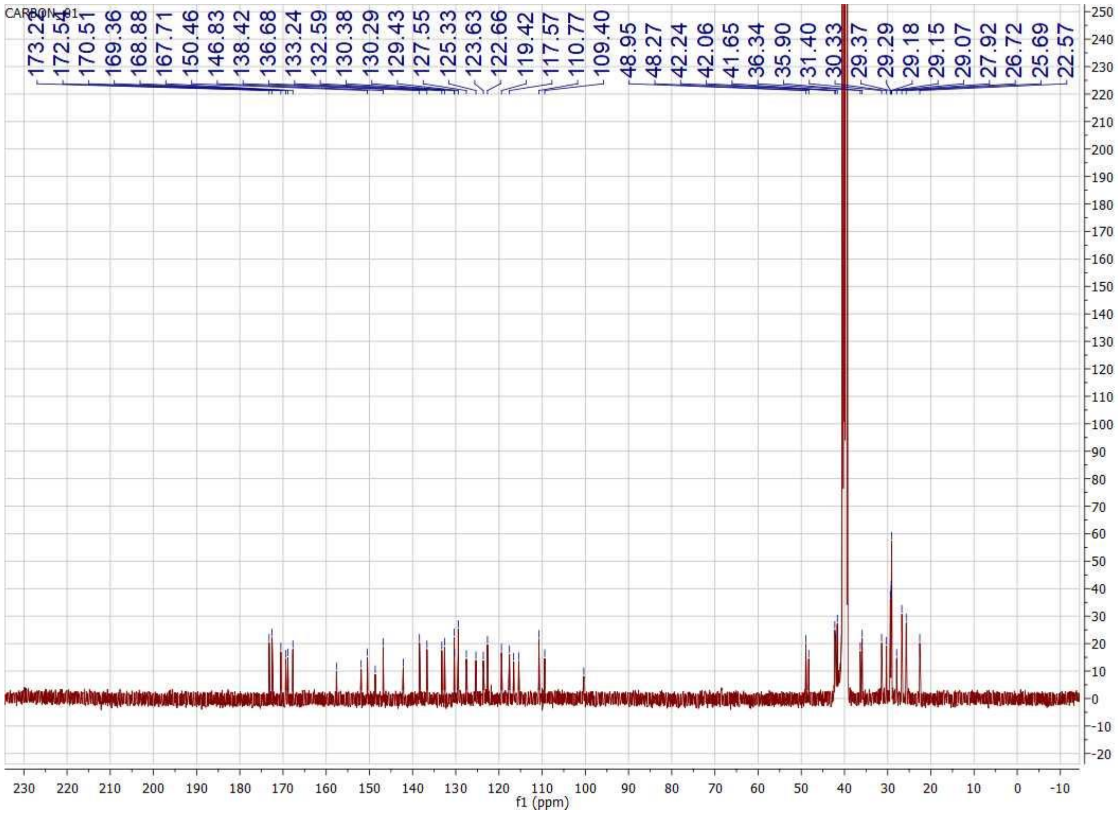
CARBON_01
173.22
172.54
170.51
169.36
168.88
167.71
150.46
146.83
138.42
136.68
133.24
132.59
130.38
130.29
129.43
127.55
125.33
123.63
122.66
119.42
117.57
110.77
109.40
48.95
48.27
42.24
42.06
41.65
36.34
35.90
31.40
30.33
29.37
29.29
29.18
29.15
29.07
27.92
26.72
25.69
22.57
f1 (ppm)

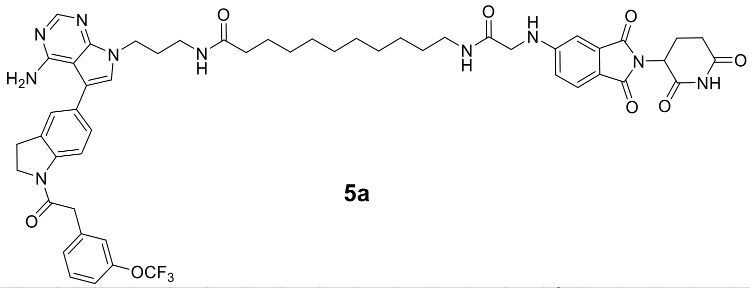

**5a**

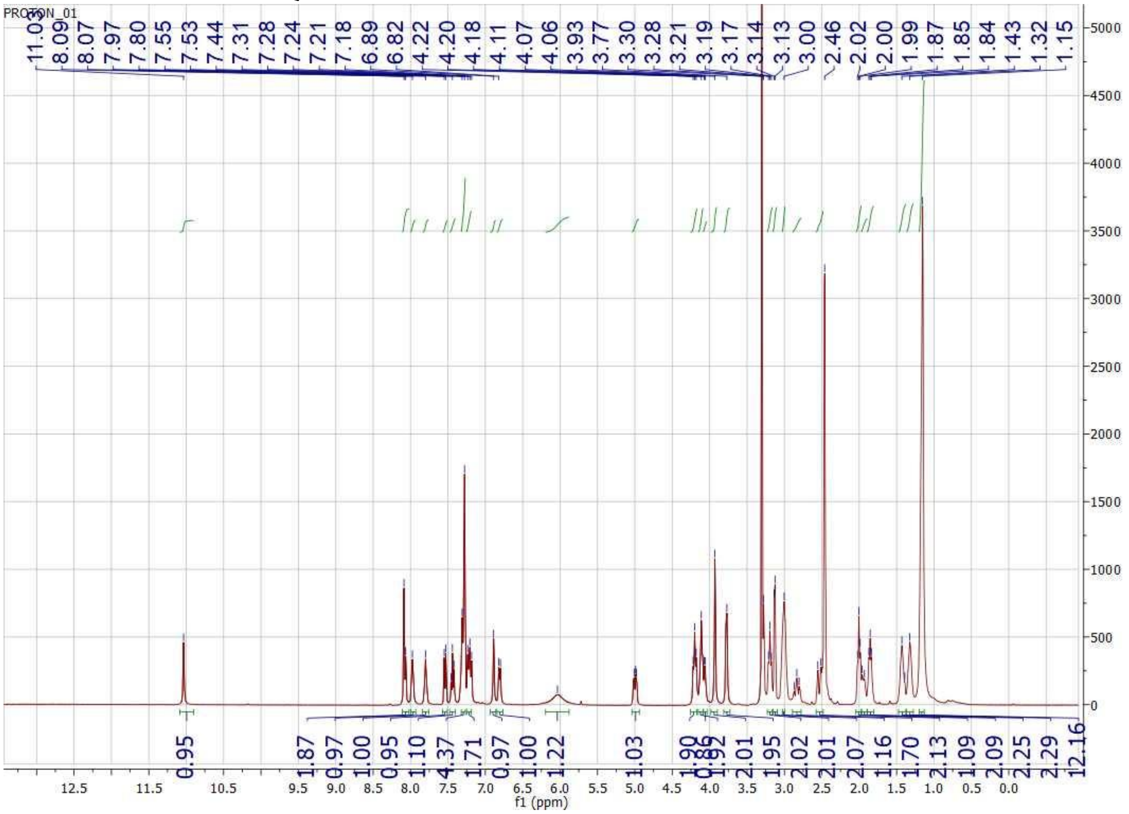

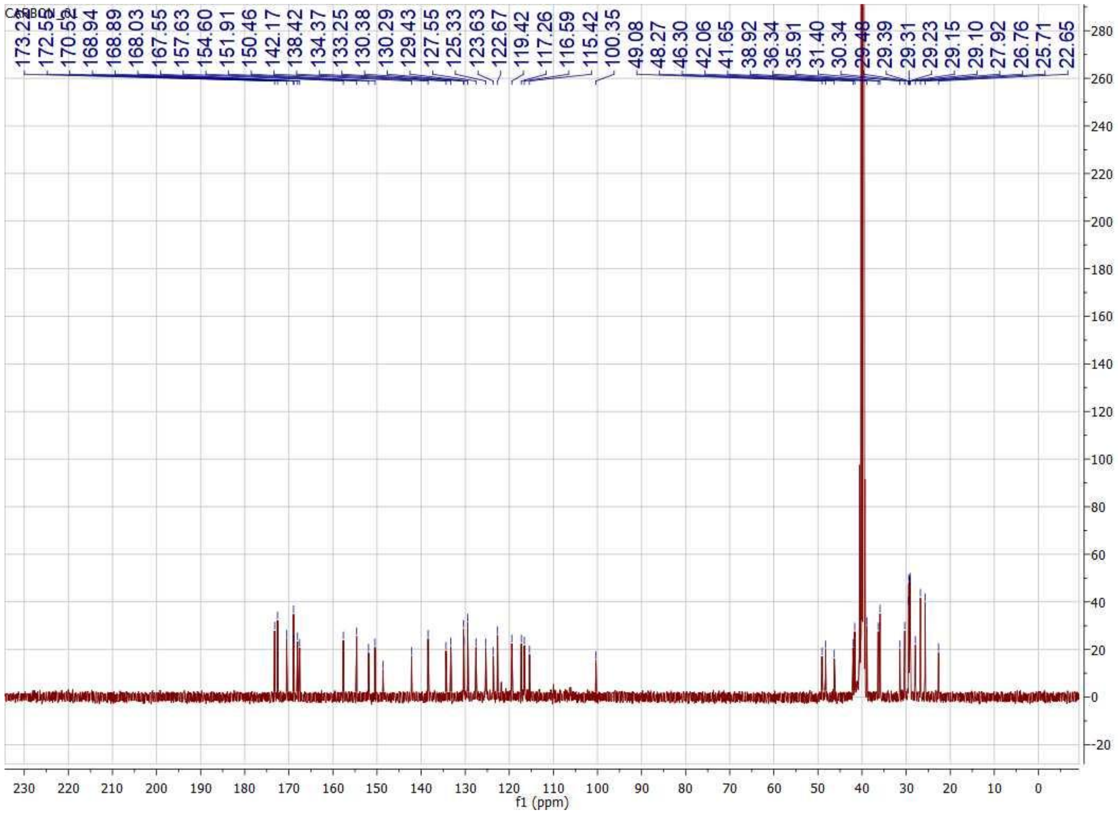
CARBON_01
173.22
172.55
170.53
168.94
168.89
168.03
167.55
157.63
154.60
151.91
150.46
142.17
138.42
134.37
133.25
130.38
130.29
129.43
127.55
125.33
123.63
122.67
119.42
117.26
116.59
115.42
100.35
49.08
48.27
46.30
42.06
41.65
38.92
36.34
35.91
31.40
30.34
29.39
29.31
29.23
29.15
29.10
27.92
26.76
25.71
22.65
f1 (ppm)

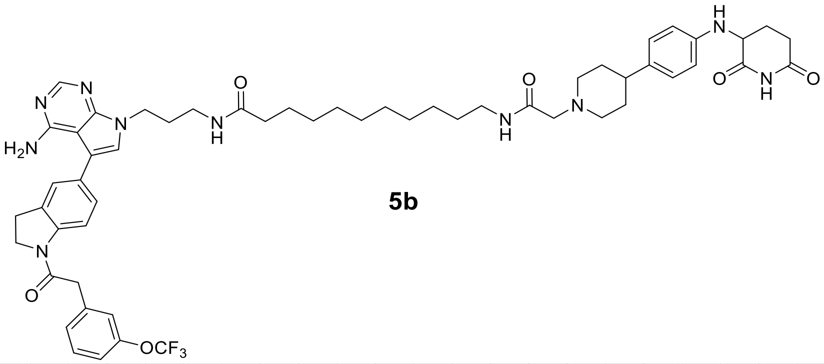

**5b**

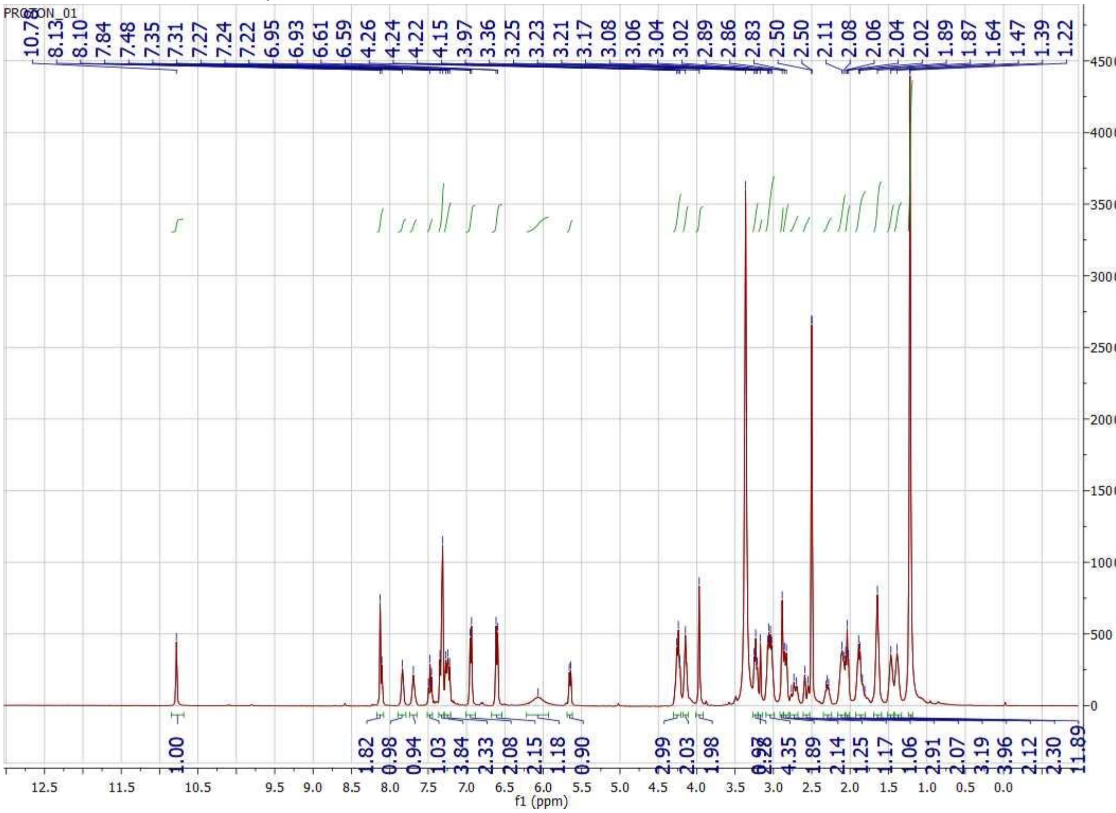

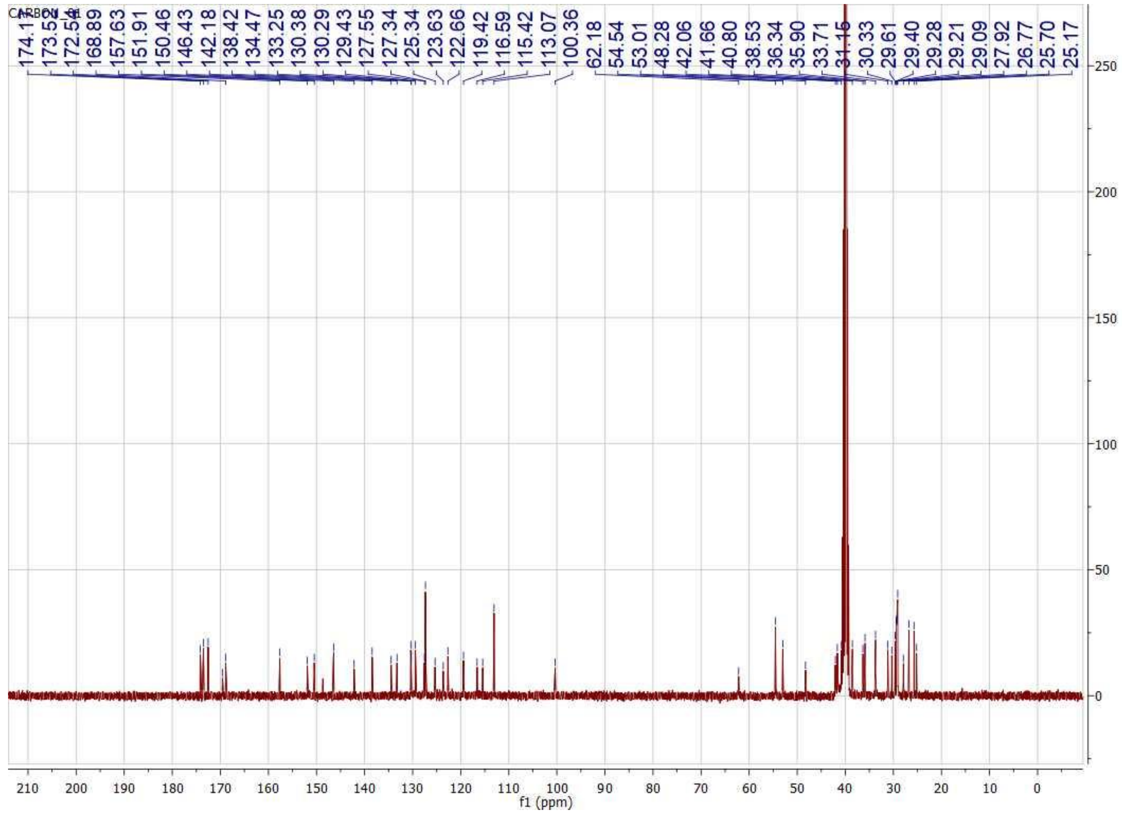
CARBON_01
174.11
173.55
172.58
168.89
157.63
151.91
150.46
146.43
142.18
138.42
134.47
133.25
130.38
130.29
129.43
127.55
127.34
125.34
123.63
122.66
119.42
116.59
115.42
113.07
100.36
62.18
54.54
53.01
48.28
42.06
41.66
40.80
38.53
36.34
35.90
33.71
31.16
30.33
29.61
29.40
29.28
29.21
29.09
27.92
26.77
25.70
25.17
f1 (ppm)

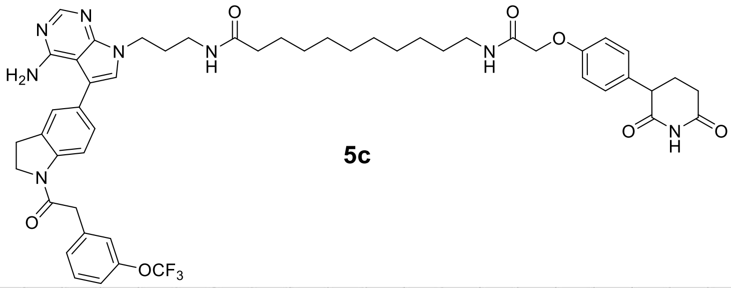

**5c**

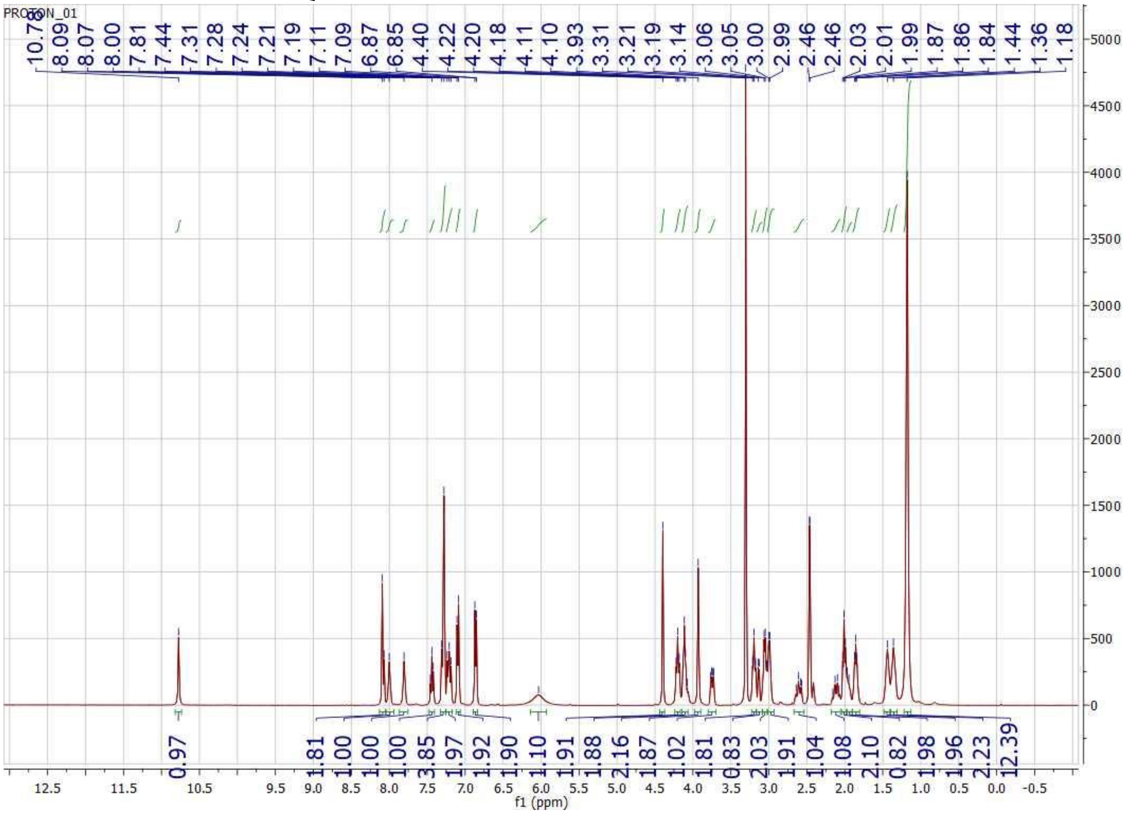

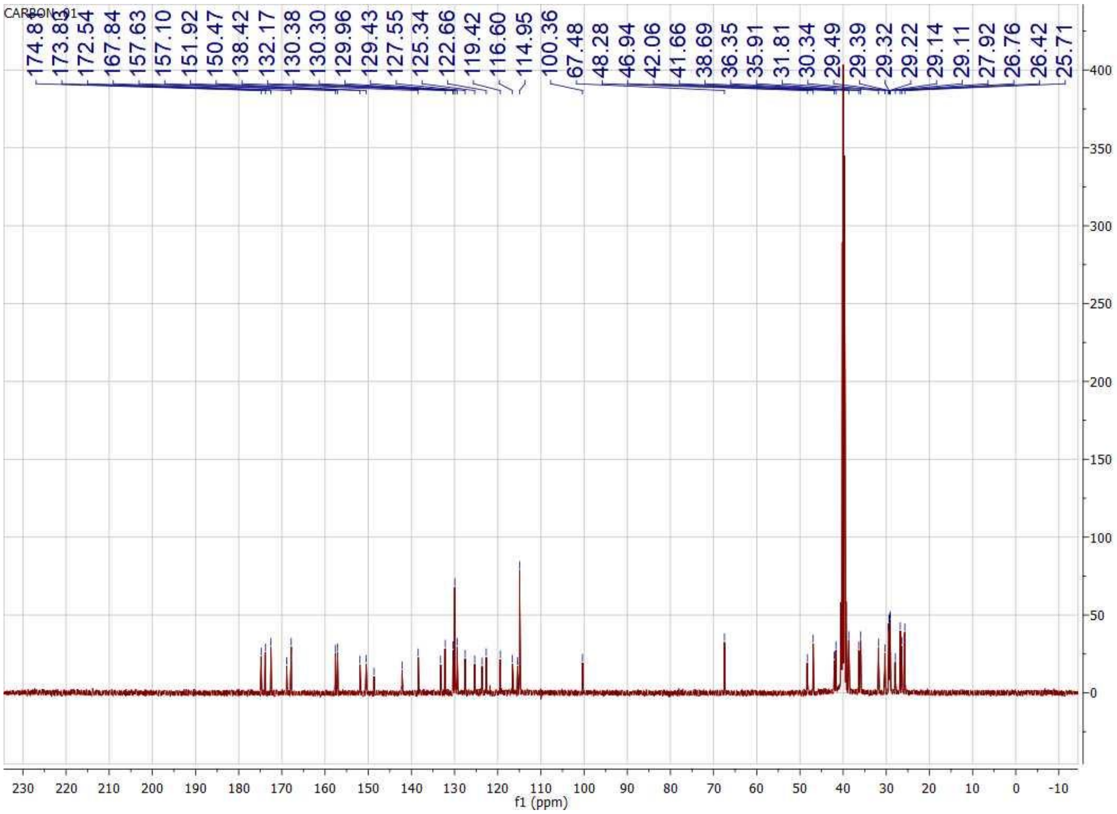

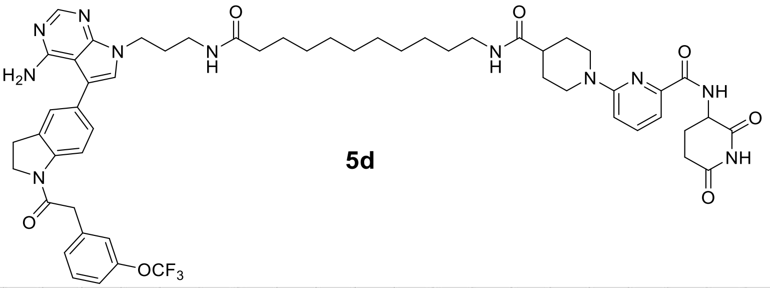

**5d**

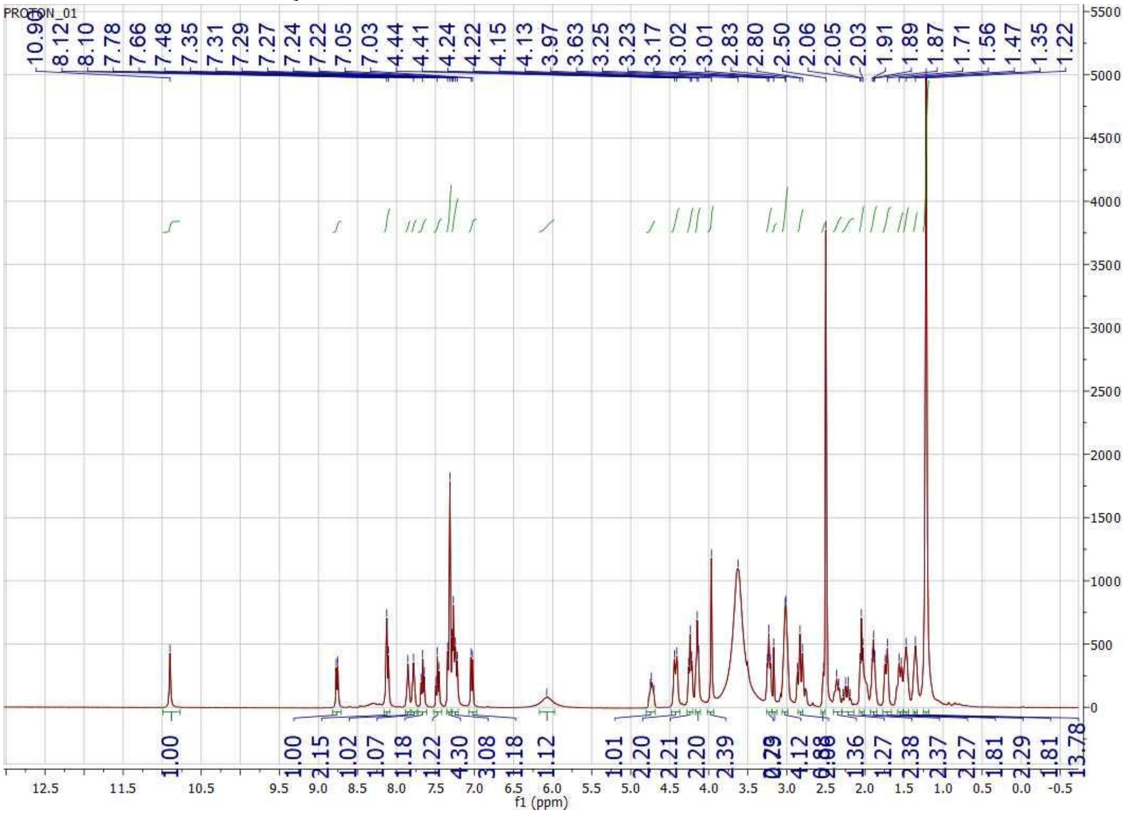
PROTON_01
10.90
8.12
8.10
7.78
7.66
7.48
7.35
7.31
7.29
7.27
7.24
7.22
7.05
7.03
4.44
4.41
4.24
4.22
4.15
4.13
3.97
3.63
3.25
3.23
3.17
3.02
3.01
2.83
2.80
2.50
2.06
2.05
2.03
1.91
1.89
1.87
1.71
1.56
1.47
1.35
1.22
1.00
1.00
2.15
1.02
1.07
1.18
1.22
4.30
3.08
1.18
1.12
1.01
2.20
2.21
2.20
2.39
1.36
1.27
2.38
2.37
2.27
1.81
2.29
1.81
13.78
f1 (ppm)

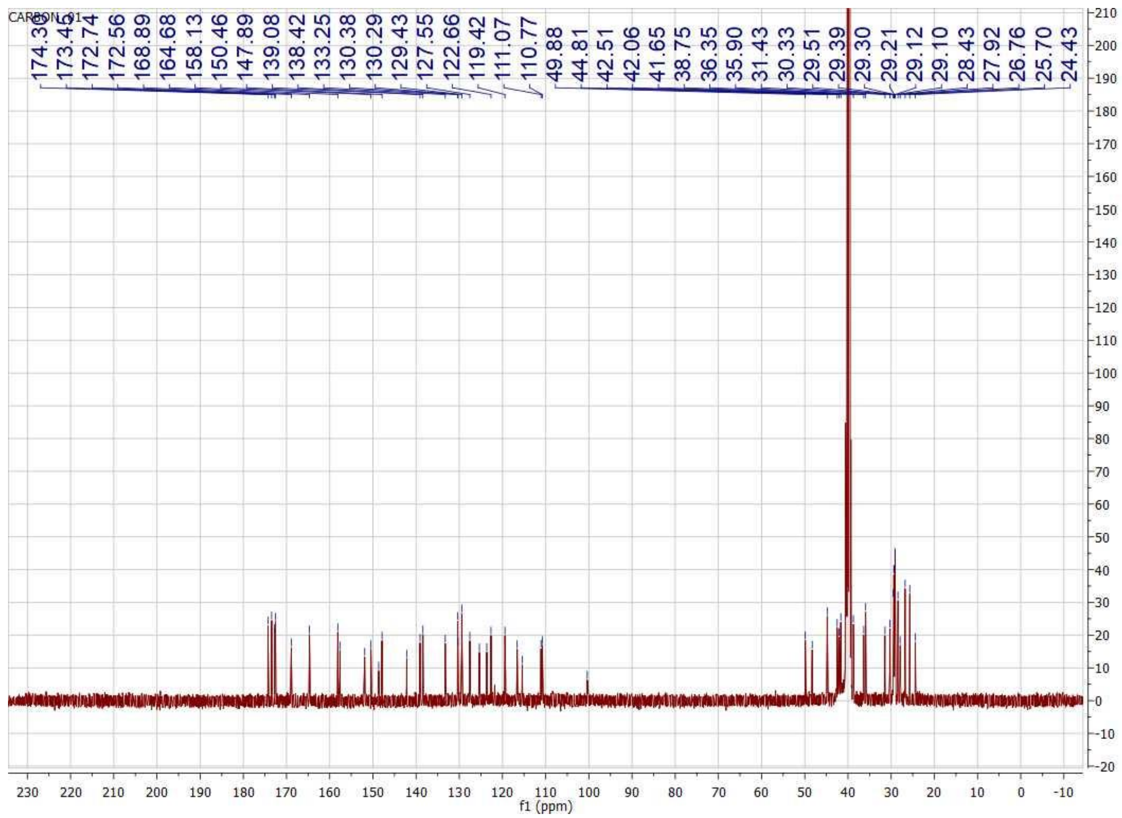
CARBON_01
174.36
173.45
172.74
172.56
168.89
164.68
158.13
150.46
147.89
139.08
138.42
133.25
130.38
130.29
129.43
127.55
122.66
119.42
111.07
110.77
49.88
44.81
42.51
42.06
41.65
38.75
36.35
35.90
31.43
30.33
29.51
29.39
29.30
29.21
29.12
29.10
28.43
27.92
26.76
25.70
24.43
f1 (ppm)

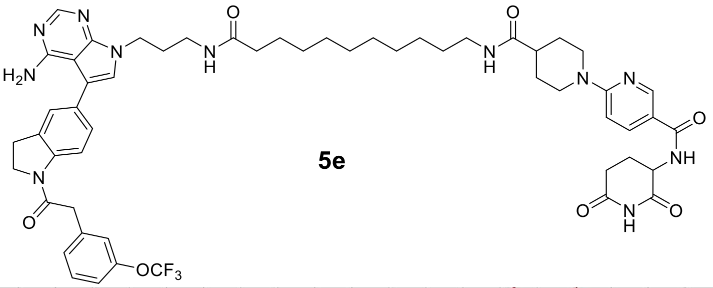
N
N
H2N
N
H
O
H
O
N
N
N
O
NH
O
N
H
O
5e
N
O
OCF3

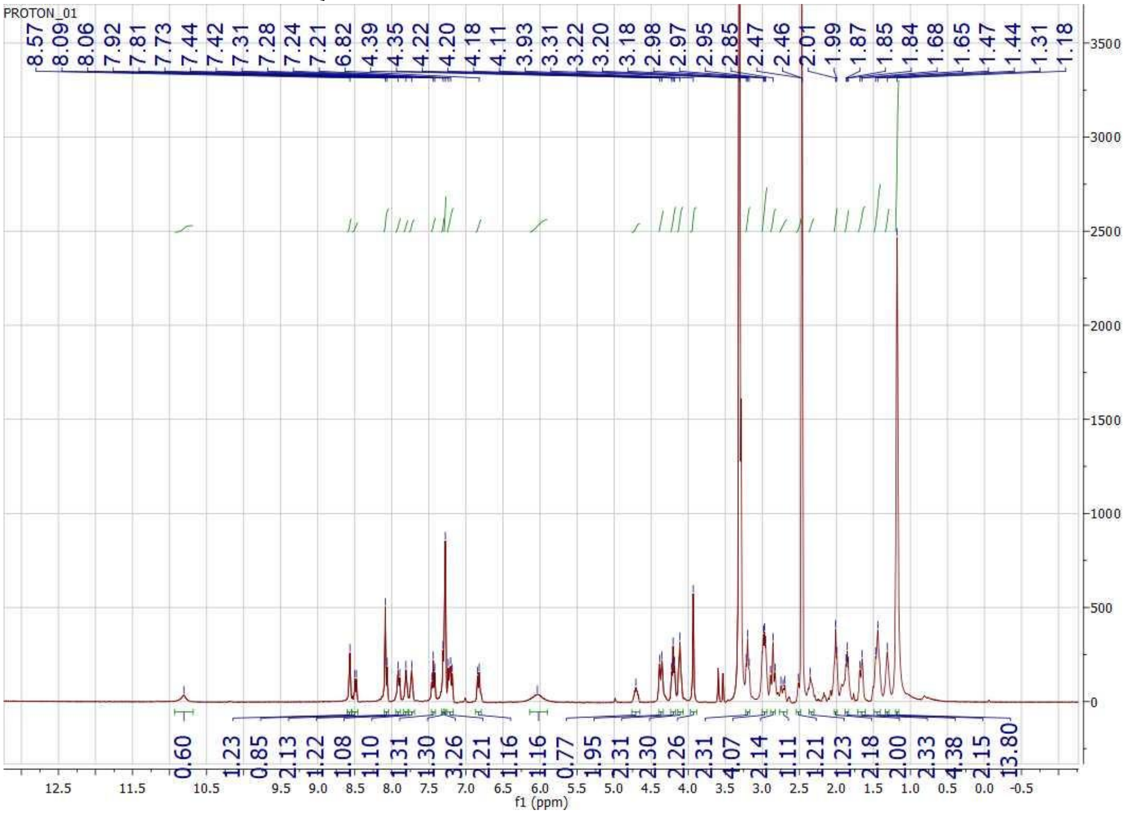
PROTON_01
8.57
8.09
8.06
7.92
7.81
7.73
7.44
7.42
7.31
7.28
7.24
7.21
6.82
4.39
4.35
4.22
4.20
4.18
4.11
3.93
3.31
3.22
3.20
3.18
2.98
2.97
2.95
2.85
2.47
2.46
2.01
1.99
1.87
1.85
1.84
1.68
1.65
1.47
1.44
1.31
1.18
0.60
1.23
0.85
2.13
1.22
1.08
1.10
1.31
1.30
3.26
2.21
1.16
1.16
0.77
1.95
2.31
2.30
2.26
2.31
4.07
2.14
1.11
1.21
1.23
2.18
2.00
2.33
4.38
2.15
13.80
f1 (ppm)

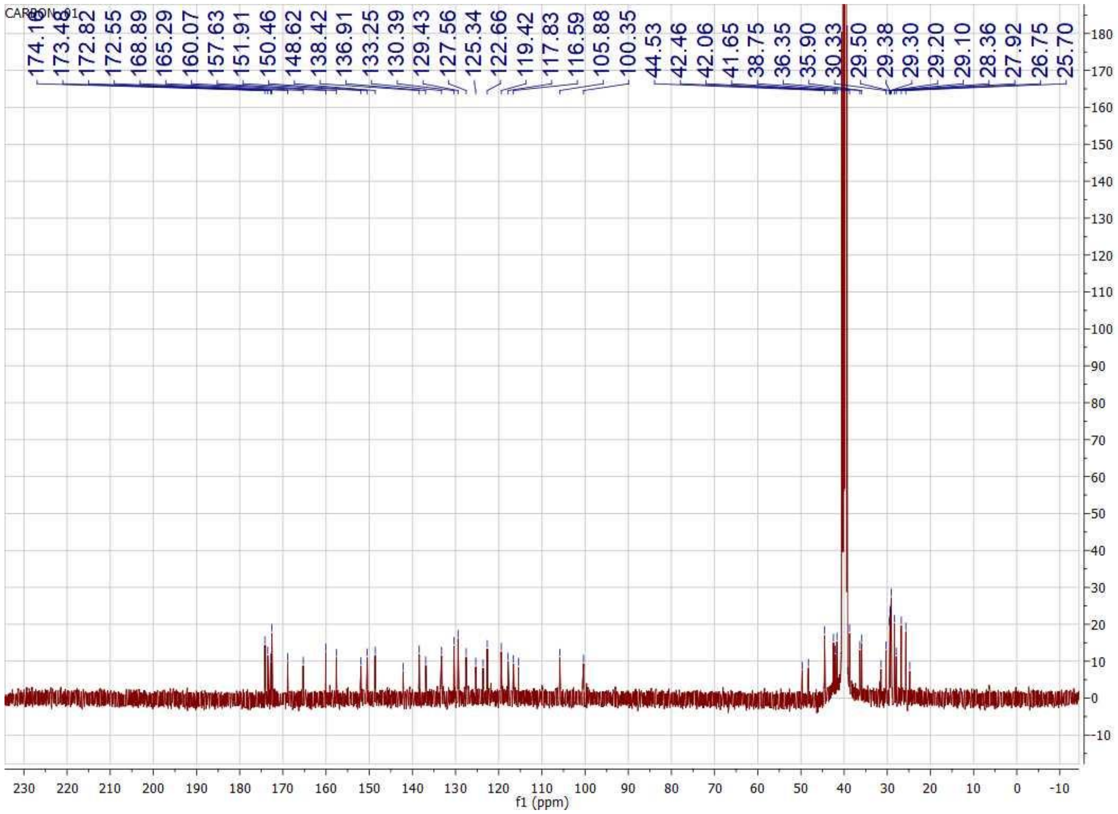
CARBON_01
174.16
173.48
172.82
172.55
168.89
165.29
160.07
157.63
151.91
150.46
148.62
138.42
136.91
133.25
130.39
129.43
127.56
125.34
122.66
119.42
117.83
116.59
105.88
100.35
44.53
42.46
42.06
41.65
38.75
36.35
35.90
30.33
29.50
29.38
29.30
29.20
29.10
28.36
27.92
26.75
25.70
f1 (ppm)

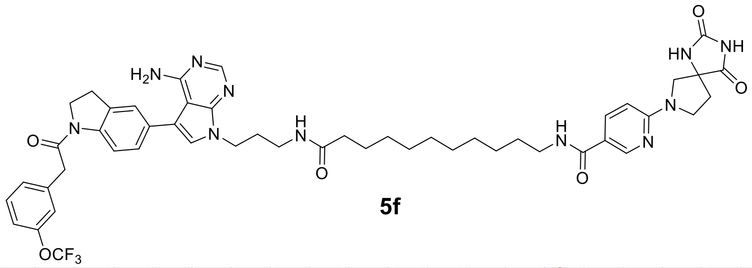

**5f**

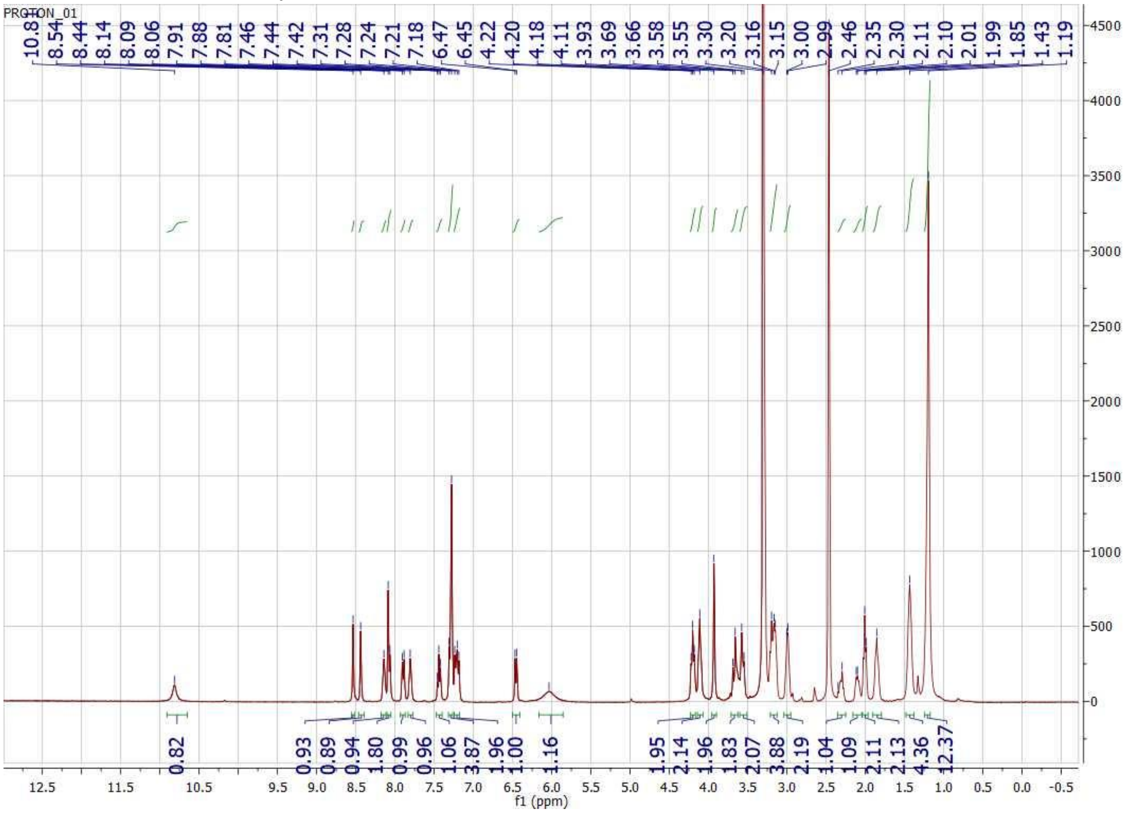
PROTON_01
10.81 8.54 8.44 8.14 8.09 8.06 7.91 7.88 7.81 7.46 7.44 7.42 7.31 7.28 7.24 7.21 7.18 6.47 6.45 4.22 4.20 4.18 4.11 3.93 3.69 3.66 3.58 3.55 3.30 3.20 3.16 3.15 3.00 2.99 2.46 2.35 2.30 2.11 2.10 2.01 1.99 1.85 1.43 1.19
0.82 0.93 0.89 0.94 1.80 0.99 0.96 1.06 3.87 1.96 1.00 1.16 1.95 2.14 1.96 1.83 2.07 3.88 2.19 1.04 1.09 2.11 2.13 4.36 12.37
f1 (ppm)

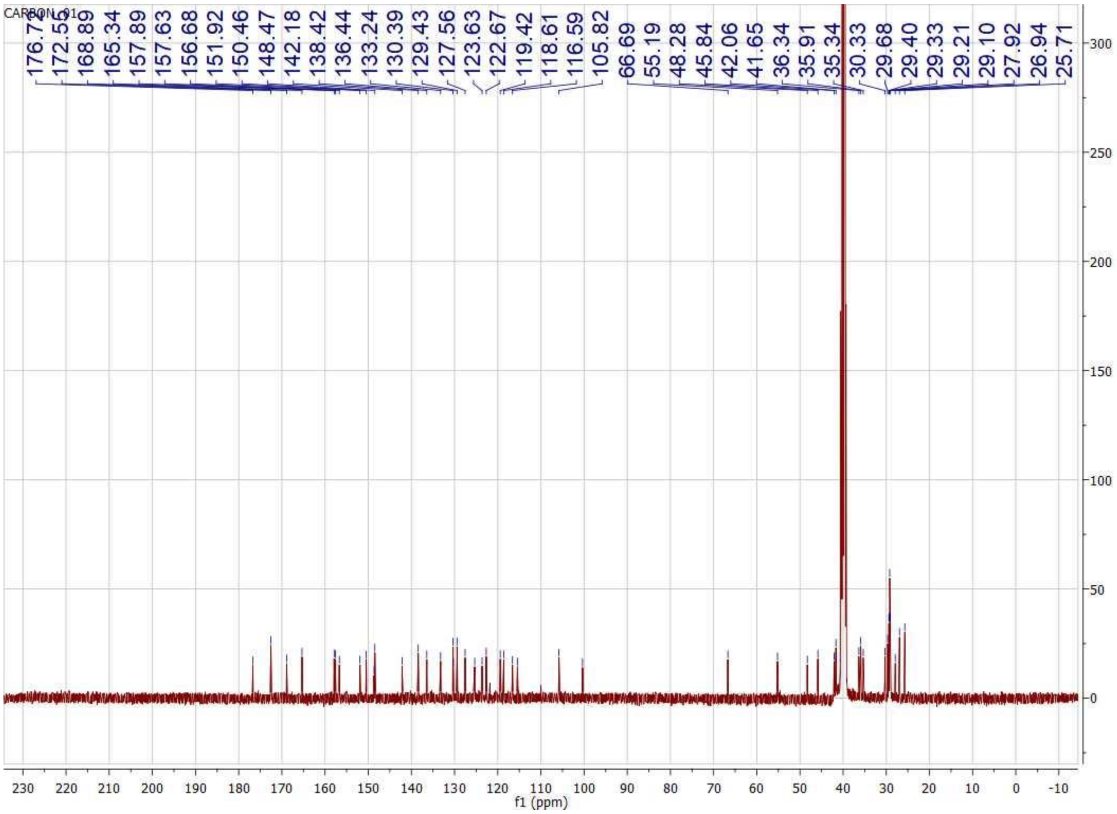
CARBON_01
176.72
172.55
168.89
165.34
157.89
157.63
156.68
151.92
150.46
148.47
142.18
138.42
136.44
133.24
130.39
129.43
127.56
123.63
122.67
119.42
118.61
116.59
105.82
66.69
55.19
48.28
45.84
42.06
41.65
36.34
35.91
35.34
30.33
29.68
29.40
29.33
29.21
29.10
27.92
26.94
25.71
f1 (ppm)

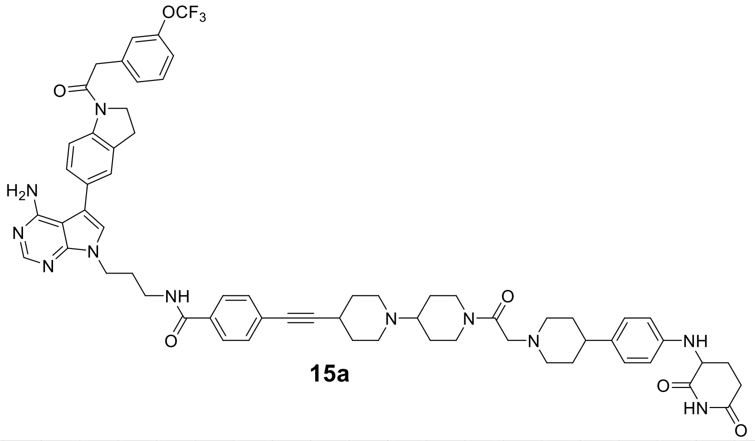

**15a**

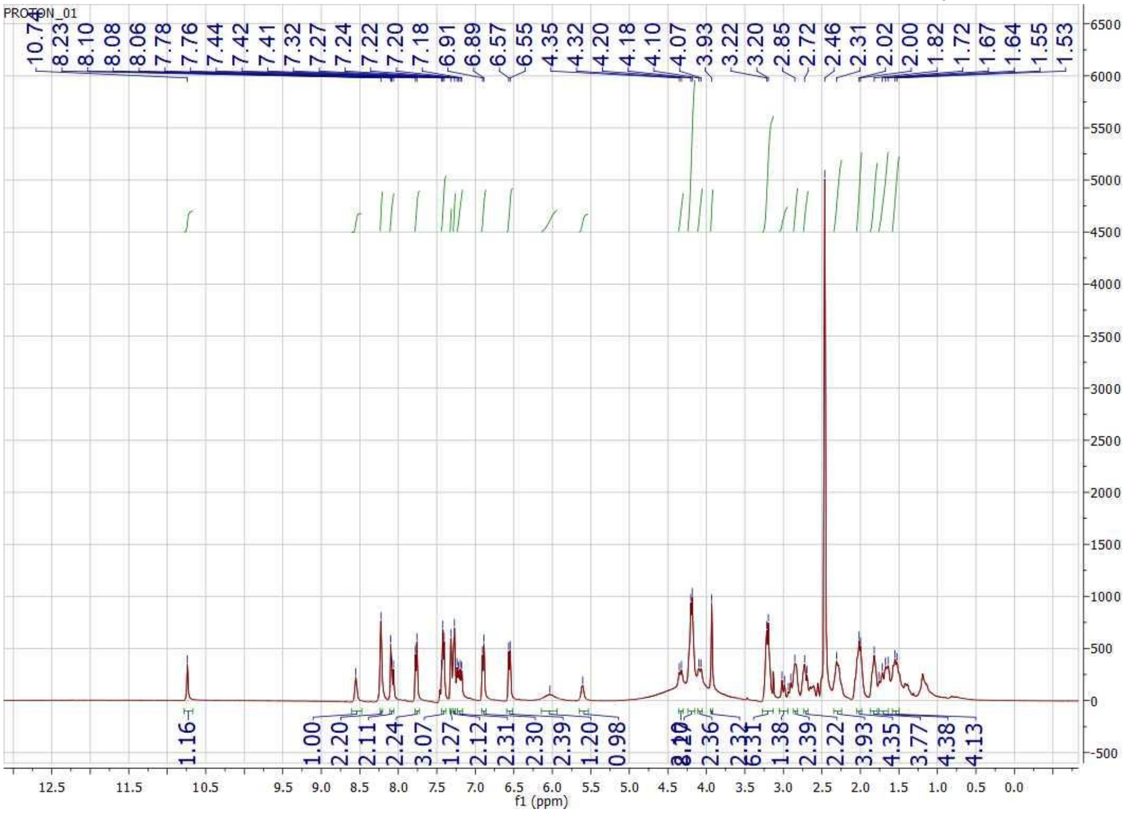

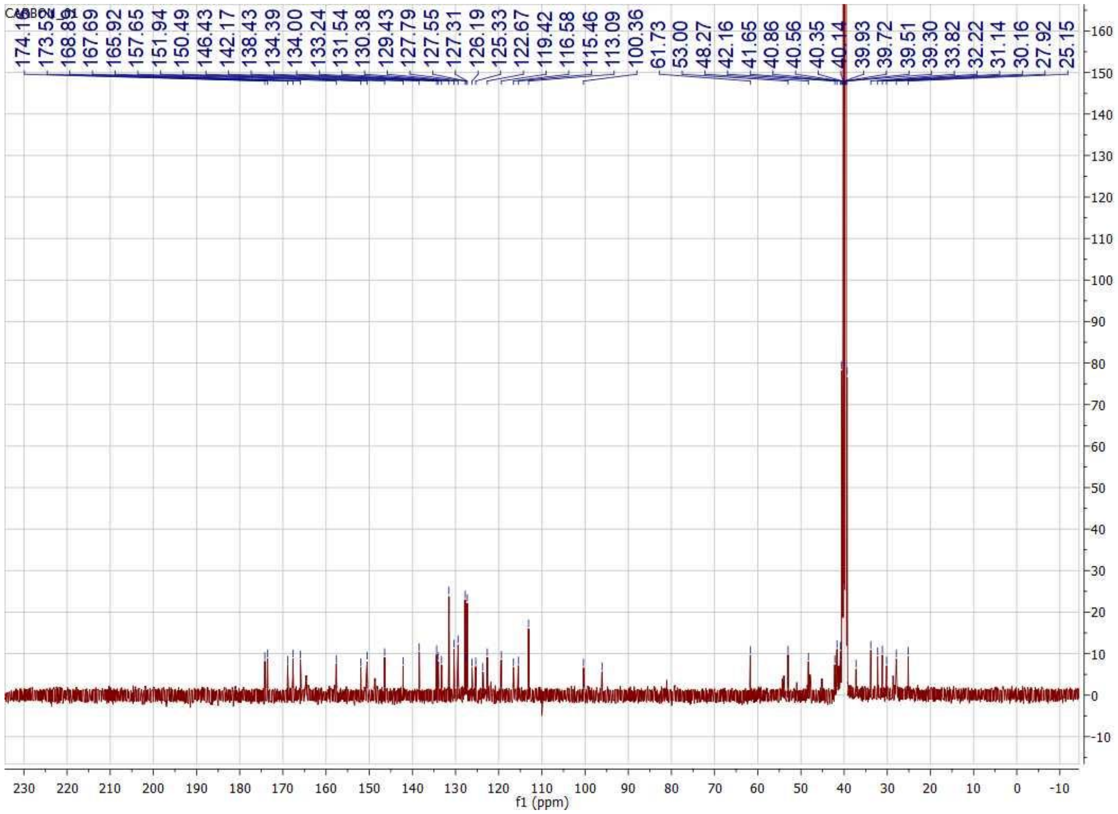
CARBON_01
174.11
173.55
168.81
167.69
165.92
157.65
151.94
150.49
146.43
142.17
138.43
134.39
134.00
133.24
131.54
130.38
129.43
127.79
127.55
127.31
126.19
125.33
122.67
119.42
116.58
115.46
113.09
100.36
61.73
53.00
48.27
42.16
41.65
40.86
40.56
40.35
40.14
39.93
39.72
39.51
39.30
33.82
32.22
31.14
30.16
27.92
25.15
f1 (ppm)

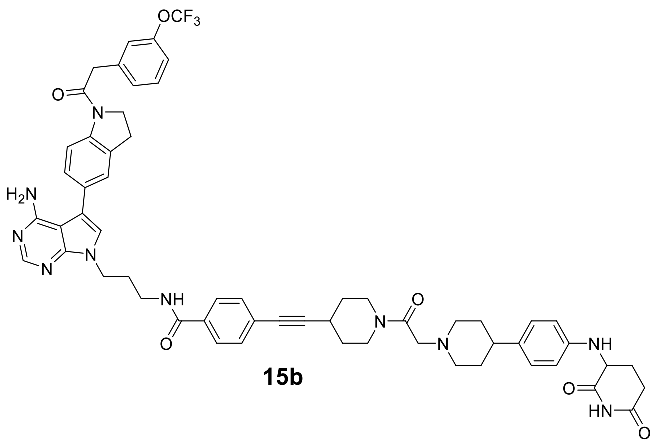

**15b**

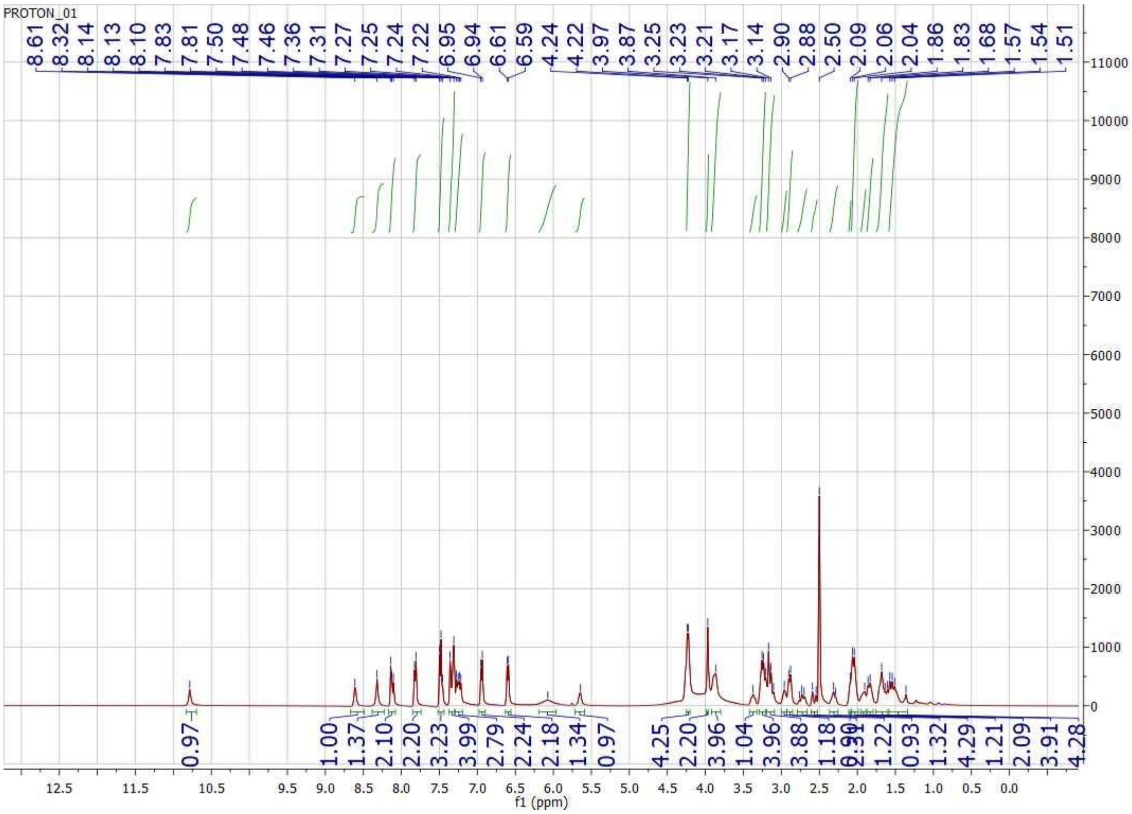

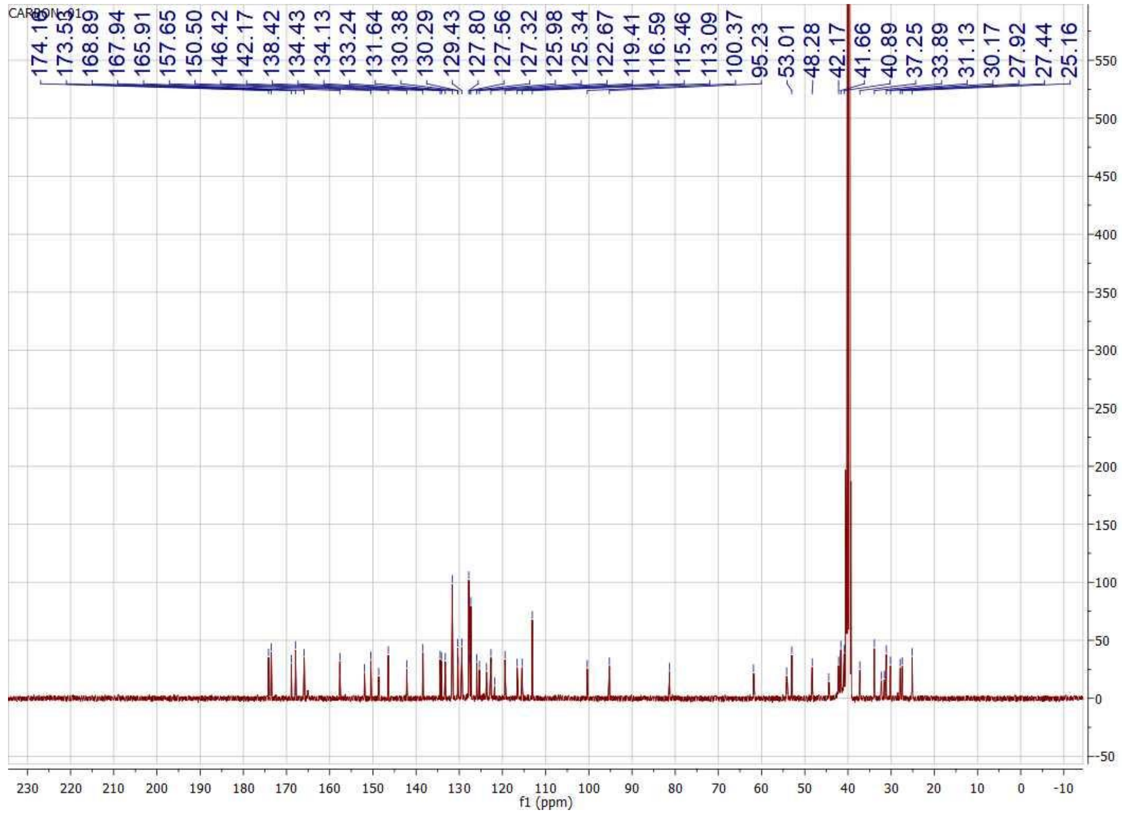
CARBON_01
174.16
173.55
168.89
167.94
165.91
157.65
150.50
146.42
142.17
138.42
134.43
134.13
133.24
131.64
130.38
130.29
129.43
127.80
127.56
127.32
125.98
125.34
122.67
119.41
116.59
115.46
113.09
100.37
95.23
53.01
48.28
42.17
41.66
40.89
37.25
33.89
31.13
30.17
27.92
27.44
25.16
f1 (ppm)

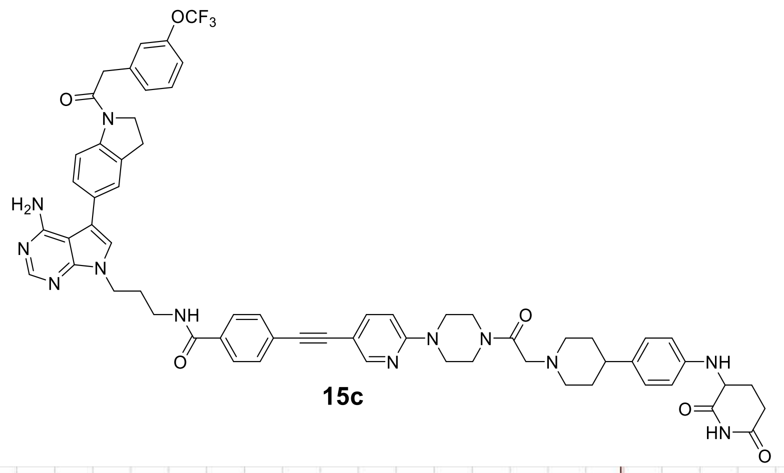


**15c**

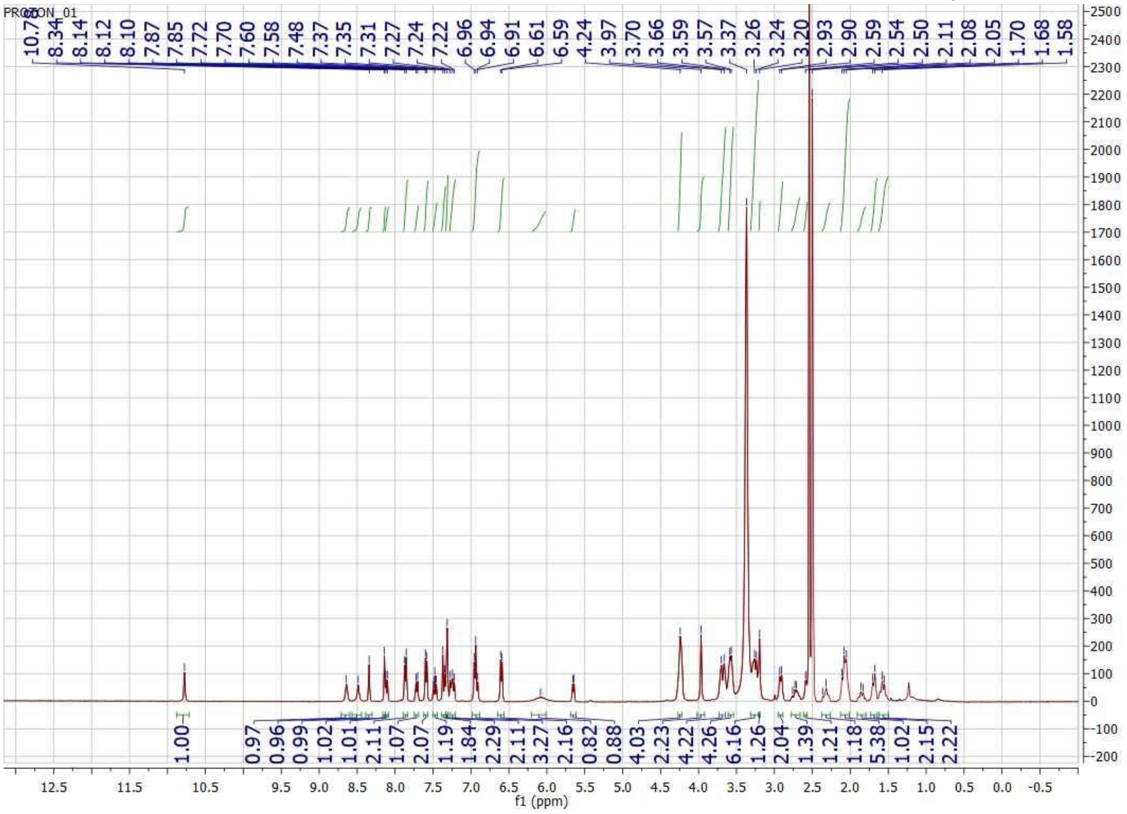

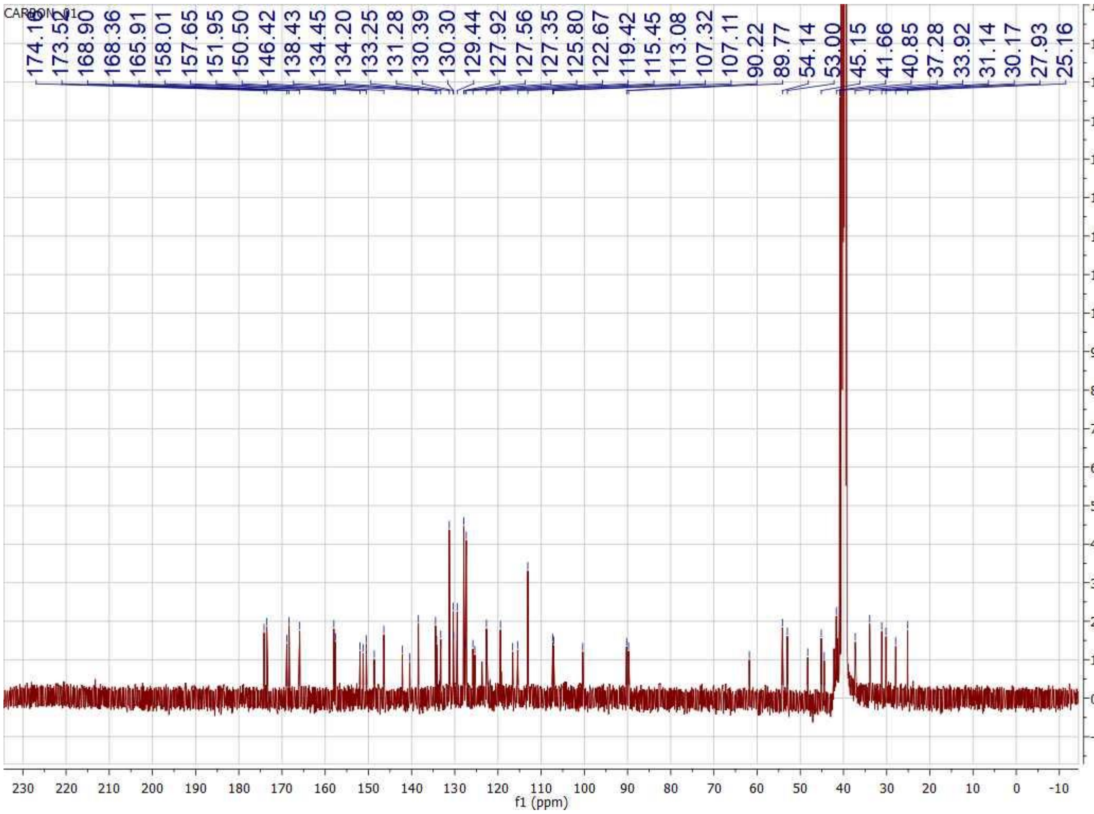
CARBON_01
174.16
173.52
168.90
168.36
165.91
158.01
157.65
151.95
150.50
146.42
138.43
134.45
134.20
133.25
131.28
130.39
130.30
129.44
127.92
127.56
127.35
125.80
122.67
119.42
115.45
113.08
107.32
107.11
90.22
89.77
54.14
53.00
45.15
41.66
40.85
37.28
33.92
31.14
30.17
27.93
25.16
f1 (ppm)

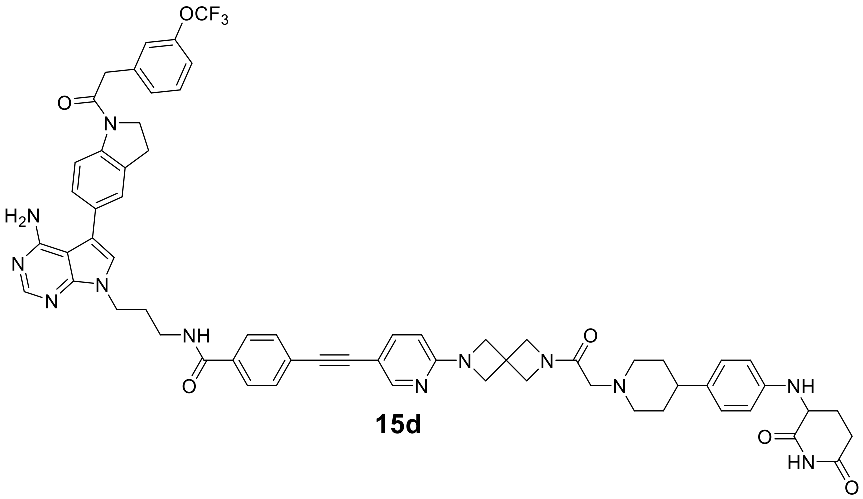

**15d**

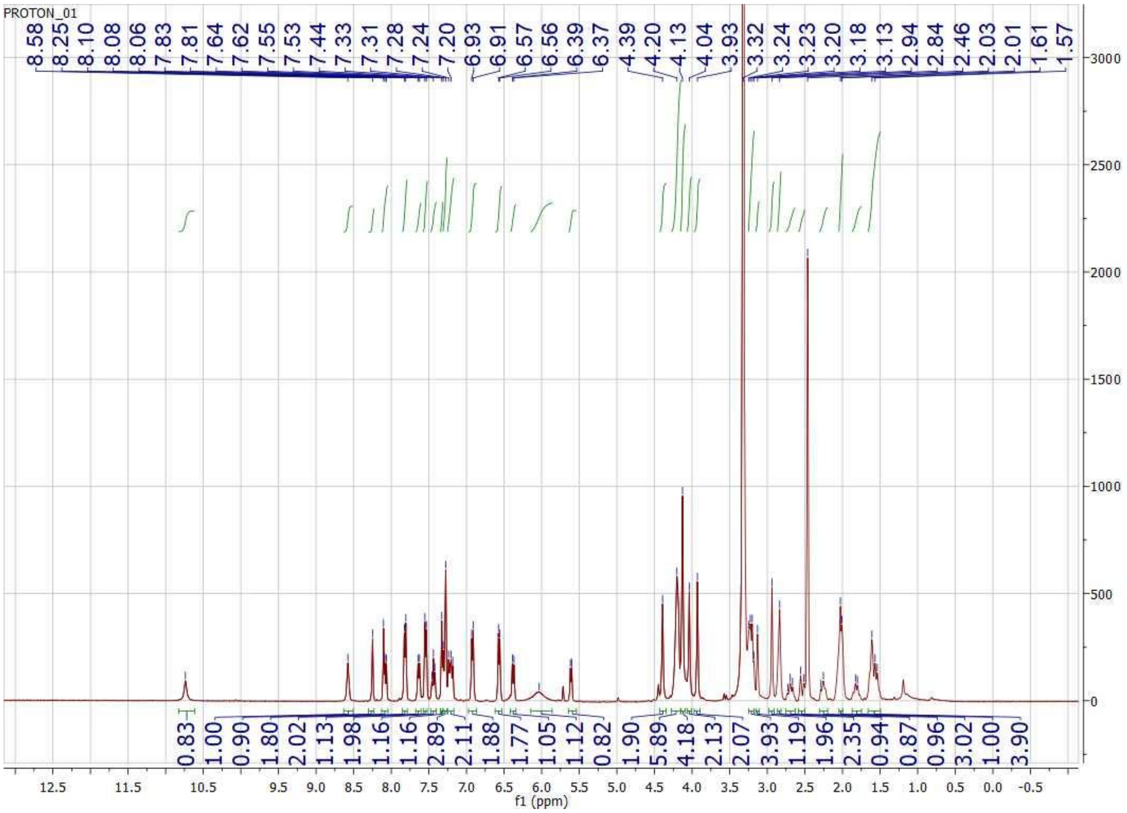

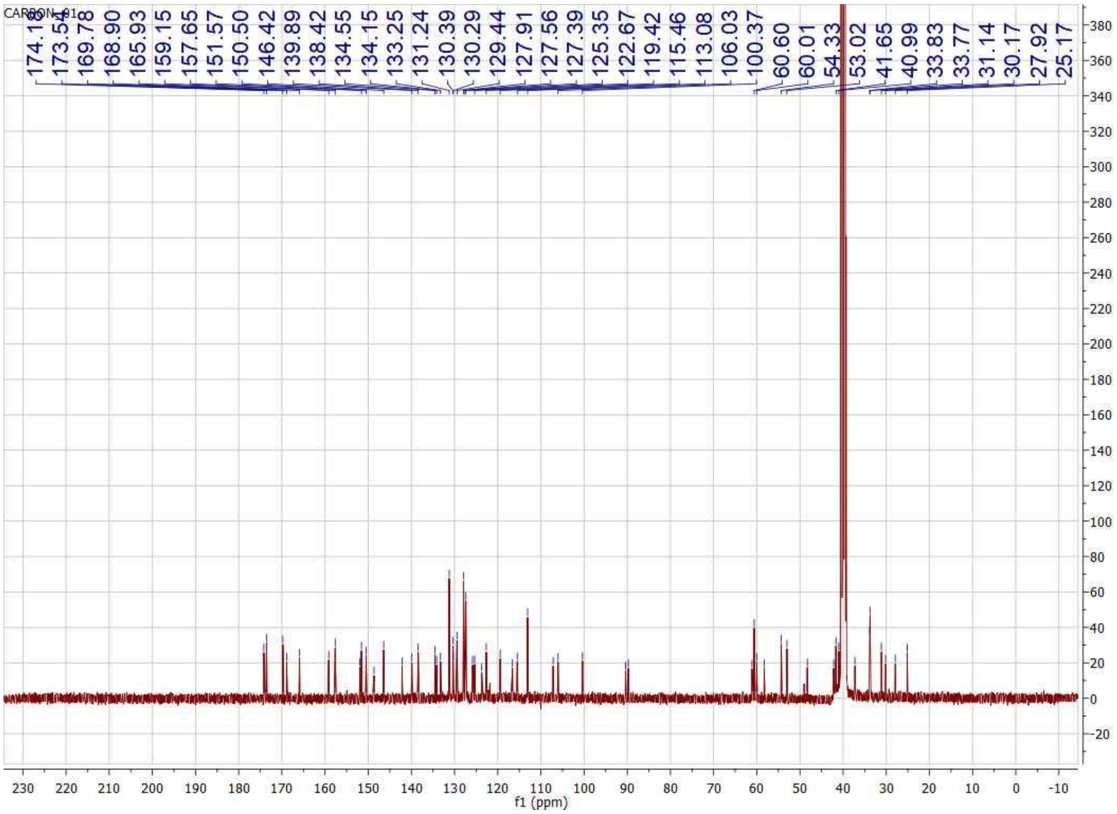
CARBON_01
174.16
173.54
169.78
168.90
165.93
159.15
157.65
151.57
150.50
146.42
139.89
138.42
134.55
134.15
133.25
131.24
130.39
130.29
129.44
127.91
127.56
127.39
125.35
122.67
119.42
115.46
113.08
106.03
100.37
60.60
60.01
54.33
53.02
41.65
40.99
33.83
33.77
31.14
30.17
27.92
25.17
380
360
340
320
300
280
260
240
220
200
180
160
140
120
100
80
60
40
20
0
-20
230 220 210 200 190 180 170 160 150 140 130 120 110 100 90 80 70 60 50 40 30 20 10 0 -10
f1 (ppm)

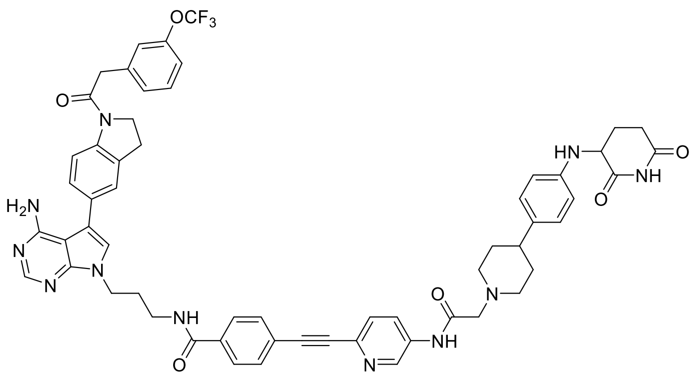

**15e**

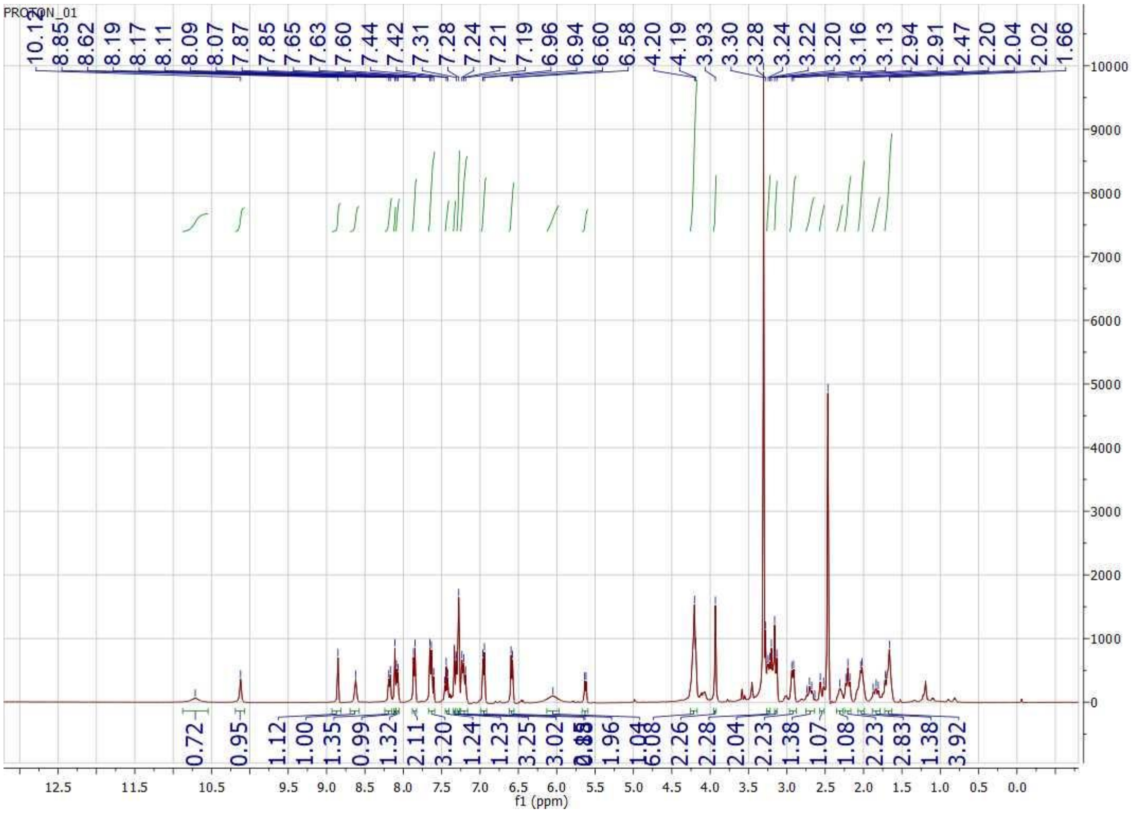
PROTON_01
10.17 8.85 8.62 8.19 8.17 8.11 8.09 8.07 7.87 7.85 7.65 7.63 7.60 7.44 7.42 7.31 7.28 7.24 7.21 7.19 6.96 6.94 6.60 6.58 4.20 4.19 3.93 3.30 3.28 3.24 3.22 3.20 3.16 3.13 2.94 2.91 2.47 2.20 2.04 2.02 1.66
0.72 0.95 1.12 1.00 1.35 0.99 1.32 2.11 3.20 1.24 1.23 3.25 3.02 1.96 1.04 6.08 2.26 2.28 2.04 2.23 1.38 1.07 1.08 2.23 2.83 1.38 3.92
f1 (ppm)

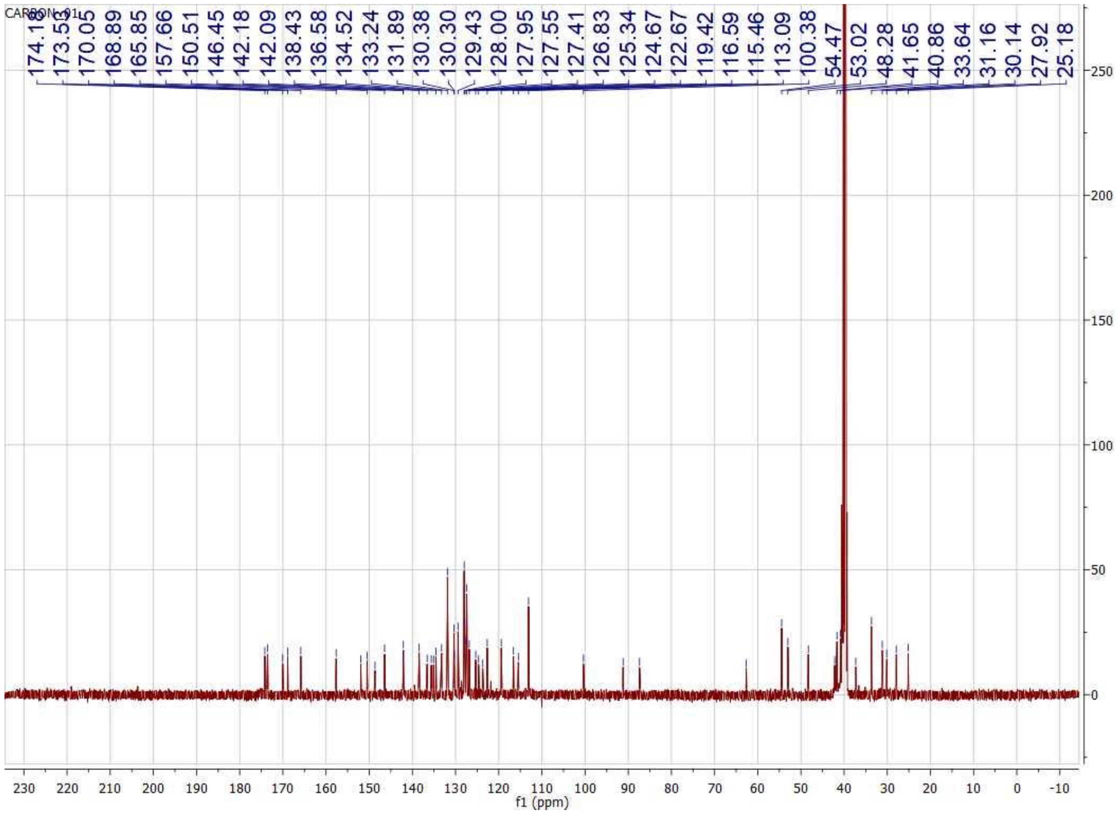
CARBON_01
174.15
173.55
170.05
168.89
165.85
157.66
150.51
146.45
142.18
142.09
138.43
136.58
134.52
133.24
131.89
130.38
130.30
129.43
128.00
127.95
127.55
127.41
126.83
125.34
124.67
122.67
119.42
116.59
115.46
113.09
100.38
54.47
53.02
48.28
41.65
40.86
33.64
31.16
30.14
27.92
25.18
f1 (ppm)

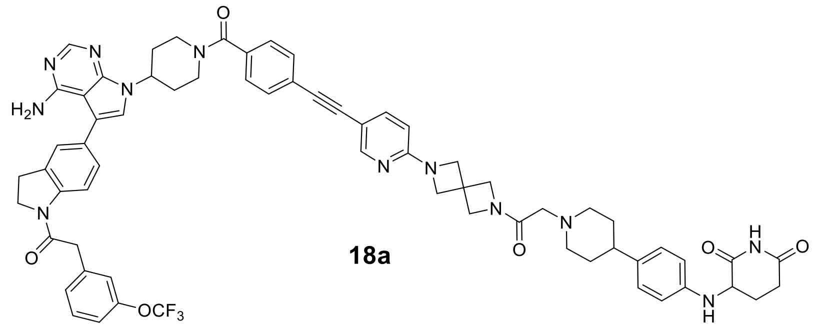

**18a**

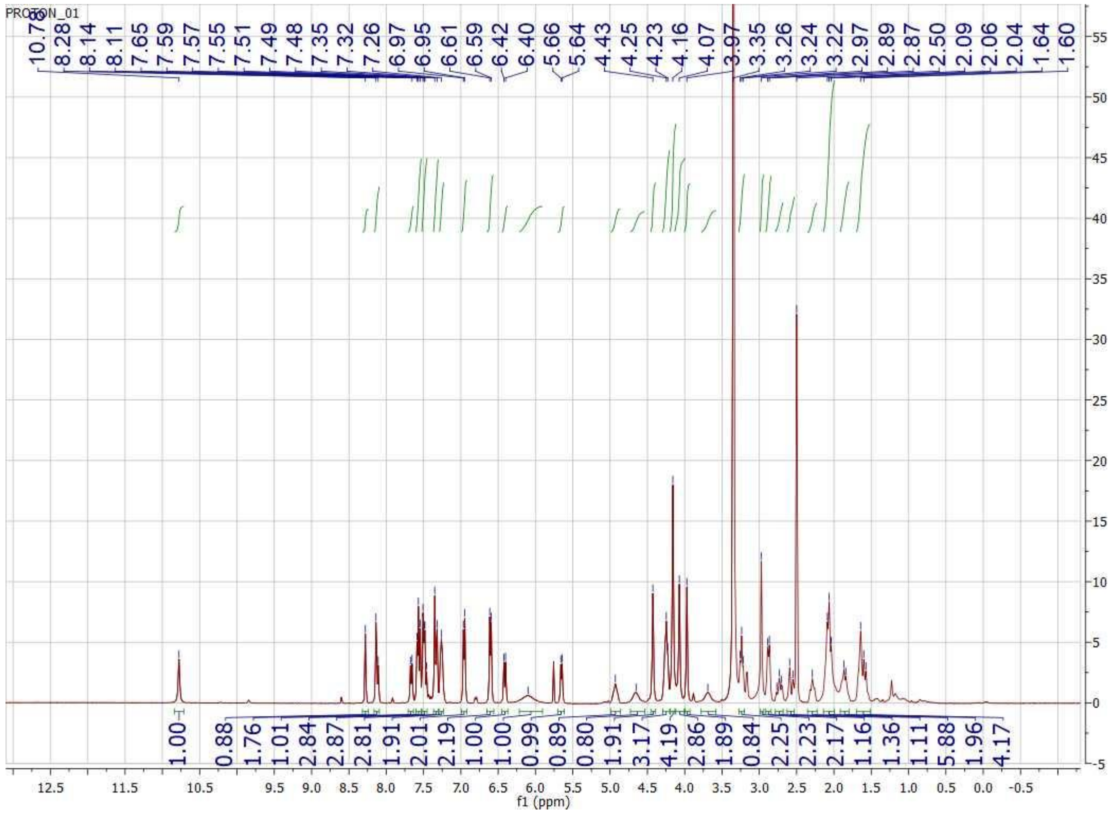
PROTON_01
10.70 8.28 8.14 8.11 7.65 7.59 7.57 7.55 7.51 7.49 7.48 7.35 7.32 7.26 6.97 6.95 6.61 6.59 6.42 6.40 5.66 5.64 4.43 4.25 4.23 4.16 4.07 3.97 3.35 3.26 3.24 3.22 2.97 2.89 2.87 2.50 2.09 2.06 2.04 1.64 1.60
1.00 0.88 1.76 1.01 2.84 2.87 2.81 1.91 2.01 2.19 1.00 1.00 0.99 0.89 0.80 1.91 3.17 4.19 2.86 1.89 0.84 2.25 2.23 2.17 1.16 1.36 1.11 5.88 1.96 4.17
f1 (ppm)

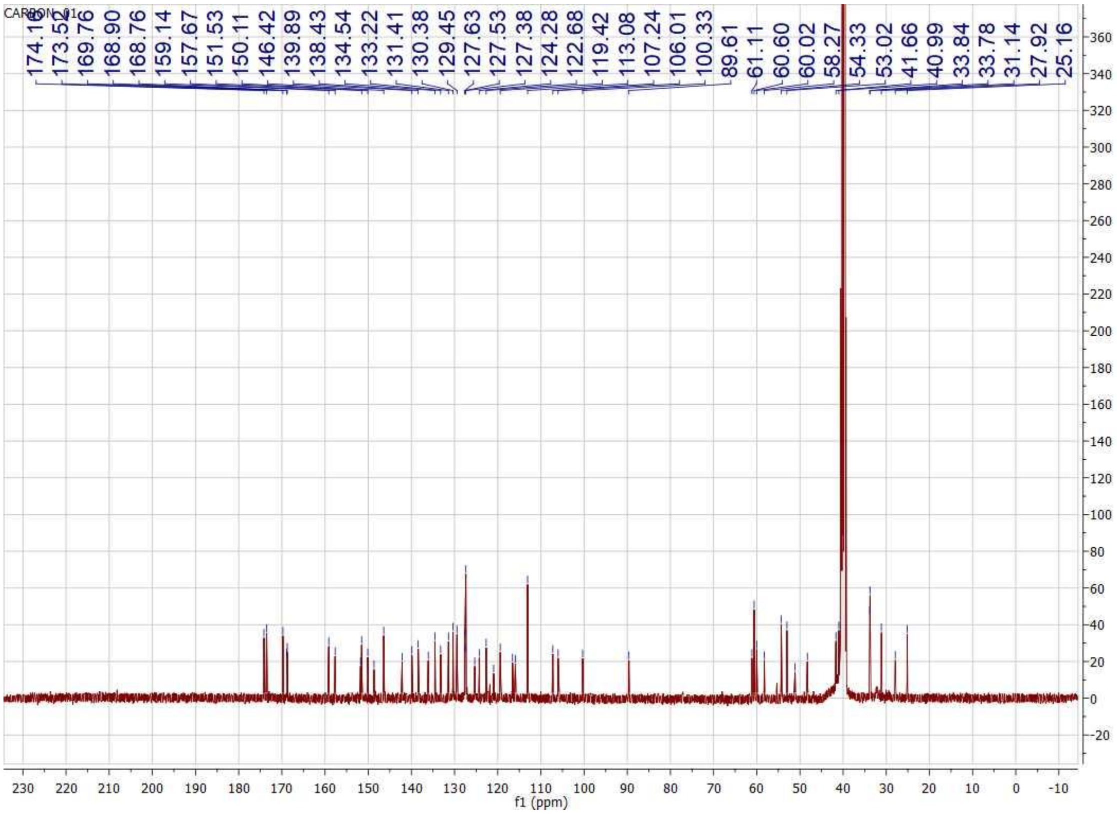
CARBON_01
174.15
173.52
169.76
168.90
168.76
159.14
157.67
151.53
150.11
146.42
139.89
138.43
134.54
133.22
131.41
130.38
129.45
127.63
127.53
127.38
124.28
122.68
119.42
113.08
107.24
106.01
100.33
89.61
61.11
60.60
60.02
58.27
54.33
53.02
41.66
40.99
33.84
33.78
31.14
27.92
25.16
f1 (ppm)

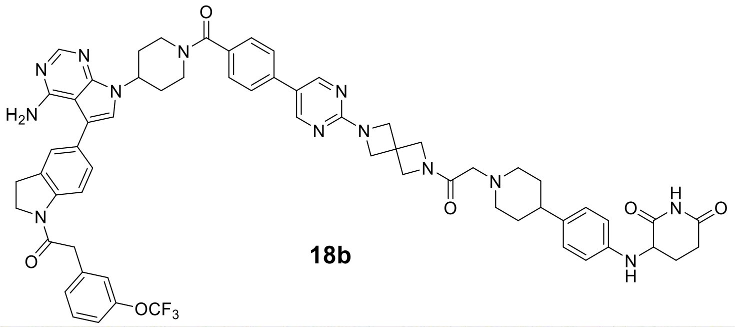
N
N
N
H2N
N
O
N
O
OCF3
O
N
N
N
N
O
N
NH
O
O
N
H

**18b**

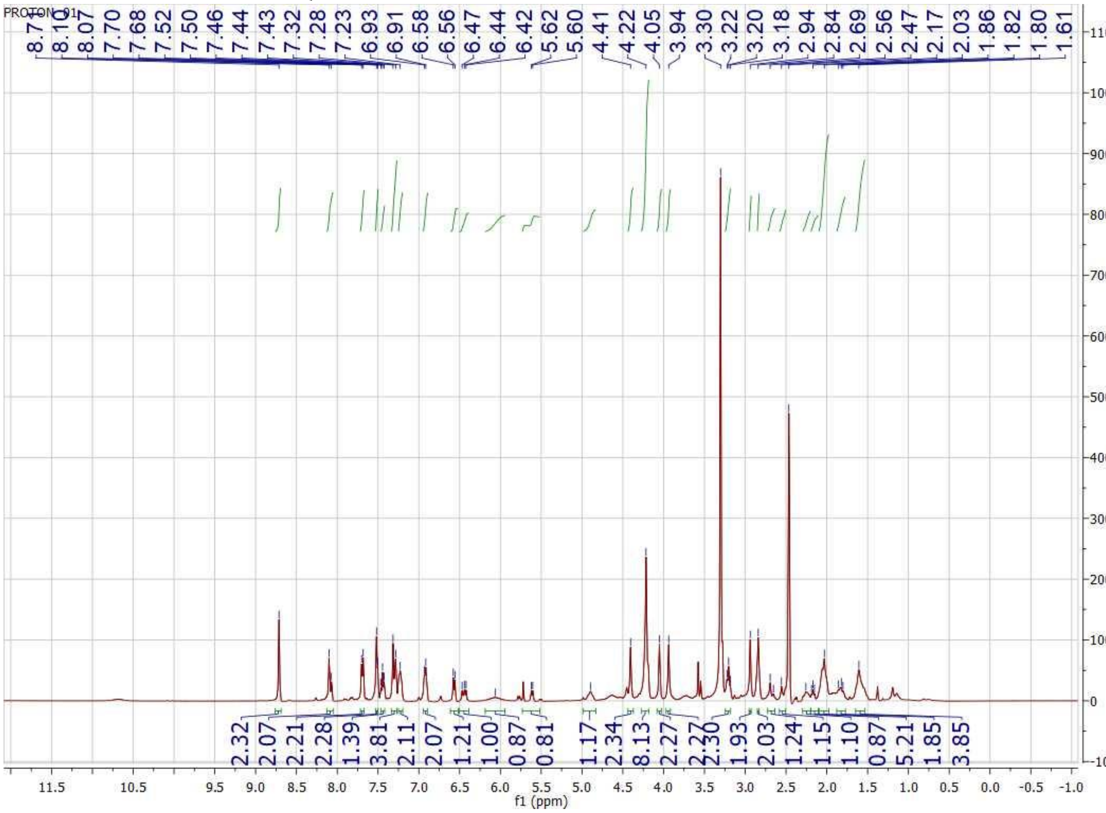
PROTON_01
8.71
8.10
8.07
7.70
7.68
7.52
7.50
7.46
7.44
7.43
7.32
7.28
7.23
6.93
6.91
6.58
6.56
6.47
6.44
6.42
5.62
5.60
4.41
4.22
4.05
3.94
3.30
3.22
3.20
3.18
2.94
2.84
2.69
2.56
2.47
2.17
2.03
1.86
1.82
1.80
1.61
2.32
2.07
2.21
2.28
1.39
3.81
2.11
2.07
1.21
1.00
0.87
0.81
0.81
1.17
2.34
8.13
2.27
2.27
2.30
1.93
2.03
1.24
1.15
1.10
0.87
5.21
1.85
3.85
f1 (ppm)

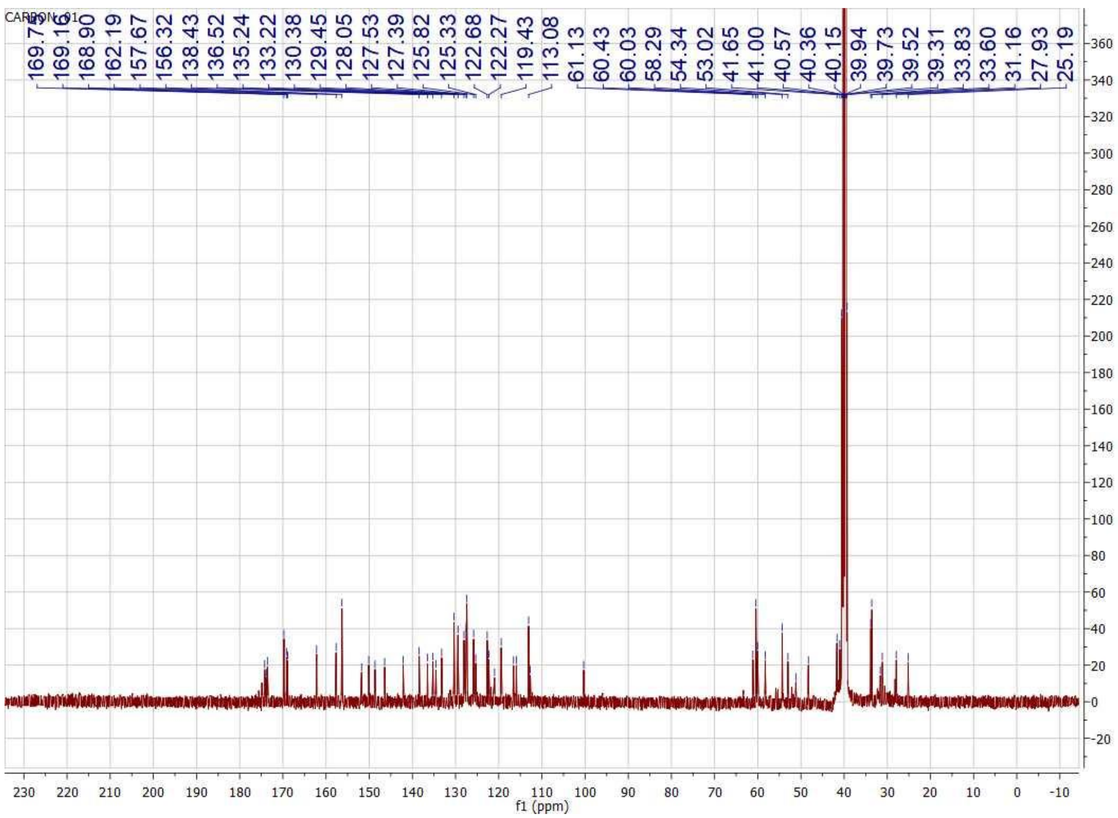
CARBON_01
169.7
169.16
168.90
162.19
157.67
156.32
138.43
136.52
135.24
133.22
130.38
129.45
128.05
127.53
127.39
125.82
125.33
122.68
122.27
119.43
113.08
61.13
60.43
60.03
58.29
54.34
53.02
41.65
41.00
40.57
40.36
40.15
39.94
39.73
39.52
39.31
33.83
33.60
31.16
27.93
25.19
f1 (ppm)

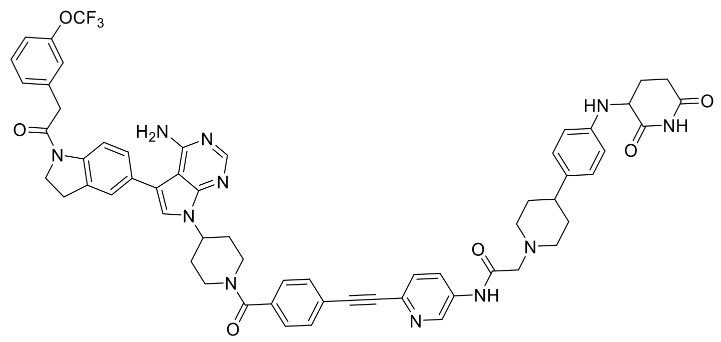

**18c**

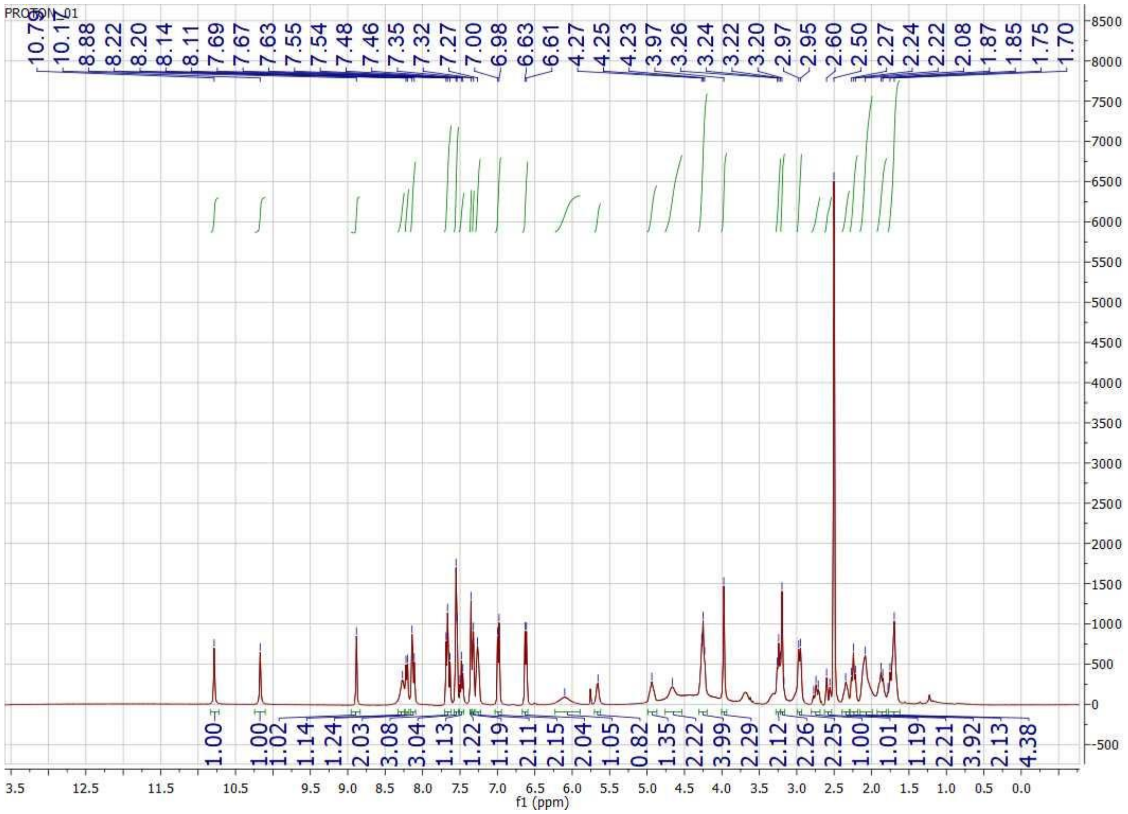

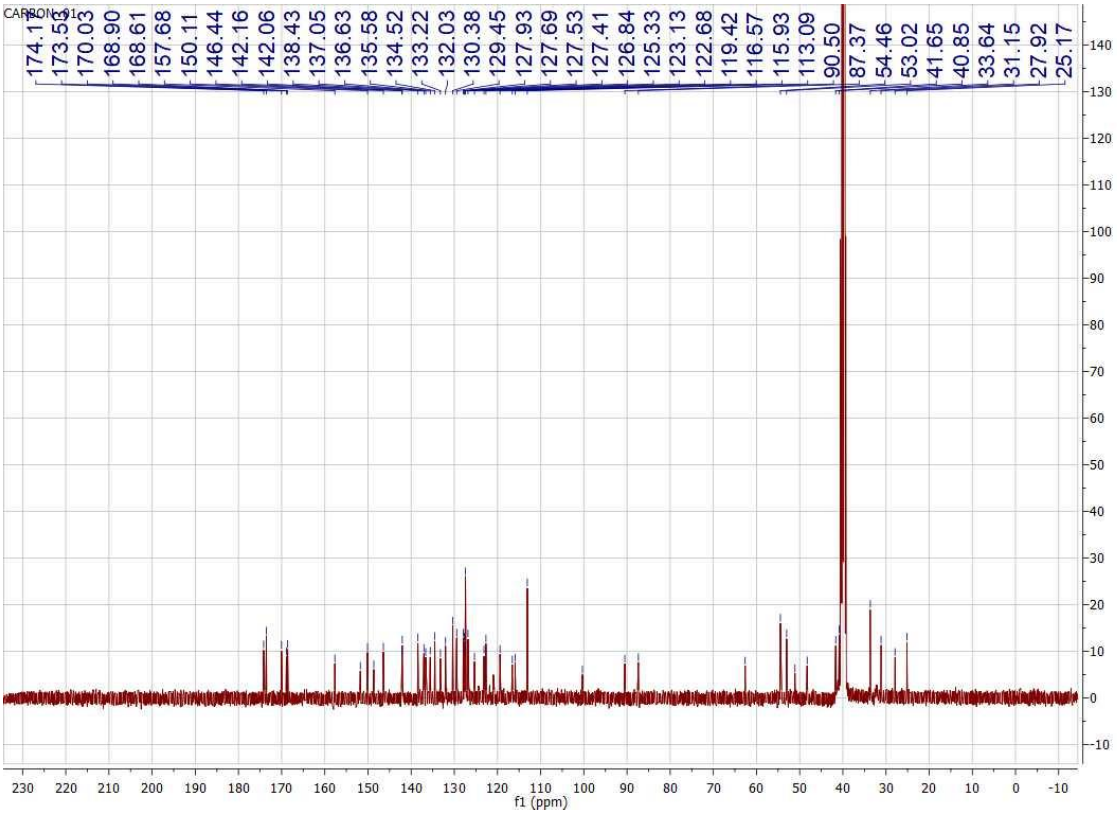
CARBON_01
174.17
173.55
170.03
168.90
168.61
157.68
150.11
146.44
142.16
142.06
138.43
137.05
136.63
135.58
134.52
133.22
132.03
130.38
129.45
127.93
127.69
127.53
127.41
126.84
125.33
123.13
122.68
119.42
116.57
115.93
113.09
90.50
87.37
54.46
53.02
41.65
40.85
33.64
31.15
27.92
25.17
f1 (ppm)

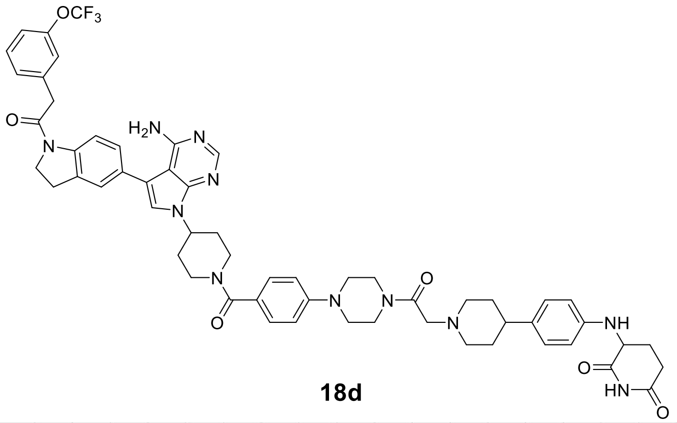
OCF3
H2N
N
O
NH
HN

**18d**

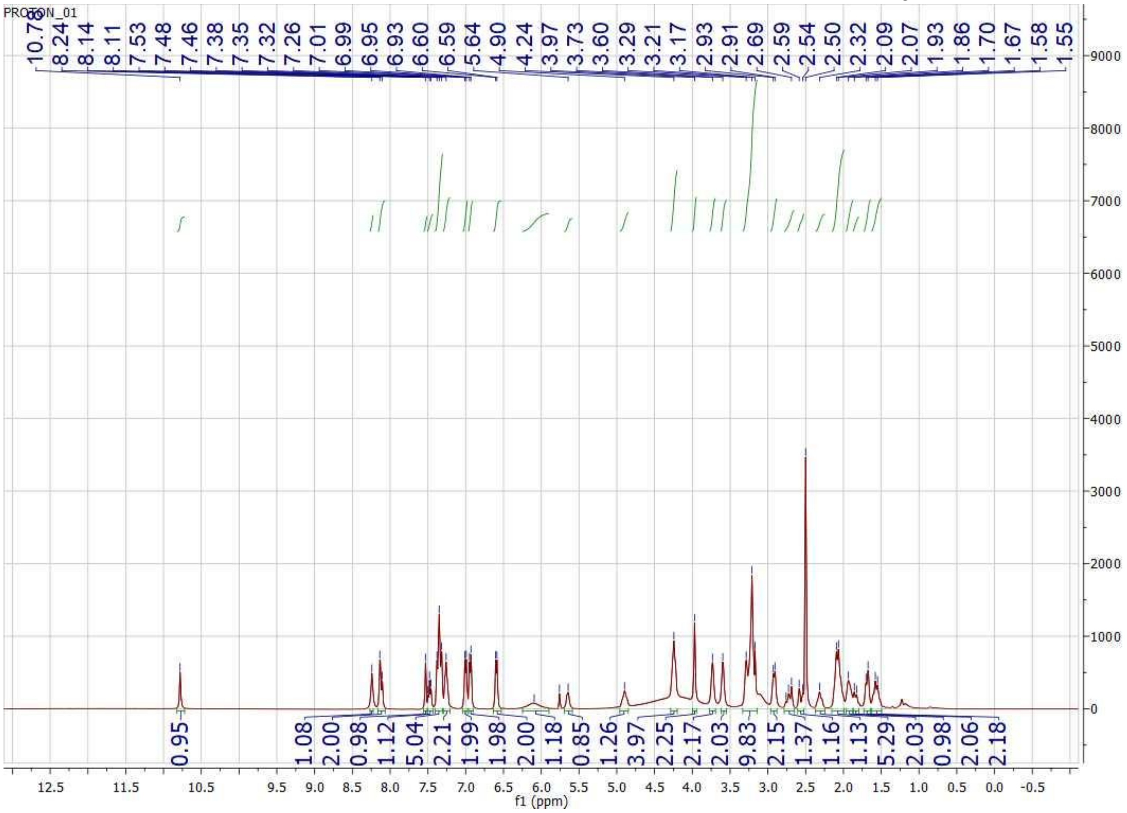
PROTON_01
10.70
8.24
8.14
8.11
7.53
7.48
7.46
7.38
7.35
7.32
7.26
7.01
6.99
6.95
6.93
6.60
6.59
5.64
4.90
4.24
3.97
3.73
3.60
3.29
3.21
3.17
2.93
2.91
2.69
2.59
2.54
2.50
2.32
2.09
2.07
1.93
1.86
1.70
1.67
1.58
1.55
0.95
1.08
2.00
0.98
1.12
5.04
2.21
1.99
1.98
2.00
1.18
0.85
1.26
3.97
2.25
2.17
2.03
9.83
2.15
1.37
1.16
1.13
5.29
2.03
0.98
2.06
2.18
f1 (ppm)

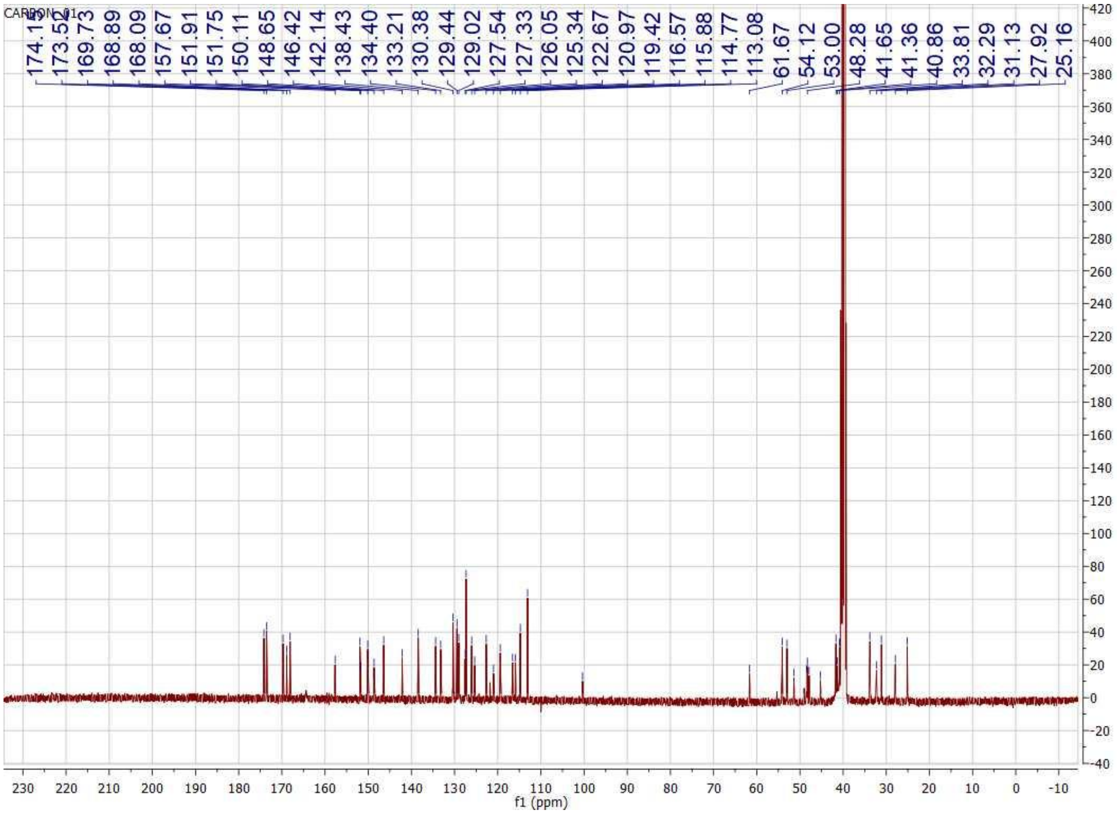

CARBON_01
174.19
173.52
169.73
168.89
168.09
157.67
151.91
151.75
150.11
148.65
146.42
142.14
138.43
134.40
133.21
130.38
129.44
129.02
127.54
127.33
126.05
125.34
122.67
120.97
119.42
116.57
115.88
114.77
113.08
61.67
54.12
53.00
48.28
41.65
41.36
40.86
33.81
32.29
31.13
27.92
25.16
f1 (ppm)

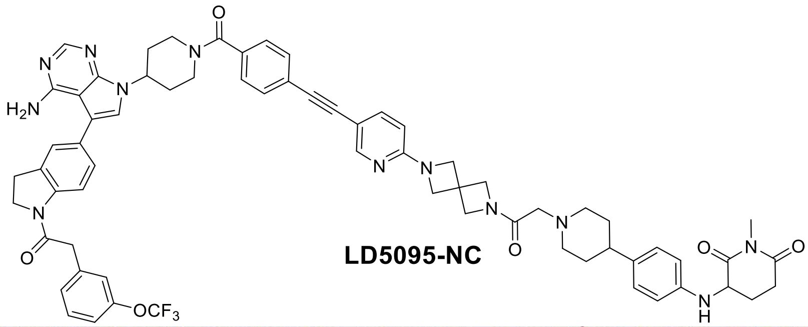
LD5095-NC

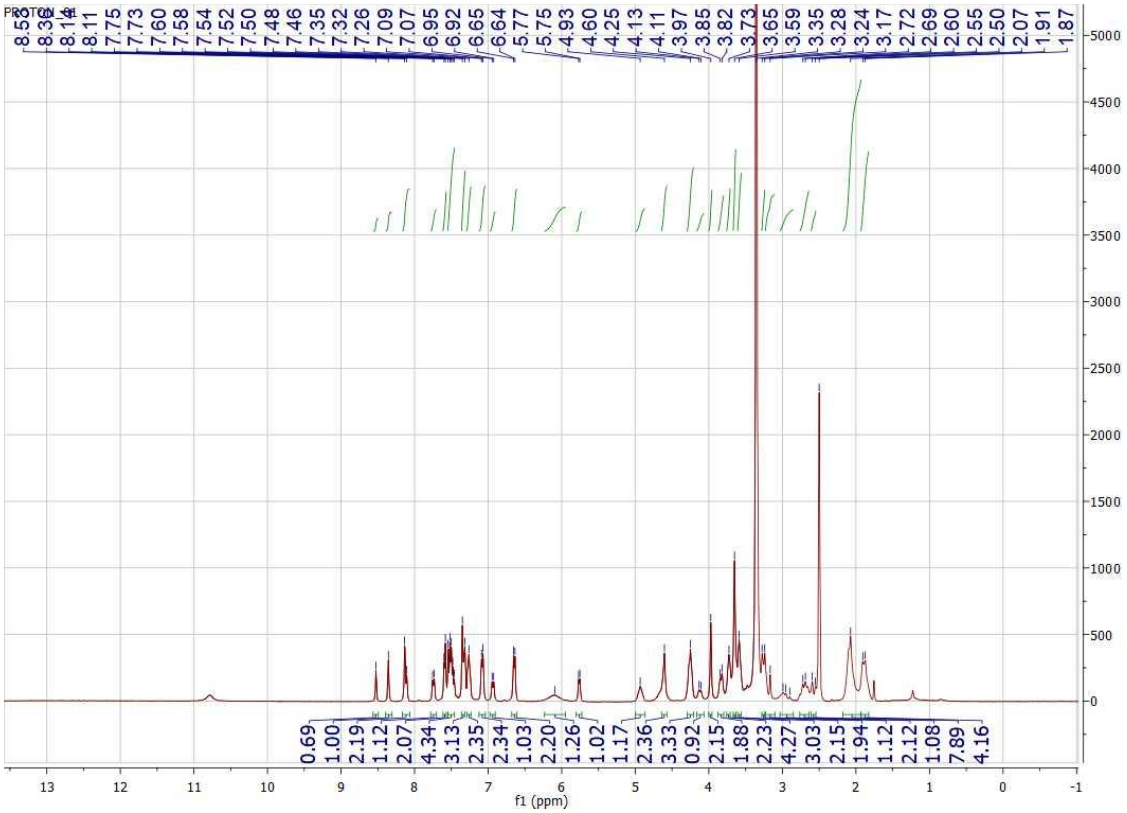

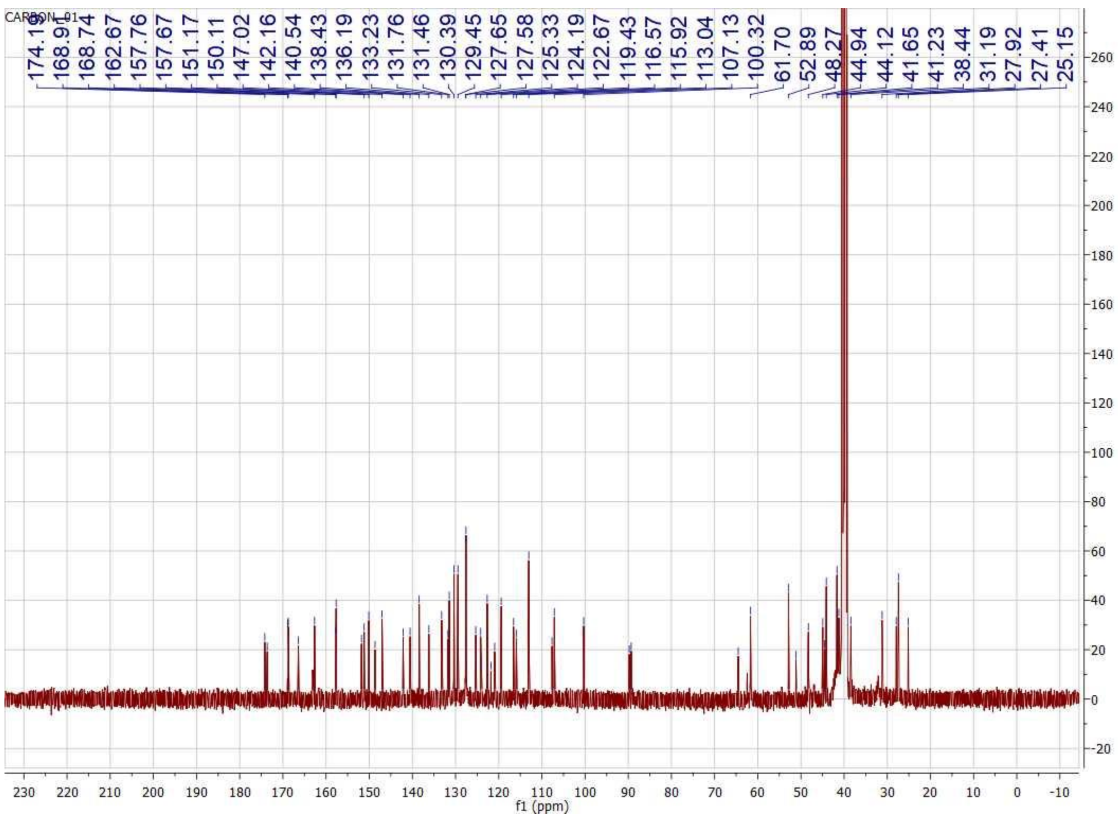
CARBON_01
174.1
168.91
168.74
162.67
157.76
157.67
151.17
150.11
147.02
142.16
140.54
138.43
136.19
133.23
131.76
131.46
130.39
129.45
127.65
127.58
125.33
124.19
122.67
119.43
116.57
115.92
113.04
107.13
100.32
61.70
52.89
48.27
44.94
44.12
41.65
41.23
38.44
31.19
27.92
27.41
25.15
f1 (ppm)

### HPLC chromatography of representative compounds

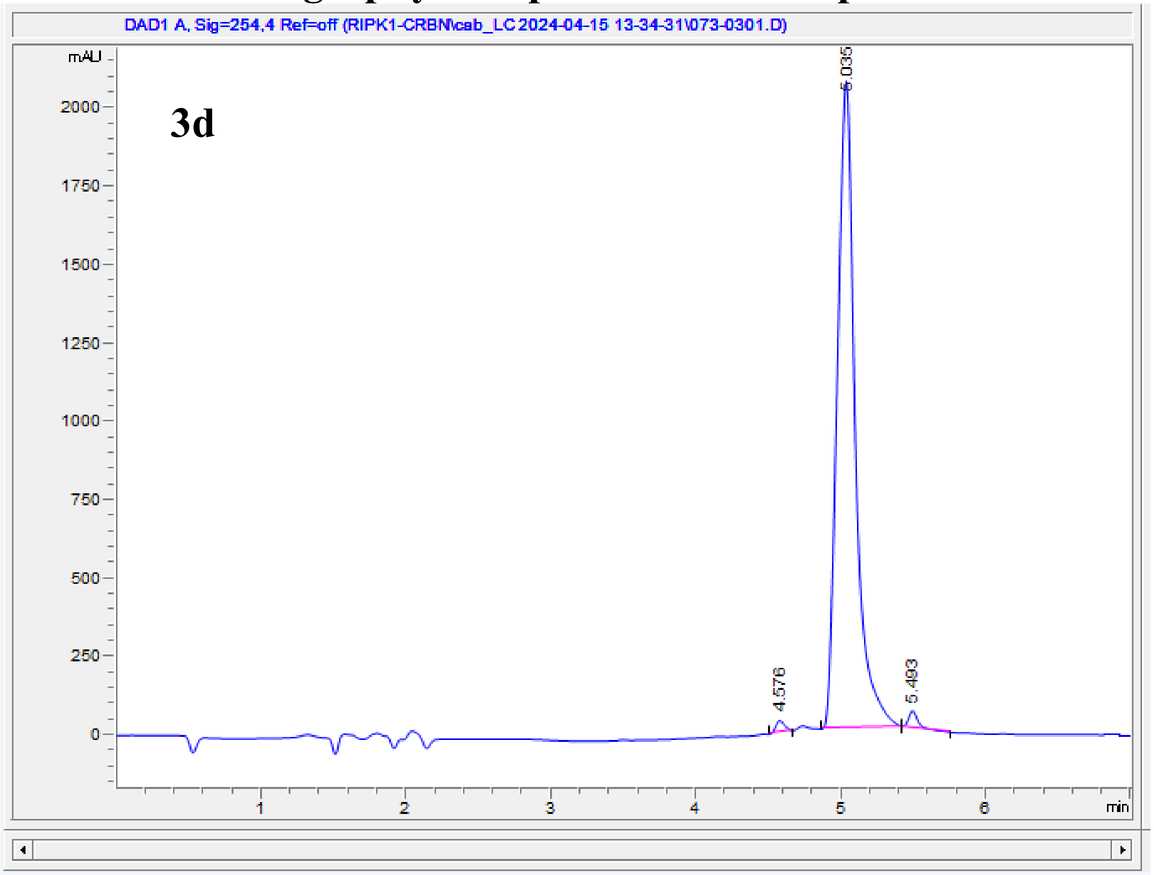


| # | Time | Area | Height | Width | Area% | Symmetry |
|---|---|---|---|---|---|---|
| 1 | 4.576 | 157.8 | 37.6 | 0.0649 | 0.874 | 0.809 |
| 2 | 5.035 | 17658.6 | 2062.5 | 0.1339 | 97.869 | 0.88 |
| 3 | 5.493 | 226.8 | 52.9 | 0.066 | 1.257 | 0.68 |

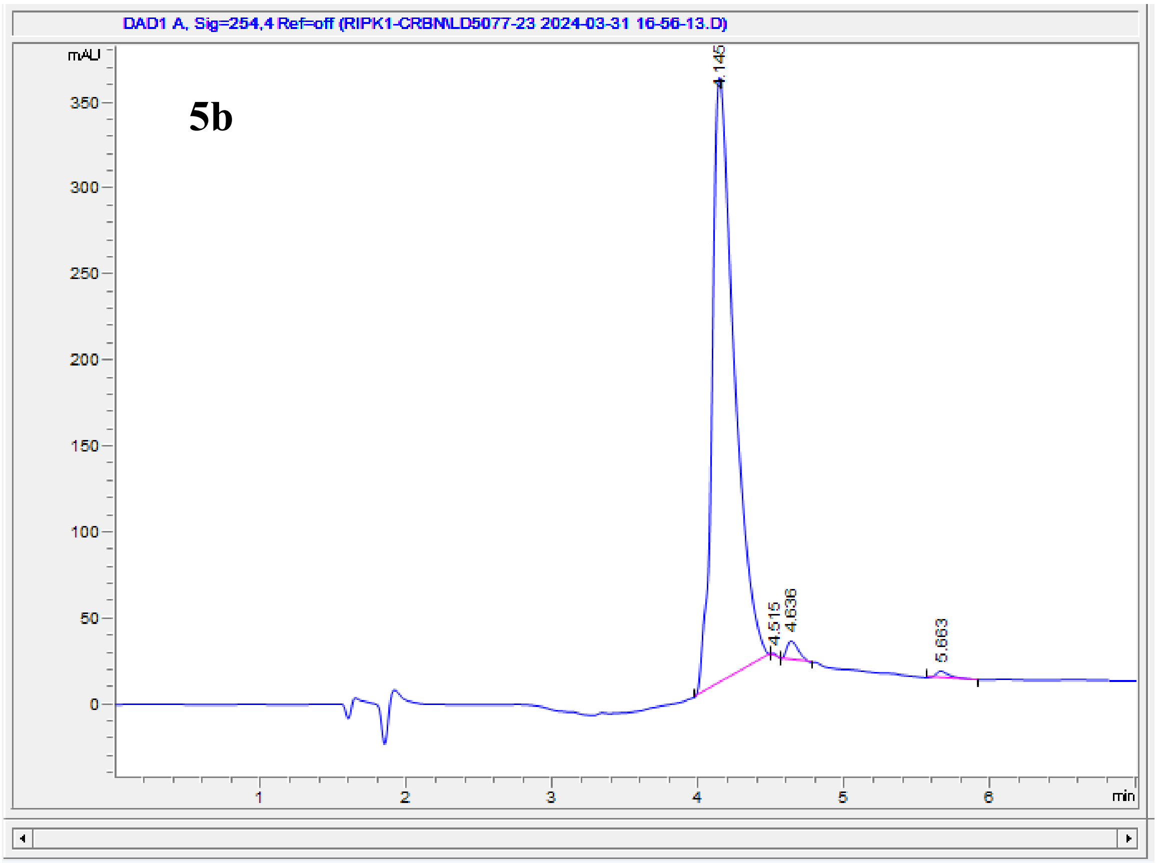


| # | Time | Area | Height | Width | Area% | Symmetry |
|---|---|---|---|---|---|---|
| 1 | 4.145 | 3726.9 | 351.6 | 0.1557 | 97.596 | 0.472 |
| 2 | 4.515 | 3.1 | 1.2 | 0.0433 | 0.081 | 0.347 |
| 3 | 4.636 | 61 | 10.9 | 0.0872 | 1.597 | 0.57 |
| 4 | 5.663 | 27.7 | 4.2 | 0.0929 | 0.726 | 0.496 |

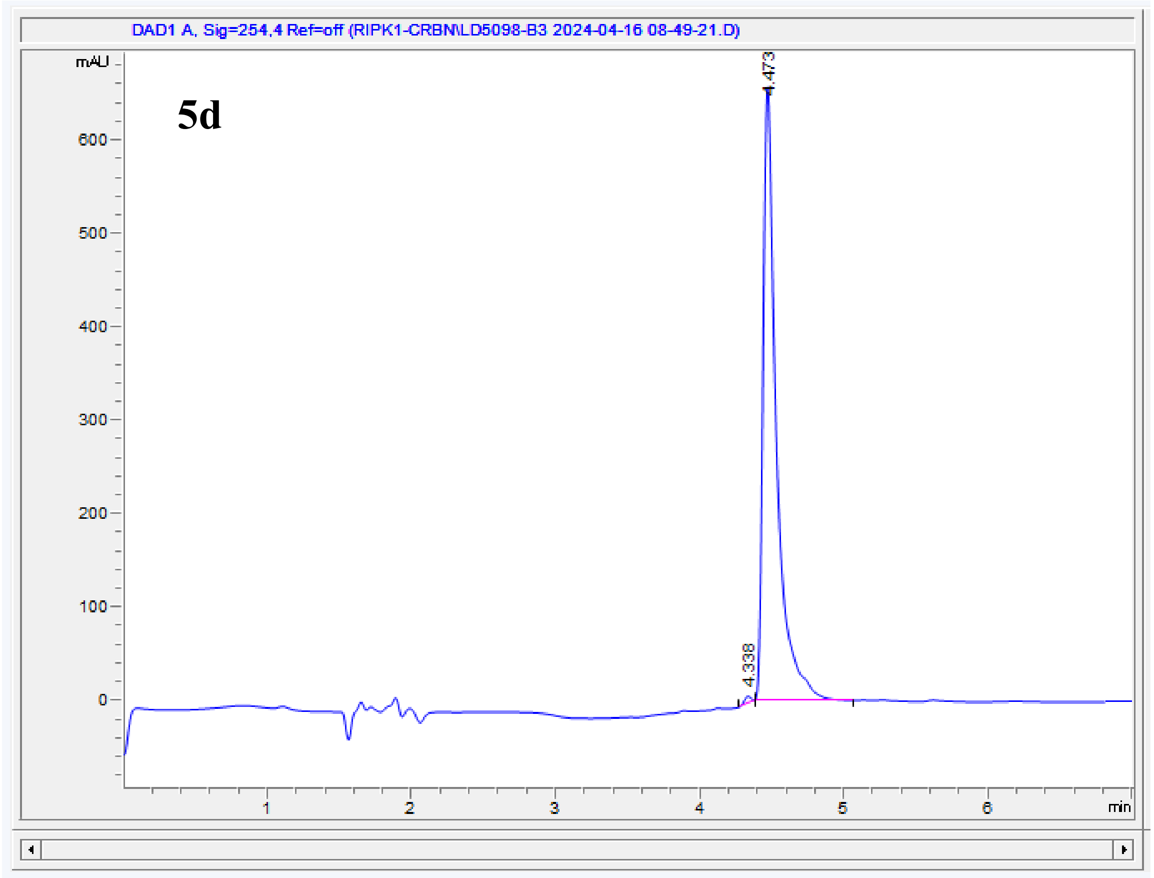


| # | Time | Area | Height | Width | Area% | Symmetry |
|---|---|---|---|---|---|---|
| 1 | 4.338 | 23 | 7.2 | 0.0514 | 0.539 | 1.424 |
| 2 | 4.473 | 4247.1 | 659.5 | 0.0931 | 99.461 | 0.449 |

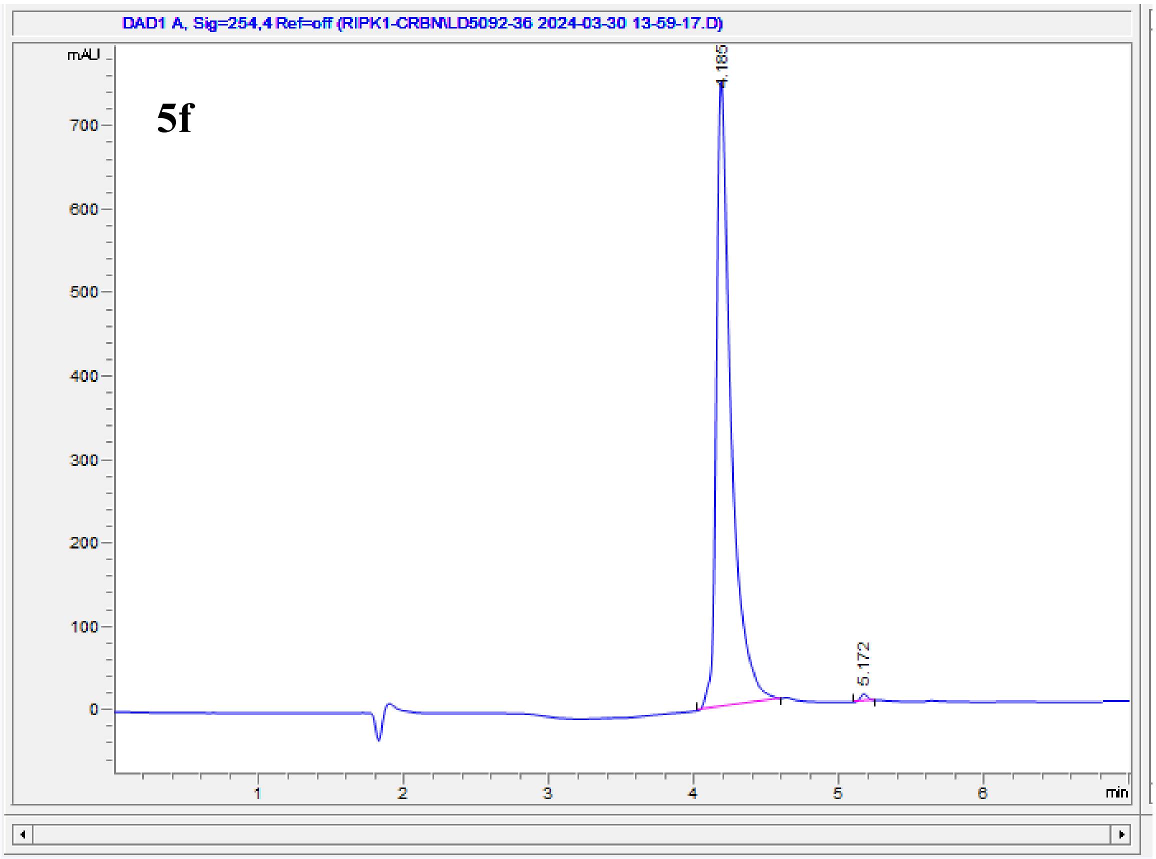


| # | Time | Area | Height | Width | Area% | Symmetry |
|---|---|---|---|---|---|---|
| 1 | 4.185 | 5392.6 | 754.9 | 0.1033 | 99.450 | 0.448 |
| 2 | 5.172 | 29.8 | 8.7 | 0.0542 | 0.550 | 0.945 |

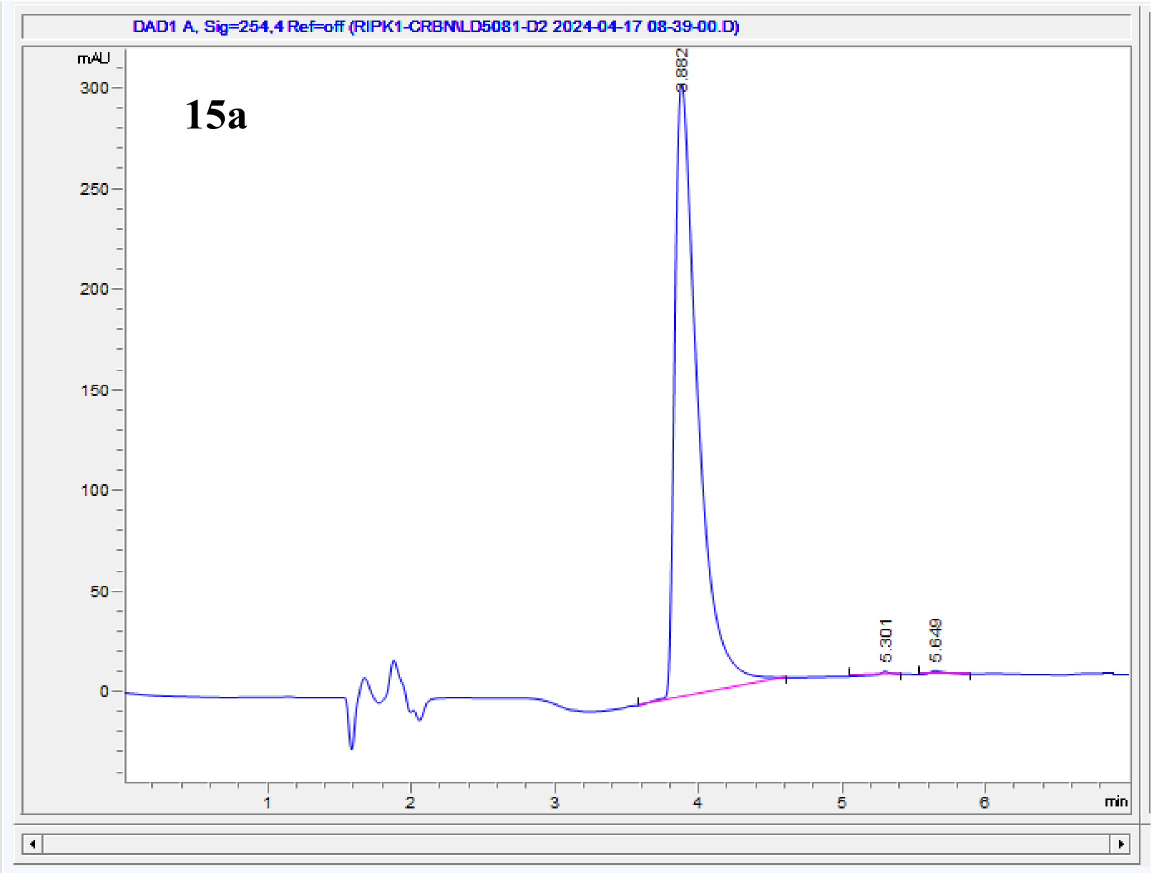


| # | Time | Area | Height | Width | Area% | Symmetry |
|---|---|---|---|---|---|---|
| 1 | 3.882 | 3436.4 | 305.5 | 0.1632 | 99.367 | 0.393 |
| 2 | 5.301 | 7.1 | 1.6 | 0.067 | 0.205 | 1.035 |
| 3 | 5.649 | 14.8 | 1.9 | 0.1059 | 0.427 | 0.459 |

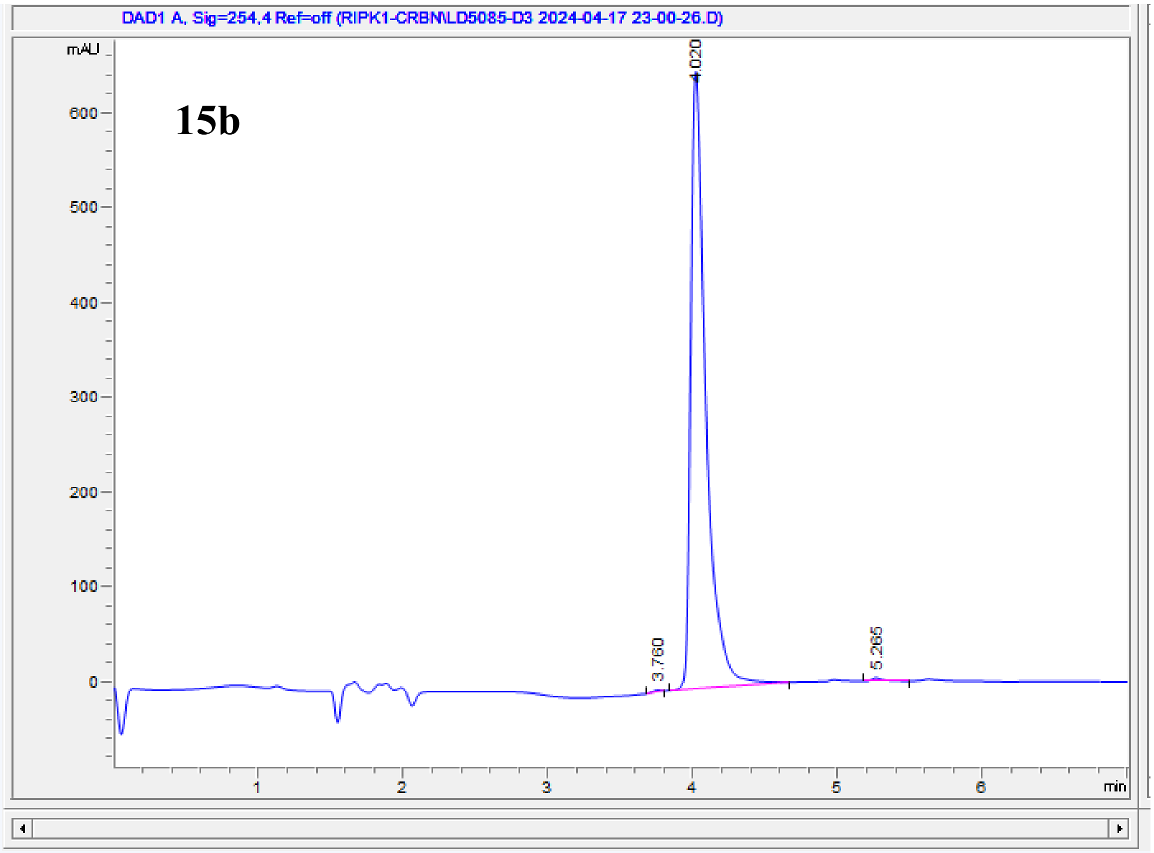


| # | Time | Area | Height | Width | Area% | Symmetry |
|---|---|---|---|---|---|---|
| 1 | 3.76 | 7.9 | 1.9 | 0.0665 | 0.166 | 1.935 |
| 2 | 4.02 | 4730 | 650.4 | 0.1068 | 99.404 | 0.447 |
| 3 | 5.265 | 20.5 | 3.6 | 0.0812 | 0.430 | 0.461 |

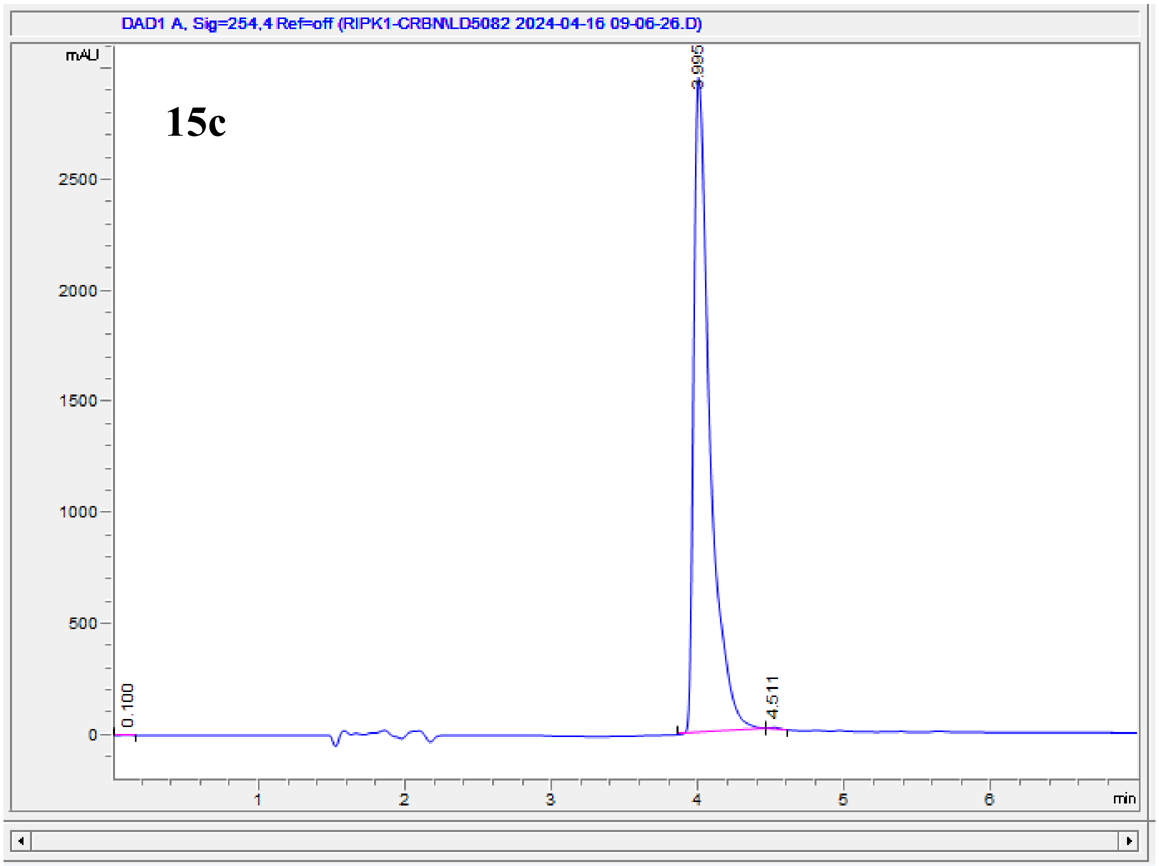


| # | Time | Area | Height | Width | Area% | Symmetry |
|---|---|---|---|---|---|---|
| 1 | 0.1 | 4.5E-1 | 1.2E-1 | 0.0577 | 0.002 | 1.488 |
| 2 | 3.995 | 22570.2 | 2944.4 | 0.1154 | 99.902 | 0.401 |
| 3 | 4.511 | 21.6 | 5.3 | 0.0634 | 0.096 | 0.554 |

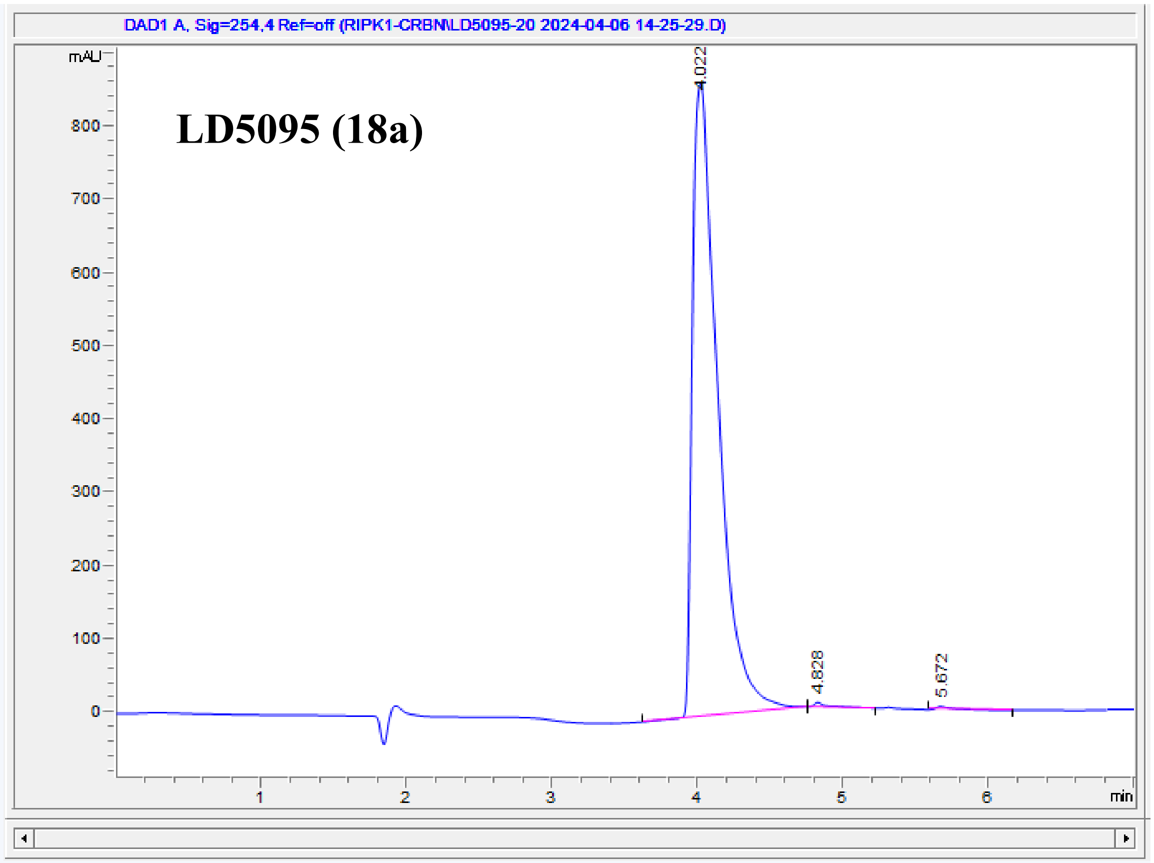


| # | Time | Area | Height | Width | Area% | Symmetry |
|---|---|---|---|---|---|---|
| 1 | 4.022 | 10577.2 | 866.9 | 0.1743 | 99.264 | 0.423 |
| 2 | 4.828 | 40 | 6.2 | 0.0878 | 0.375 | 0.305 |
| 3 | 5.672 | 38.4 | 4.7 | 0.11 | 0.360 | 0.293 |

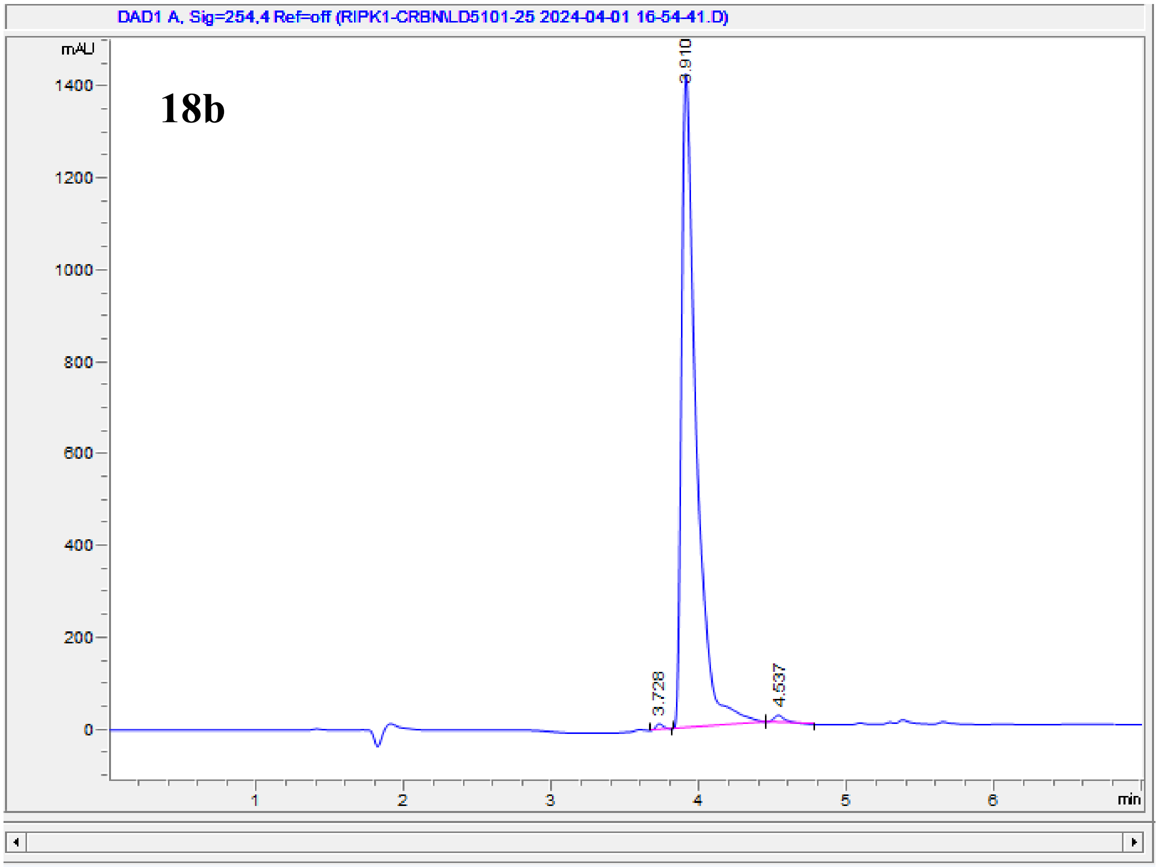


| # | Time | Area | Height | Width | Area% | Symmetry |
|---|---|---|---|---|---|---|
| 1 | 3.728 | 47.9 | 13.1 | 0.0546 | 0.476 | 0.76 |
| 2 | 3.91 | 9935.7 | 1424.2 | 0.0994 | 98.663 | 0.398 |
| 3 | 4.537 | 86.8 | 15.7 | 0.0804 | 0.862 | 0.582 |

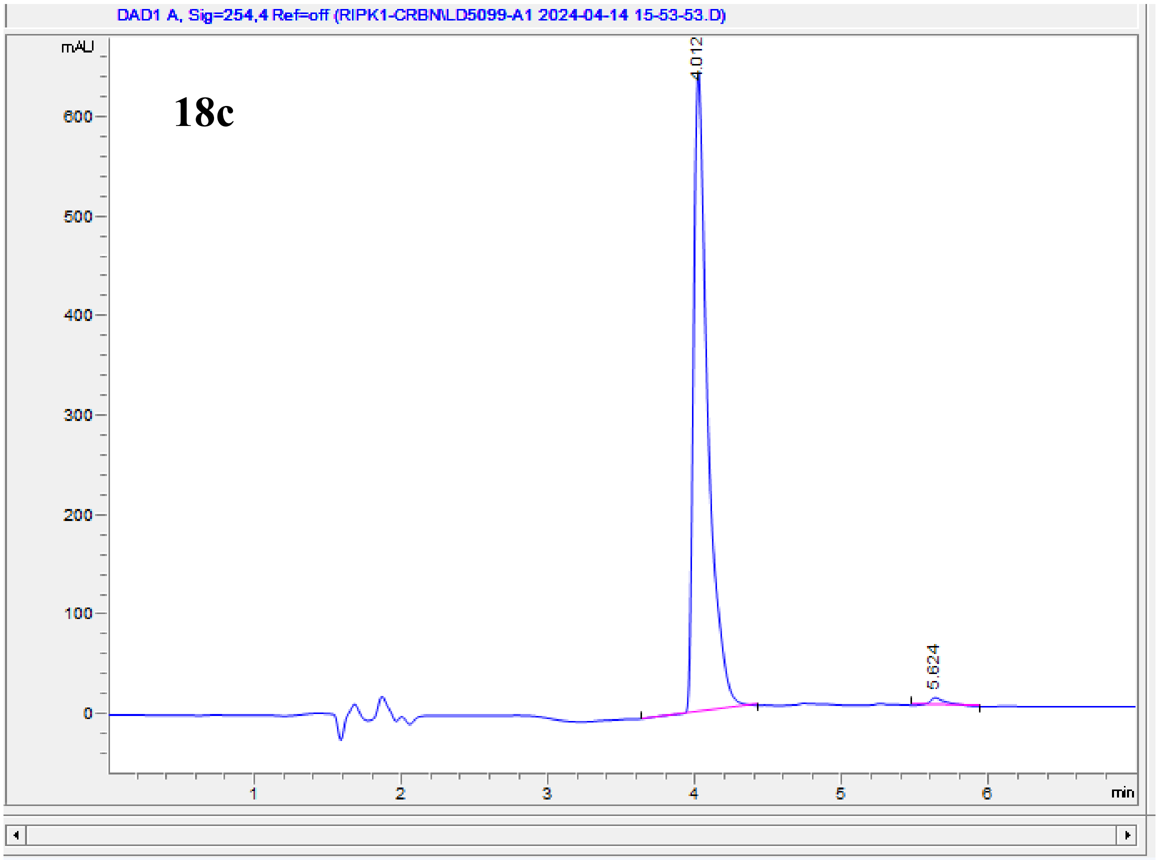


| # | Time | Area | Height | Width | Area% | Symmetry |
|---|---|---|---|---|---|---|
| 1 | 4.012 | 4386.2 | 643.6 | 0.1015 | 98.483 | 0.449 |
| 2 | 5.624 | 67.6 | 8 | 0.1141 | 1.517 | 0.463 |

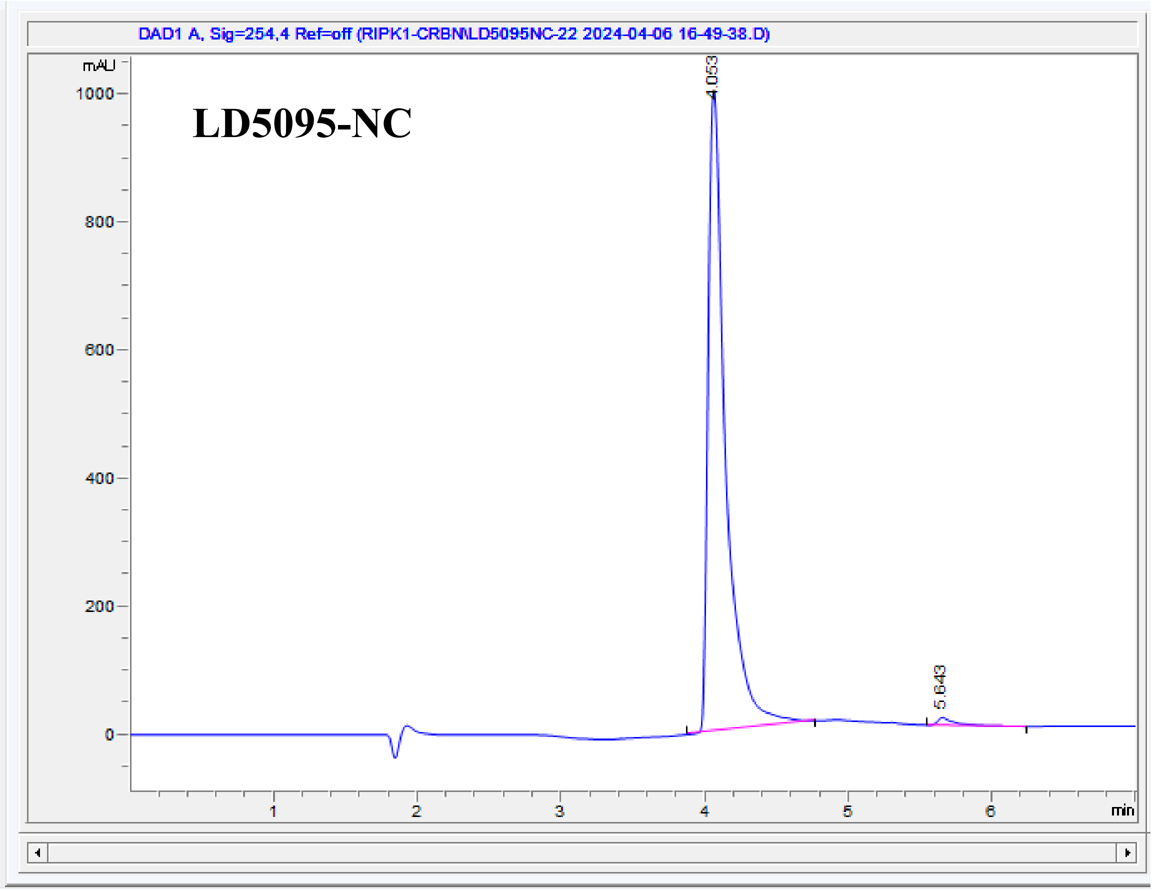


| # | Time | Area | Height | Width | Area% | Symmetry |
|---|---|---|---|---|---|---|
| 1 | 4.053 | 8515.9 | 1002.3 | 0.1251 | 98.842 | 0.423 |
| 2 | 5.643 | 99.8 | 12.3 | 0.1106 | 1.158 | 0.367 |
